# Bleeding in patients and mice with Alagille syndrome: sex differences and risk factors

**DOI:** 10.1101/2021.04.13.439679

**Authors:** Simona Hankeova, Noemi Van Hul, Jakub Laznovsky, Naomi Hensen, Elvira Verhoef, Tomas Zikmund, Feven Dawit, Michaela Kavkova, Jakub Salplachta, Liting Li, Marika Sjöqvist, Bengt R. Johansson, Mohamed Hassan, Vitezslav Bryja, Urban Lendahl, Andrew Jheon, Jian She Wang, Florian Alten, Kristina Teär Fahnehjelm, Björn Fischler, Jozef Kaiser, Emma R. Andersson

## Abstract

Alagille syndrome (ALGS) is generally thought of as a pediatric cholestatic liver disease, but spontaneous bleeds are a major cause of death. Here, we investigated bleeding in patients and a mouse model for ALGS (*Jag1*^*Ndr/Ndr*^ mice) and asked whether phenotypes identified in mice could be detected in patients non-invasively. *Jag1*^*Ndr/Ndr*^ mice spontaneously bled, exhibiting a thin skull and vascular defects. Vascular abnormalities included artery-vein crossings, tortuous vessels, and capillary breakdown. Furthermore, *Jag1*^*Ndr/Ndr*^ arteries had sparse vascular smooth muscle cell coverage, which was further aggravated by hypertension. Retinographs from patients with ALGS, biliary atresia, or CADASIL confirmed tortuous blood vessels and artery-vein crossings in ALGS. We also identified sex-specific differences: fewer major vessels in female *Jag1*^*Ndr/Ndr*^ mice, and four-fold more female than male patients with intracranial hemorrhage. In conclusion, dysfunctional *Jag1* disrupted vascular growth and homeostasis, with sex-specific differences. The mouse model for Alagille syndrome demonstrated multiple bleeding risk factors and vascular defects that can be identified in patients non-invasively.

## Introduction

Alagille syndrome (ALGS) is characterized by liver and heart defects, as well as vertebral abnormalities, characteristic faces, and posterior embryotoxon (a white line visible through a clear cornea caused by a thickened Schwalbe’s ring) ^1–3^. In addition to these hallmark features, 9% manifest non-cardiac vascular features such as vascular stenosis, Moyamoya syndrome, or absence of specific vessels, and up to 15% experience spontaneous intracranial hemorrhage ^3–6^. Vascular defects are also a significant cause of transplant-associated complications ^6^. Bleeding in patients with ALGS is a major cause of death in this group, in which up to 25% of deaths can be attributed to intracranial hemorrhage ^3,6,7^. Bleeding may occur spontaneously, or upon minor head trauma, and is not correlated with liver-disease-induced coagulopathy ^8^. In cases of minor head trauma leading to hemorrhage, cranial bone has been described as inordinately thin ^4,9^, sometimes associated with “thin-walled arteries” ^4,9,10^.

ALGS is a genetic disorder caused by mutations in the Notch ligand *JAGGED1* or in rare cases in *NOTCH2* receptor (ALGS1; Online Mendelian Inheritance in Man/OMIM no. 118450, and ALGS2, OMIM no. 610205) ^11–13^. Notch signaling is a major blood vessel regulator, and Notch1/3/4 receptors, Delta like (Dll) 1/4 and Jagged1 (Jag1) ligands have key roles in endothelial cell (EC) growth, proliferation, arterial specification, and mural cell recruitment ^14–16^. Appropriate angiogenesis and vascular homeostasis are essential for organism health. Notch pathway mutations are the cause of at least two congenital diseases with vascular involvement: Cerebral Autosomal Dominant Arteriopathy with Subcortical Infarcts and Leukoencephalopathy (CADASIL, mutations in *NOTCH3*) ^17^ and Adams Oliver syndrome (mutations in *NOTCH1*, *DLL4* or *RBPJk* ^18–21^, for review see ^22^). The tight association of Notch mutations with congenital vascular diseases underscores the need for appropriate animal models of Notch-mutant development and disease, in which to test potential therapies.

We recently developed a mouse model for ALGS, *Jag1*^*Ndr/Ndr*^ mice, which recapitulates the hallmark hepatic and cardiac features of ALGS ^23^. In this report, we explored whether *Jag1*^*Ndr/Ndr*^ mice provided insights into cellular mechanisms contributing to vascular compromise and uncovered potential risk factors for vascular events in patients with ALGS.

Despite cholestatic liver disease, *Jag1*^*Ndr/Ndr*^ mice did not display coagulopathy, as assessed by a Thrombin Anti-Thrombin (TAT) assay. Instead, bleeding in *Jag1*^*Ndr/Ndr*^ mice could be attributed to a combination of skull thinning, female sex, abnormal vascular development and vascular compromise. Our data recapitulated previous reports of the role of *Jag1* in vascular development, including reduced endothelial vascular outgrowth, decreased branching and fewer tip cells. Novel phenotypes, specific to *Jag1*^*Ndr/Ndr*^ mice, were a thinner skull, and vascular defects such as aberrant arteriovenous crossings, tortuous vessels and degenerating retinal intermediate plexus capillaries. *Jag1*^*Ndr/Ndr*^ arterioles exhibited poor vascular smooth muscle cell coverage (VSMC), which was exacerbated with age or upon an increase in blood pressure, resulting in thin-walled arteries and occasionally an aneurysm. Although sparse VSMC coverage was reminiscent of CADASIL, blood vessels in patients with ALGS showed significant defects in vessel numbers and venous tortuosity, while patients with CADASIL did not show defects in these analyses. *Jag1*^*Ndr/Ndr*^ mice are thus a useful model to study ALGS-related skull and vascular defects and could be used to test potential treatments. Importantly, vascular pathology could be quantified non-invasively in retinographs from patients with ALGS, suggesting fundus photography should be further investigated as a predictive biomarker for vascular health and bleeding in this pediatric population.

## Results

### Patients with Alagille syndrome exhibit sporadic spontaneous and provoked hemorrhages, with sex-differences

Previous studies have shown that vascular defects are common in ALGS, but have not identified any sex differences in disease presentation ^6,10,24^. First, we asked which vascular events were most often identified in patients, and whether vascular events were affected by sex. To test whether sex impacted on bleeding events, we retrospectively analyzed a cohort of 156 patients with ALGS from the Children’s Hospital of Fudan University. In this cohort, the frequency of intracranial bleeds was 3.21% (3 males of 94 males, and 2 females of 62 females). There were no overt sex differences in this patient cohort, though the low frequency of bleeds, and low numbers of patients with bleeds, limits comparisons. The most common type of bleeding events were intracranial bleeding and upper gastrointestinal bleeding (Table 1). In order to methodically map vascular defects in a larger group of patients with vascular events, we performed a systematic literature search, and identified 150 patients with ALGS and a vascular structural abnormality (e.g. stenosis, aneurysm or occlusion) or with a spontaneous/provoked bleed or ischemic event (Figure 1A – C, Supp Table 1, for search terms and strategy see Material and methods). The most commonly reported structural abnormalities were stenosis, aneurysm and Moyamoya (Figure 1B). Stenosis (61/150, 41%)^10,24–59^ (Figure 1B), was equally prevalent among males and females. Forty-five patients (45/150, 29.3%) ^25,26,28,29,31–34,37,38,44,45,47–49,52,53,56,59,60^ were reported with abdominal region vessel stenosis and twenty-nine patients (29/150, 19%) ^10,24,26,28,35,40,41,43,44,47,48,50,54,57–59^ were reported with head and neck arterial stenosis, of which six patients had both (6/150, 4%) ^26,28,44,47,48,59^. The predominant form of stenosis (apart from pulmonary artery stenosis) was renal artery stenosis (23/150, 15.3%) ^26,28,29,31–33,37,38,44,45,49,52,53,56,59–61^, followed by aortic stenosis or coarctation (22/150, 14.7%) ^25,27,29–36,38,39,42,44,46,47,51,53,55,59,62,63^, internal carotid stenosis (16/150, 10.7%) ^10,24,26,35,40,41,43,50,54,58^ and celiac artery stenosis (11/150, 7.3%) ^25,26,32,34,47–49^. The second most commonly reported structural abnormality was an aneurysm (26/150 patients, 17.3%) (Figure 1B), mostly in the basilary (8/150 patients, 5.3%) ^24,32,37,39,58,64^, abdominal (8/150 patients, 5.3%) ^31,32,39,49,60,65^ and carotid (7/150 patients, 4.7%) ^10,24,39,48,66,67^ arteries. Vessel agenesis and hypoplasia (10 males vs 6 females, 1.7x more) ^24,25,28,29,33,38,41,42,58,68–71^ and esophageal varices were most reported in male patients (6 males vs 3 females, 2x more) ^26,30,67,72–77^ and patent ductus arteriosus was more highly reported in female patients (7 females vs 2 males, 3.5x more) ^37,38,42,45,63,78–81^ (Figure 1B). Esophageal varices develop secondary to portal hypertension and can lead to gastrointestinal bleeding ^82^. The most frequently reported functional events were bleeding events (55/150, 36.7%) and intracranial infarctions (12/150, 8%) ^10,24,41,48,83,84^ (Figure 1C). Among the bleeding events, the predominantly reported type was intracranial bleeding (25/150, 16.7%) ^24,38,39,51,64,65,67,79,85–91^, followed by pulmonary hemorrhage (15/150, 10%) ^45,92^. Other bleeding events included gastrointestinal (10/150, 6.7%) ^9,26,73,74,76,77,93–96^, nasal (2/150, 1.3%) ^61,73^ intrathoracic ^97^, intraperitoneal ^98^, renal ^60^, and ocular ^89^ bleeding, as well as hematochezia ^73^, hematuria ^48^ and hematemesis ^61^ (each 1/150, 0.7%) (Figure 1D). Intracranial bleeds included hematoma (generally induced by a head trauma due to thin skull), and hemorrhages (Figure 1E). Hematoma was reported equally frequently for males and females ^9,24,39,64,85,86,88,91^, whereas more females were reported with intracranial hemorrhages (12 females vs 3 males, 4x more, Figure 1E) ^24,38,39,51,64,65,67,79,85–90^.

**Table 1.**
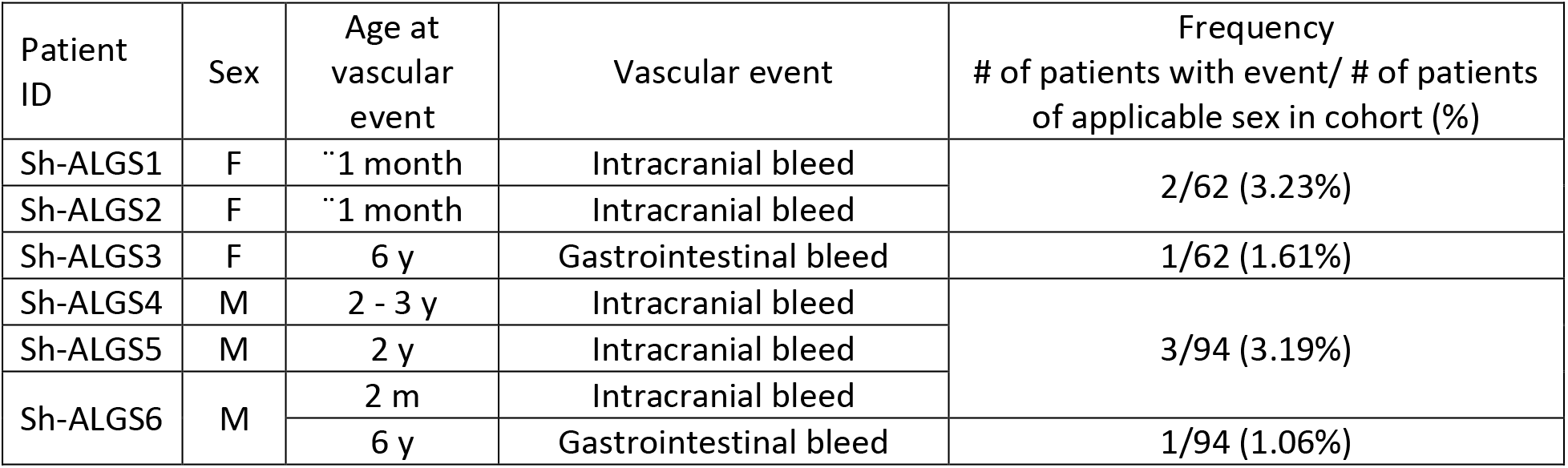
Vascular bleeds in a cohort of 156 patients from Children’s Hospital of Fudan University. F, female; M, male; y, year;

**Figure 1.**
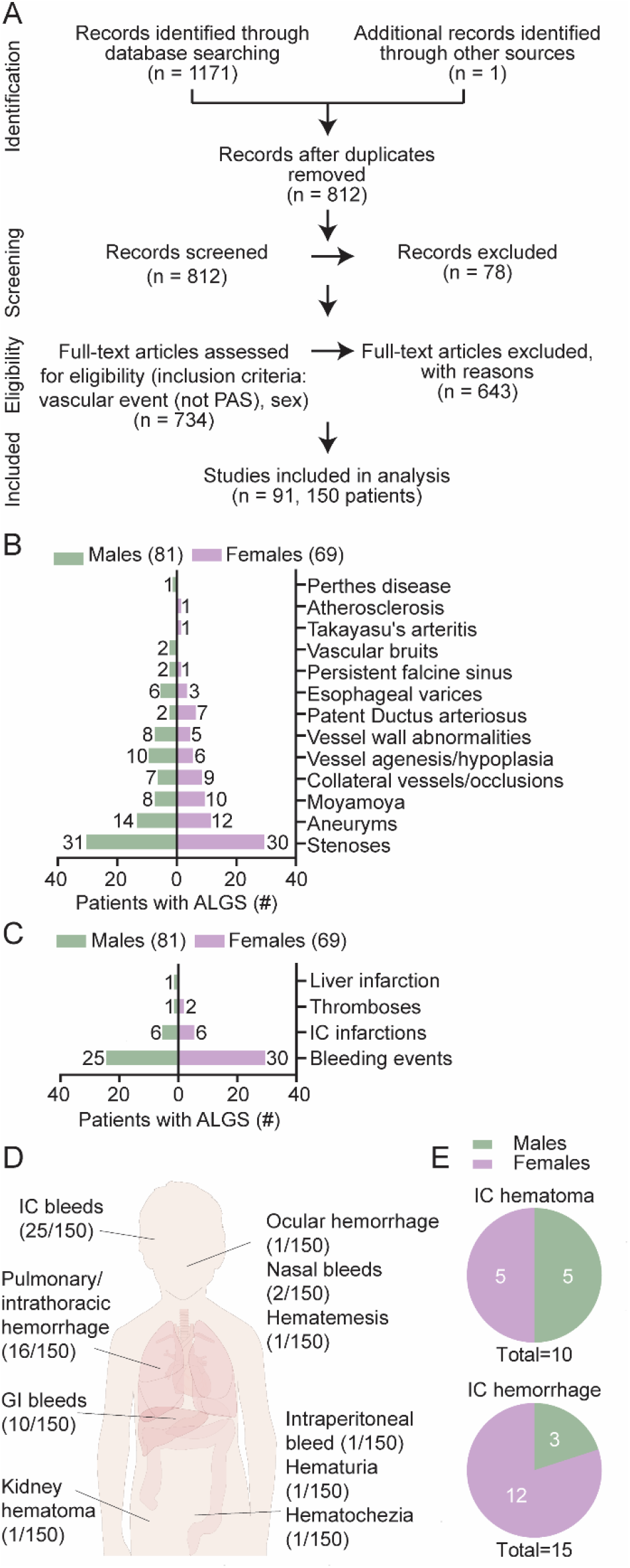
Patients with Alagille syndrome exhibit sporadic spontaneous and provoked hemorrhages, with sex-differences. **(A)** Systematic review search strategy that identified 91 relevant publications describing 150 individuals with Alagille syndrome (ALGS) and vascular events (excluding pulmonary artery stenosis) including sex data. **(B)** Structural vascular abnormalities grouped by prevalence in male or female patients with ALGS. **(C)** Functional vascular abnormalities grouped by prevalence in male or female patients with ALGS. The numbers next to the graph bars in B and C depict the number of individuals reported with a given anomaly per sex. Some patients had multiple vascular events and are included in several categories. For references and full details, see also Supplementary Table 1. **(D)** Fifty-five patients were reported with 59 bleeding events, most often intracranial (IC) bleeds. **(E)** Intracranial bleeds (25 patients) included 10 patients with hematoma (top chart) and 15 with hemorrhage (bottom chart). Pie graphs depict male/female distribution for hematoma and hemorrhage, with the number of reported patients as an inset. ALGS, Alagille syndrome; GI, gastrointenstinal; IC, intracranial; PAS, pulmonary artery stenosis.

Our data confirm previous reports that vascular events, including intracranial hemorrhage, are present in pediatric patients, and suggest that sex may impact on the likelihood of specific types of vascular events. Importantly, females with ALGS are reported four times more frequently with intracranial hemorrhages compared to males with ALGS.

### Cholestatic Jag1^Ndr/Ndr^ mice exhibit normal coagulation but a thinner skull and sporadic spontaneous and provoked hemorrhages

To investigate mechanisms contributing to bleeds, and whether the recently reported *Jag1*^*Ndr/Ndr*^ mouse model for ALGS mimicked risk factors reported in patients, we next explored known factors associated with bleeding events, including coagulopathy and fragile intracranial bones. We assessed coagulation activity by measuring the levels of Thrombin-Antithrombin (TAT) complexes in blood plasma. TAT levels are usually elevated in cholestasis in both mice and humans indicating liver disease-induced disrupted coagulation function ^99,100^. At postnatal day 10 (P10), all *Jag1*^*Ndr/Ndr*^ mice were jaundiced and cholestatic (Figure 2A and ^23^), but there were no consistent differences in coagulation function at this stage (Figure 2B). Next we investigated the skull of adult *Jag1*^*Ndr/Ndr*^ mice using micro computed tomography (μCT), since thin cranial bone was reported in patients with ALGS with intracranial bleeds ^4,9^. A qualitative analysis of the *Jag1*^*Ndr/Ndr*^ skulls revealed craniosynostosis in all *Jag1*^*Ndr/Ndr*^ mice examined (n=3) (Figure 2C, green arrowhead), which was previously described in patients with ALGS ^3,24,101^. We evaluated the thickness of the cranial bones, focusing on the full thickness (Figure 2C middle panels) and the compact bone (Figure 2C right panels), and found that *Jag1*^*Ndr/Ndr*^ cranial bones were thinner. The parietal bone of *Jag1*^*Ndr/Ndr*^ mice was thinner compared to the wild type mice, and often consisted of a single layer of compact bone, rather than two layers of compact bone encasing the spongy bone and trabeculae. This is clear when comparing, in wild type mice, the solid color in the middle panel (full thickness width, with corresponding cross-section at right) to the speckled pattern in the right panel (shortest distance width = top layer of compact bone separated from bottom layer by trabeculae, notable in corresponding cross-section at right). In contrast, *Jag1*^*Ndr/Ndr*^ parietal bone had similar widths in the whole width and the shortest width/compact bone analyses. To quantitatively assess the changes in the cranial bone, we first measured the length of the skull from the occipital bone (back of the skull) to the nasal bone. *Jag1*^*Ndr/Ndr*^ pups are strikingly smaller than wild type animals ^23^ however, the *Jag1*^*Ndr/Ndr*^ adult skull lengths were similar in size to wild types and were, on average, 94.4% of the wild type skull lengths (Figure 2D, p=0.0652). However, the volume of the *Jag1*^*Ndr/Ndr*^ cranial bone full width (Figure 2E, ~ 80% of wild type volume) and the compact bone (Figure 2F, ~ 78% of wild type volume) were both significantly lower than wild type skull volumes. The greater decrease in the cranial bone volume compared to length (Figure 2D – F, 80% vs 94%), and the bone fusion seen in parietal bone (Figure 2C) suggest a thinner skull in *Jag1*^*Ndr/Ndr*^ mice.

**Figure 2.**
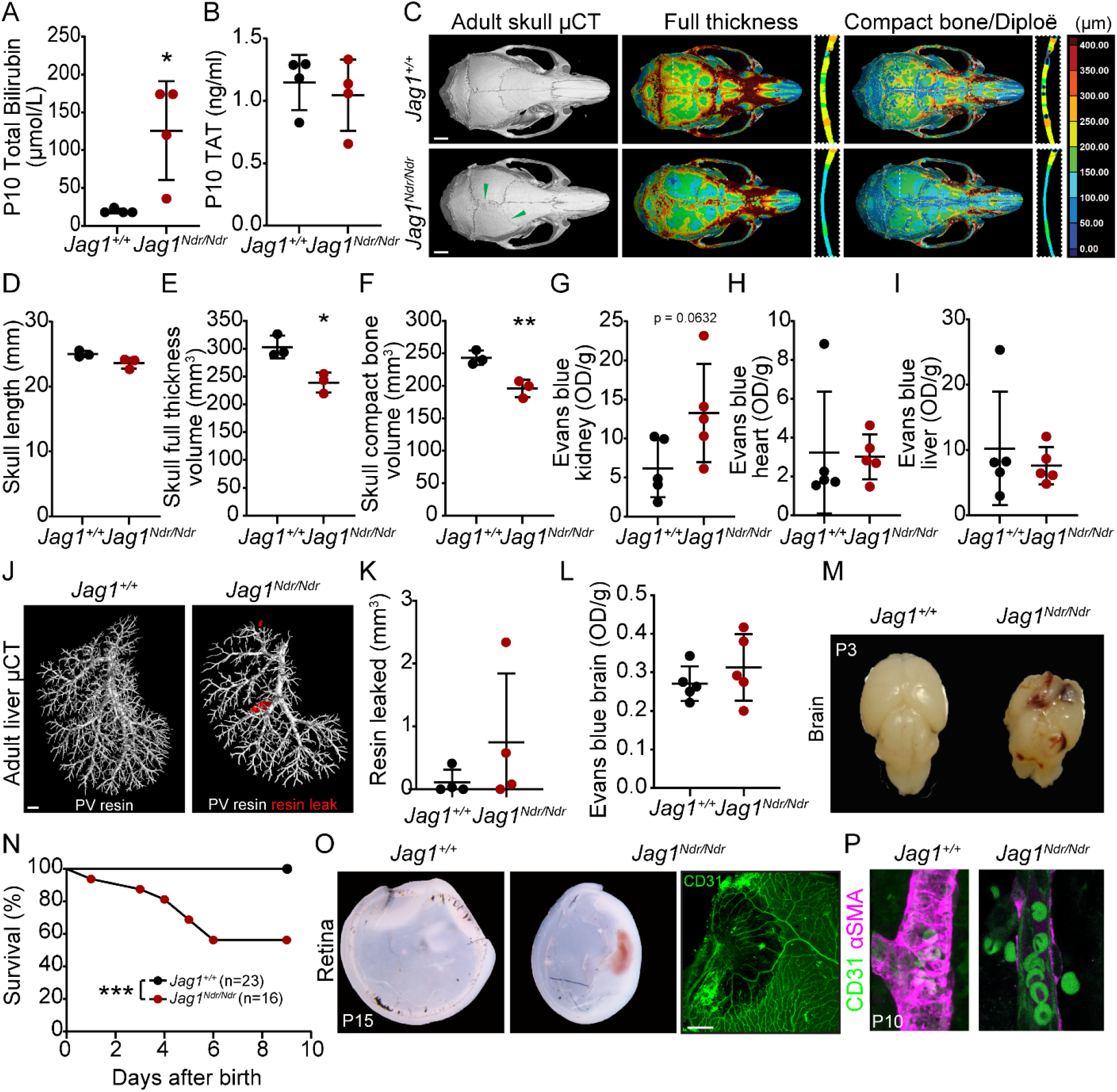
Cholestatic *Jag1*^*Ndr/Ndr*^ mice exhibit normal coagulation but a thinner skull and sporadic spontaneous and provoked hemorrhages. **(A)** Total bilirubin levels in plasma at P10. **(B)** Thrombin Antithrombin (TAT) enzyme-linked immunosorbent assay (ELISA) from plasma of postnatal day 10 (P10) mice. **(C)** Micro computed tomography (μCT) of adult skulls (left panel), the green arrowhead marks fused sutures in *Jag1*^*Ndr/Ndr*^ mice. Color map depicting wall thickness of periosteum. Dotted line highlights the area magnified in boxed region at right (middle panel). Color map depicting wall thickness of compact bone, including trabeculae (diploë). Dotted line highlights the area magnified in boxed region (right panel) in which diploë was excluded (black spaces in bone). Scale bar 2 mm. **(D)** Occipital to nasal bone skull length. **(E)** The total volume of intracranial periosteum (based on wall thickness analysis). **(F)** The total volume of intracranial compact bone (excluding diploë, based on wall thickness analysis). **(G)** Blood vessel permeability assessed by intravenous Evans blue assay. Evans blue was extracted from the kidney, **(H)** heart and **(I)** liver. **(J)** Resin injection into the portal vein followed by **(K)** quantification of resin leakage outside of the blood vessels (in red) in adult mice. Scale bar 1 mm. **(L)** Blood brain barrier permeability tested by Evans blue assay of the adult brain parenchyma. **(M)** Brain from P3 *Jag1*^*Ndr/Ndr*^ pup, which died due to brain hemorrhages (n = 2 of 53 *Jag1*^*Ndr/Ndr*^ mice). **(N)** Pup survival analysis between P0 and P10. Each dot represents the absolute percent of remaining animals in the group. **(O)** P15 *Jag1*^*Ndr/Ndr*^ mouse with spontaneous retinal bleeding. Scale bar 20 μm. **(P)** Red blood cells present outside the arterial vessel wall in P10 *Jag1*^*Ndr/Ndr*^ retina. Statistical significance was tested with unpaired Students t-tests (A, B, D – I, K, L) and survival analysis (N). Each dot represents quantifications from one animal in (A, B, D – I, K, L); graphs depict mean values ± standard deviation; p < 0.05 (*), p < 0.01 (**). μCT, micro computed tomography; OD, optical density; TAT, Thrombin Antithrombin; P, postnatal.

In order to test bleeding and blood vessel permeability in the kidney, heart and liver, we used an Evans Blue assay. There was a mild, non-statistically significant, increase in kidney vessel leakage in adult *Jag1*^*Ndr/Ndr*^ mice (p=0.0632) (Figure 2G), in which three of five *Jag1*^*Ndr/Ndr*^ mice displayed higher Evans blue values in kidney than the highest wild type values. Evans blue values were similar for wild type and *Jag1*^*Ndr/Ndr*^ mice in heart and liver (Figure 2H, I), though this interpretation should consider that one wild type also showed unusually high leakage of Evans blue. As a further measure of blood vessel resilience, we analyzed blood vessel leakage from synthetic resin injections into the hepatic portal vein ^102^. One of four *Jag1*^*Ndr/Ndr*^ mice (a female) had 21x more resin outside of the portal vein than control animals, with macroscopically obvious leakage (leaks pseudo colored red, Figure 2J, K). The three *Jag1*^*Ndr/Ndr*^ mice with no/less extravasated resin were males.

In order to explore central nervous system vascular accidents, we next examined the brain and the retina. Evans blue injection revealed that two adult *Jag1*^*Ndr/Ndr*^ mice (two females, the remaining three animals were males), of five tested, had slightly increased Evan blue leakage in the brain compared to the wild type animals, but on average there was no difference (Figure 2L). Fifty-three *Jag1*^*Ndr/Ndr*^ pups were monitored daily from birth until postnatal day 10 (P10), and in this group macroscopically obvious brain hemorrhages occurred in 2 *Jag1*^*Ndr/Ndr*^ pups (3.77%) but were not observed in wild type or heterozygous mice (Figure 2M). Survival data were collected for 16 of the *Jag1*^*Ndr/Ndr*^ pups, and in addition to one pup dying as a result of a macroscopic brain bleed (one of the 2/53 reported above), six pups were lost to analysis (deaths = 7/16, 43.8%, Figure 2N). These *Jag1*^*Ndr/Ndr*^ pups generally died between P1 – P6, and were cannibalized by the mothers, preventing analysis (Figure 2N). In surviving pups at P10 and P15, *Jag1*^*Ndr/Ndr*^ pups also sporadically displayed retinal hemorrhage (Figure 2O) or leaky retinal arterioles (Figure 2P).

In summary, our data show that a small percentage (ca 3-4%) of patients with ALGS and *Jag1*^*Ndr/Ndr*^ mice exhibit central nervous system bleeds. Taken together, *Jag1*^*Ndr/Ndr*^ mice recapitulate spontaneous and rare bleeds in different organs, in the absence of coagulation defects. Similar to the patients, skull thinning in *Jag1*^*Ndr/Ndr*^ mice may have contributed to nervous system bleeds.

### Delayed retinal vascular outgrowth and remodeling in Jag1^Ndr/Ndr^ mice

JAG1 is a proangiogenic ligand balancing the number of tip cells and stalk cells ^103^. We therefore investigated retinal sprouting angiogenesis in the mouse model for ALGS. The retina is part of the central nervous system and is vascularized postnatally in mice: during the first fifteen days after birth three capillary plexuses are established in the tissue (schematic Figure 3A, S – superficial, I – intermediate and D – deep capillary plexus)^104^. At P5, retinal blood vessels in *Jag1*^*Ndr/Ndr*^ mice were significantly delayed in outgrowth, a defect that was not symmetric between left and right eyes (Figure 3B, C). The delayed extension and growth persisted throughout angiogenesis, marked by delayed vertical sprouting to the deep capillary plexus at P10 (Supp Figure 1A), and reduced extension of the intermediate capillary plexus at P15 (Supp Figure 1B, green). The number of tip cells was significantly decreased at the vascular front at P5 (Figure 3D left panels, Figure 3E), while the remaining tip cells in *Jag1*^*Ndr/Ndr*^ mice displayed an increase in tips per tip cell, with filopodia extending in several directions per cell (Figure 3D blue arrowheads in right panels, Figure 3F). The overall number of filopodia per tip or tip cell was similar in wild type and *Jag1*^*Ndr/Ndr*^ mice (Supp Figure 1C).

**Figure 3.**
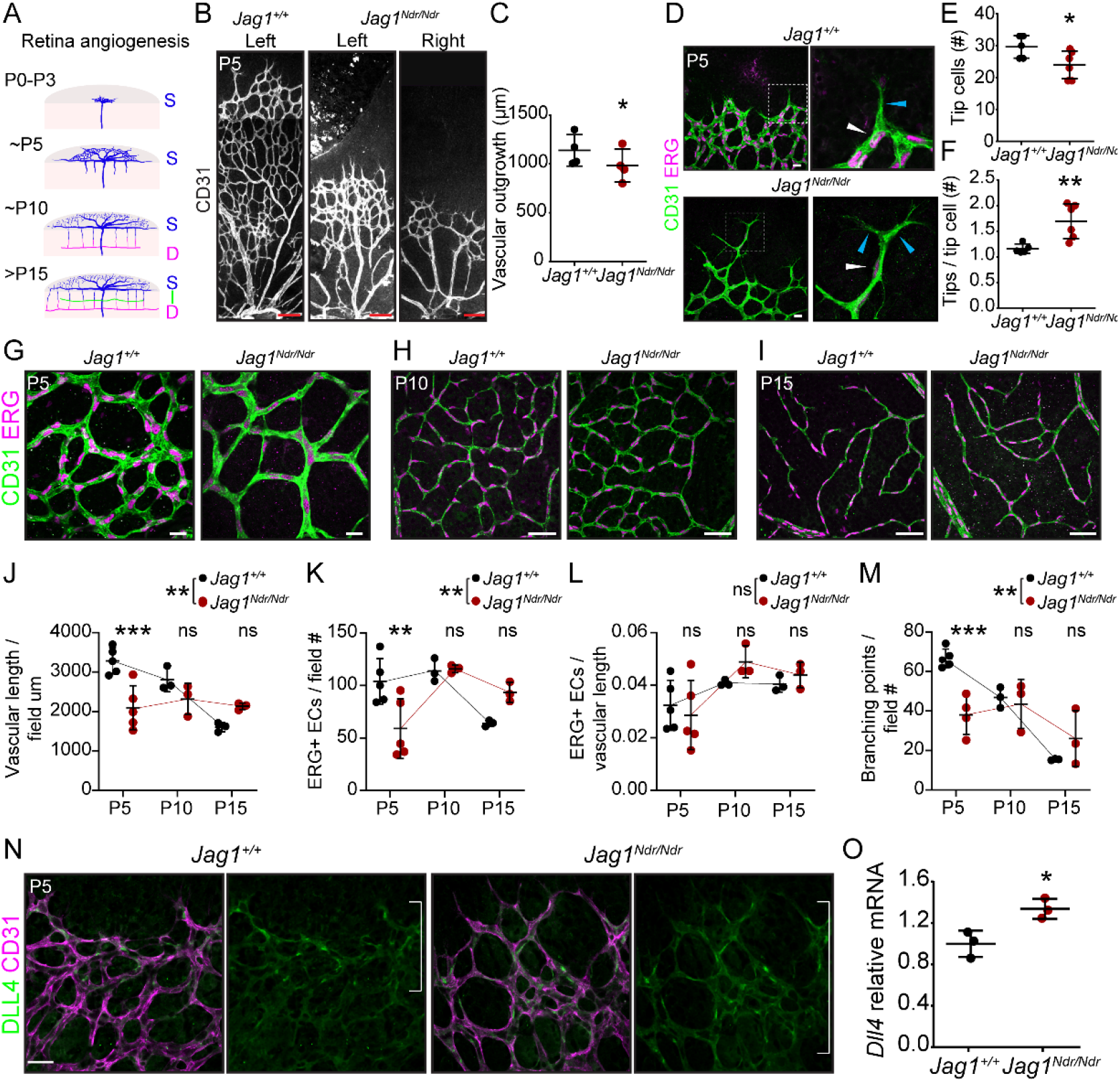
Delayed retinal vascular outgrowth and remodeling in *Jag1*^*Ndr/Ndr*^ mice. **(A)** Schematic depicting retinal angiogenesis between postnatal day 0 (P0) and P15. During the first 15 days after birth, three capillary plexuses are established S – superficial, I – intermediate, D – deep capillary plexus. **(A)** *Jag1*^*Ndr/Ndr*^ retinal vasculature stained for CD31 (Pecam1) was delayed in outgrowth, quantified in **(B)**. Scale bar 100 μm. **(C)** Immunofluorescence staining of vascular front at P5 with endothelial tip cells (boxed region). White arrowhead marks example of a tip cell, quantified in **(D)**. Blue arrowheads label tips (bundles of filopodia) of tip cells quantified in **(E)**. Scale bar 20 μm. **(F)** Retinal vasculature at postnatal day 5 (P5), **(G)** P10 and **(H)** P15. Scale bars 20 μm (F), 50 μm (G, H). **(I – L)** Quantification across retinal blood vessel development at P5, P10 and P15 of **(I)** vascular length per field, **(J)** number of endothelial cells (ERG+) per field, **(K)** number of endothelial cells (ERG+) per vascular length and **(L)** number of branching points per field. **(M)** Delta like 4 (DLL4) immunofluorescence in the vascular front at P5. Scale bar 20 μm. White brackets denotes area with high Dll4 intensity. **(N)** *Dll4* relative mRNA levels in whole retina lysates. Statistical significance was tested by unpaired Students t-test (B, D, E, N) or two-way ANOVA (I – L, difference between genotypes), followed by multiple comparison (I – L, differences at individual stages). Each dot represents quantification from one animal; graphs depict mean values ± standard deviation; p < 0.05 (*), p < 0.01 (**), p < 0.001 (***). D, deep capillary plexus; I, intermediate capillary plexus; P, postnatal; S, superficial capillary plexus.

We analyzed vascular plexus remodeling during angiogenesis at P5, P10 and P15 (Figure 3G – M), and total vascular length per field was significantly shorter at P5, but not at P10 or P15, in *Jag1*^*Ndr/Ndr*^ retinas (Figure 3G – J), indicating both delayed outgrowth and reduced remodeling. The total vascular length reflected differences in EC numbers, as evidenced by a significant decrease in ERG+ ECs at P5 in *Jag1*^*Ndr/Ndr*^ mice (Figure 3G – J, K). The number of ECs normalized to vascular length was similar between *Jag1*^*Ndr/Ndr*^ and wild type mice (Figure 3L), suggesting the delayed outgrowth (Figure 3B, C) is not due to migratory defects. The number of branching points was significantly lower at P5 in *Jag1*^*Ndr/Ndr*^ retinas but similar at P10 and P15 (Figure 3G – J, M). To determine whether the differences in blood vessel outgrowth were a consequence of altered proliferation, we quantified the number of phosphorylated histone 3 (PH3) and Pecam1 (CD31) double positive cells per area, in 200 μm long zones (in total 12 zones) from the optic nerve (Supp Figure 1D, scheme). The number of proliferating cells was significantly lower in *Jag1*^*Ndr/Ndr*^ mice at P5, and the bell-shaped proliferation curve peak was in Zone 4 (800 μm from the optic nerve) in *Jag1*^*Ndr/Ndr*^ mice, while the peak was in Zone 5 (1,000 μm from the optic nerve) in wild type mice (Supp Figure 1E). At P10 and P15 the number of proliferating ECs was lower in both *Jag1*^*Ndr/Ndr*^ and wild type retinas, and their distribution was similar (Supp Figure 1F, 1G). At P15, the *Jag1*^*Ndr/Ndr*^ vascular front had reached the retina periphery and the *Jag1*^*Ndr/Ndr*^ vascular length was similar to wild types (Supp Figure 1G and data not shown).

JAG1 competes with the ligand DLL4 for the NOTCH1 receptor ^103^ and loss of JAG1 leads to upregulation of DLL4 ^103,105^. Analysis of Dll4 expression revealed that DLL4 protein expression was highest in the vascular front and in tips cells in *Jag1*^*+/+*^ mice while it was more widespread in the *Jag1*^*Ndr/Ndr*^ vascular front (Figure 3N, white brackets) and *Dll4* mRNA levels in *Jag1*^*Ndr/Ndr*^ whole retina lysates were significantly increased (Figure 3O). Taken together, our results corroborated the view that JAG1 is necessary for correct angiogenesis, specifically for blood vessel outgrowth, branching, remodeling and proliferation, as reported previously for the EC-specific *Jag1* knockout ^103^. This may reflect a Notch gain of function phenotype via excess DLL4 signaling, as reported previously ^103^. Our analysis further revealed that, although *Jag1*^*Ndr/Ndr*^ tip cells are reduced in number, they form hyperbranched elongated tips. Finally, despite the delay in proliferation and outgrowth at P5, the *Jag1*^*Ndr/Ndr*^ superficial vasculature plexus was normalized by P15.

### Jag1^Ndr/Ndr^ mice display vascular guidance defects, with fewer and more tortuous blood vessels

Abnormal blood vessel growth or patterning can cause changes in blood flow leading to vessel occlusion ^106^, changes in shear stress ^107^ and changes to the vessel wall ^108^. We analyzed the presence of arteriovenous crossings (Figure 4A), which are considered pathological in mouse retina ^109,110^, and found a mean value of two arteriole/venule crossings per *Jag1*^*Ndr/Ndr*^ retina in both males and females (Figure 4A, B). *Jag1*^*Ndr/Ndr*^ females had significantly fewer arterioles (Figure 4C) and venules (Figure 4D) compared to female *Jag1*^*+/+*^ mice, and fewer venules compared to *Jag1*^*Ndr/Ndr*^ males (Figure 4D). Another pathological feature associated with changes in vessel walls, aneurysms, hypertension and aging, is vessel tortuosity ^108^. Arterial tortuosity was increased in two of four *Jag1*^*Ndr/Ndr*^ mice at P30 (Figure 4E) and in two of three *Jag1*^*Ndr/Ndr*^ mice at one year (Figure 4F), but the overall differences were not statistically significant. In contrast, overall venous tortuosity was significantly increased in *Jag1*^*Ndr/Ndr*^ mice independent of age (Figure 4G, 4H). Together, our data demonstrate patterning defects in major arterioles and venules of *Jag1*^*Ndr/Ndr*^ mice, with reduced numbers of vessels, and increased tortuosity, with sex-specific differences in the number of major arterioles and venules.

**Figure 4.**
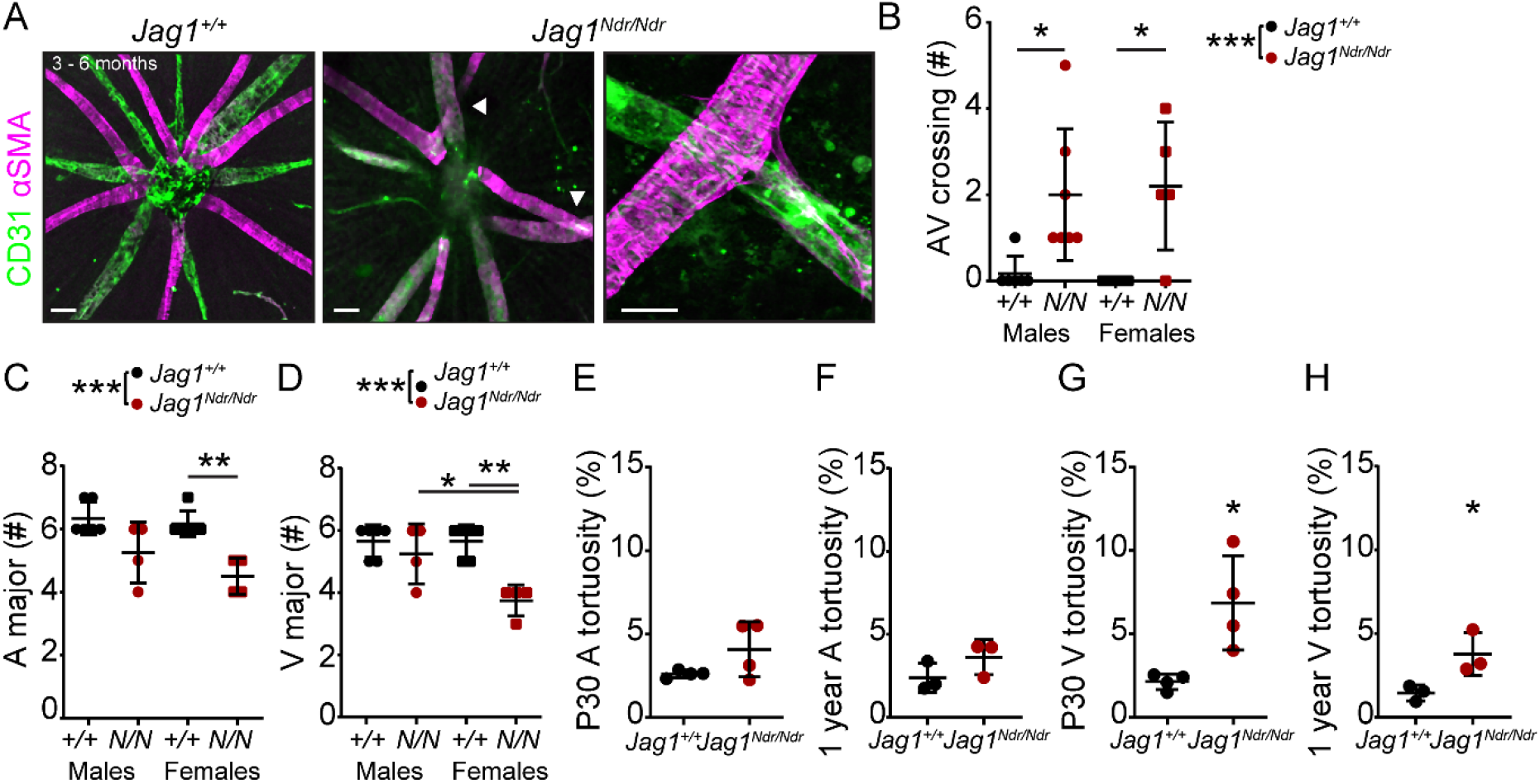
*Jag1*^*Ndr/Ndr*^ mice display vascular guidance defects, with fewer and more tortuous blood vessels. **(A)** Radial arrangement of arterioles (magenta) and venules (green) from the optic nerve. White arrowheads label arteriovenous crossings in *Jag1*^*Ndr/Ndr*^ retina. Scale bar left and middle panels 50 μm, and right panel 20 μm. **(B)** The number of arteriovenous crossings per retina grouped by genotype and sex. **(C, D)** Quantification of **(C)** major arterioles and **(D)** venules coming from the optic nerve head, grouped by genotype and sex. **(E, F)** Measurement of arterial tortuosity at **(E)** postnatal day 30 (P30) and at **(F)** 1 year. **(G, H)** Measurement of venous tortuosity at **(G)** P30 and at **(H)** 1 year. Statistical significance was tested by unpaired Students t-test (E – H) and two-way ANOVA followed by multiple comparison (B – D). Each dot represents quantifications from one animal; graphs depict mean value ± standard deviation; p < 0.05 (*), p < 0.01 (**). A, arteriole; AV, arteriovenous; αSMA, alpha smooth muscle actin; F, female; M, male; P, postnatal; V, venule.

### Jag1^Ndr/Ndr^ mice display sparse vascular smooth muscle cell coverage of arteries with large apoptotic gaps in old mice

Blood vessels are composed of two principal cell types: endothelial cells and mural cells, which include VSMCs and pericytes. The foremost Notch-related vascular disorder CADASIL (caused by mutations in *NOTCH3*) is characterized by poor arterial VSMC coverage ^85^ and in some cases is associated with neurovascular dementia ^112^. JAG1 also regulates arterial VSMC maturation and maintenance ^103,113,114^ and a ruptured aneurysm in a patient with ALGS suggested VSMC loss ^115^. We therefore investigated VSMCs and pericytes (another type of mural cell required for blood brain/retina barrier integrity ^116^) in *Jag1*^*Ndr/Ndr*^ mice and compared these to known CADASIL or *Notch3* mutant phenotypes.

In adult mice, CD13 positive pericytes were not reduced in *Jag1*^*Ndr/Ndr*^ retinas compared to *Jag1*^*+/+*^ retinas (Supp Figure 2A), comparable to previous reports for *Notch3*^*-/-*^ mice ^16^. We therefore focused on VSMC development and homeostasis. At P10, α smooth muscle cell actin (αSMA) positive VSMCs covered *Jag1*^*+/+*^ arterioles, with VSMCs oriented perpendicular to the blood vessel axis, while in *Jag1*^*Ndr/Ndr*^ retinas VSMC coverage was not as high and cells exhibited a more migratory morphology with their cellular axis parallel to the blood vessel axis (Figure 5A, boxed regions). Adult *Jag1*^*Ndr/Ndr*^ mice (at 3 – 6 months) displayed sparse VSMC arteriole coverage, with clear gaps between VSMCs and stenoses (green arrowheads) (Figure 5B). By one year of age, *Jag1*^*Ndr/Ndr*^ arteriolar VSMCs degenerated, resulting in αSMA-weak (Figure 5C blue arrowheads) or negative (Figure 5C white arrowheads) areas. αSMA negative areas were sometimes associated with aneurysms in *Jag1*^*Ndr/Ndr*^ arterioles (Figure 5C white arrows, also note the enlarged ECs). To determine whether the sparse VSMC coverage, identified using immunofluorescence, indeed reflected gaps between VSMCs we further analyzed VSMC morphology and arteriolar coverage using transmission electron microscopy of retinal and cardiac blood vessels. Retinal *Jag1*^*+/+*^ arteriolar VSMCs were present in a uniform layer surrounding the endothelial cells with wide VSMC-VSMC contacts (Figure 5D, VSMCs pseudocolored in magenta and ECs in green, white arrowheads refer to VSMC contacts). In contrast, retinal *Jag1*^*Ndr/Ndr*^ arteriolar VSMCs were sparse, thin, and not in contact with neighboring VSMCs (Figure 5D, white arrowheads show ends of VSMC not in contact with another VSMC). These data confirmed that narrow and widely spaced αSMA bands across arterioles (Figure 5B, C) reflected a thinner VSMC layer with fewer contacts. The VSMCs in *Jag1*^*Ndr/Ndr*^ coronary arteries displayed similar gaps (Supp Figure 2B, white arrowheads), suggesting this VSMC defect is not specific to the nervous system. The tight junctions between ECs were normal in *Jag1*^*Ndr/Ndr*^ mice (Figure 5D, Supp Figure 2B, green arrowheads). Arteriole diameter appeared greater in one-year-old *Jag1*^*Ndr/Ndr*^ retinas (Figure 5C), with thinning of VSMCs and ECs at 3 – 6 months (Figure 5B, D) suggesting a loss of a vascular tone. Although all *Jag1*^*Ndr/Ndr*^ mice displayed an increase in arterial diameter compared to wild types, this difference was not statistically significant (p = 0.0536), (Figure 5E). No change in venous diameter was detected in one-year-old *Jag1*^*Ndr/Ndr*^ retinas (Figure 5F). To investigate whether the changes in VSMCs could be due to altered JAG1 – NOTCH3 levels, we analyzed JAG1 and NOTCH3 protein expression on arterioles (Supp Figure 2C). JAG1 levels were significantly increased in *Jag1*^*Ndr/Ndr*^ ECs and VSMCs (Supp Figure 2C, D), in line with our previous report that dysfunctional JAG1^NDR^ accumulates in vivo ^117^. NOTCH3 protein levels were similar in *Jag1*^*+/+*^ and *Jag1*^*Ndr/Ndr*^ arterioles (Supp Figure 2C, E) and *Notch3* mRNA levels were similar in whole retina lysates (Supp Figure 2F).

**Figure 5.**
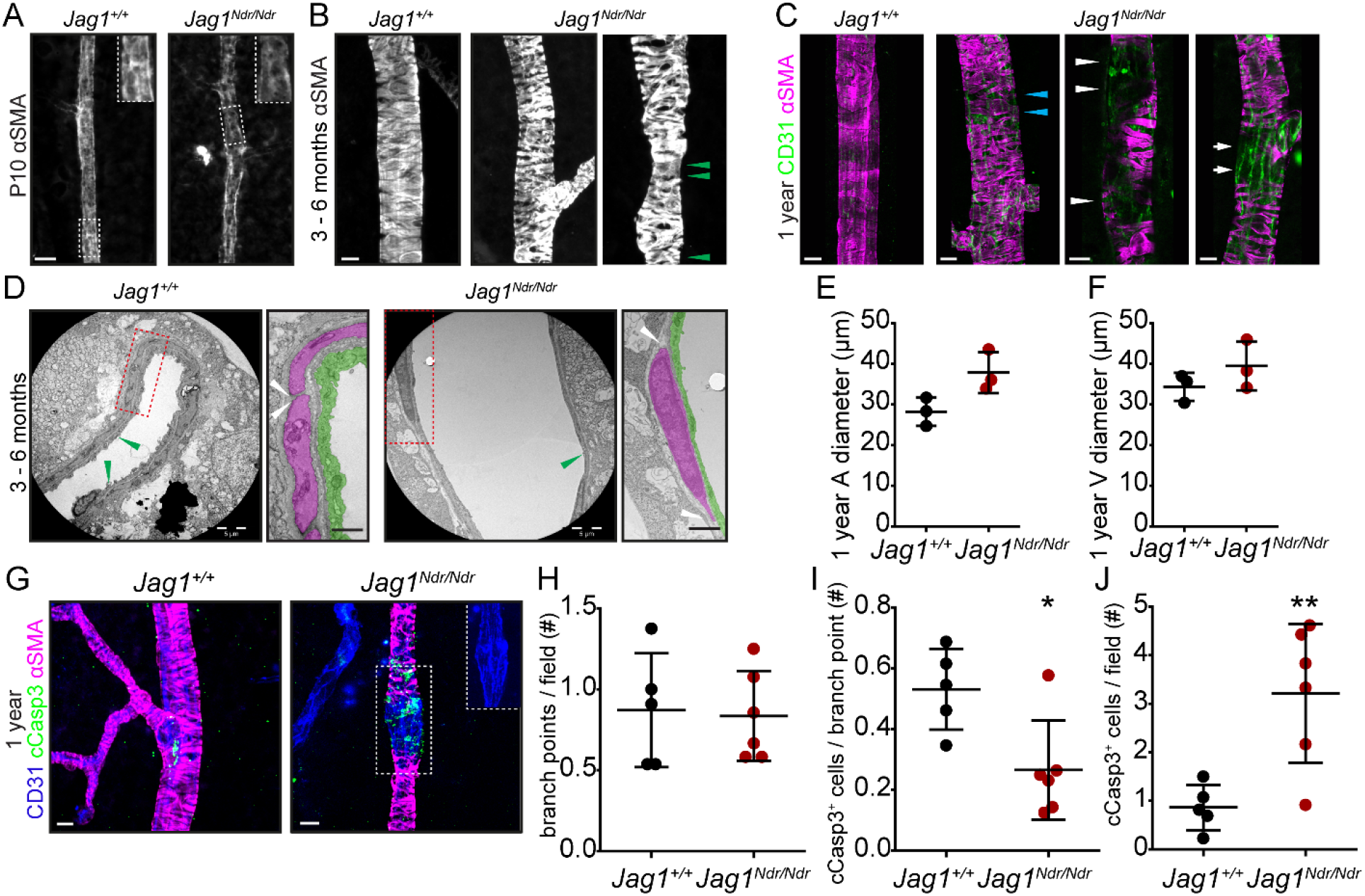
*Jag1*^*Ndr/Ndr*^ mice display sparse vascular smooth muscle cell coverage of arteries with large apoptotic gaps in old mice. **(A)** Alpha smooth muscle actin (αSMA) coverage of postnatal day 10 (P10) retinal arterioles, boxed regions highlight vascular smooth muscle cells orientation. In *Jag1*^*Ndr/Ndr*^ arterioles the VSMCs were oriented parallel to the blood vessel axis, rather than perpendicular and encircling the retinal. **(B)** Small gaps in αSMA VSMCS coverage of retinal arterioles in 3 – 6 months old mice. Green arrowhead labels focal arterial narrowing/stenosis. **(C)** αSMA coverage of one-year-old retinal arterioles. One-year-old *Jag1*^*Ndr/Ndr*^ mice arterioles display weaker αSMA coverage (blue arrowheads), αSMA negative gaps (white arrowheads) and an aneurysm (white arrows). Cropped images were placed on a black background for esthetic purposes. Scale bar (A – C) 10 μm. **(D)** Transmission electron microscopy of retinal arterioles. VSMCs color-coded in magenta, endothelial cells (EC) in green. Scale bar 5 μm, boxed region 1 μm. White arrowheads label the edges of VSMCs and the distance between vascular smooth muscle cells. Green arrowheads label EC tight junctions. **(E)** Diameter measurements of one-year-old retinal arterioles (p = 0.0536), and **(F)** venules (p=0.2690). **(G)** Cleaved caspase 3 (cCasp3) immunofluorescence staining of apoptotic cells in one-year-old retinal arterioles. Boxed region is a magnification of an aneurysm in a *Jag1*^*Ndr/Ndr*^ arteriole, note the distended endothelial cells. Scale bar 10 μm. **(H – J)** Quantifications of **(H)** branching points per field, **(I)** number of cCasp3 positive cells associated with a branching point and **(J)** number of cCasp3 positive cells per field, excluding those associated with branching point. Statistical significance was tested using unpaired Students t-test (E, F, H – J). Each dot represents one animal; graphs depict mean values ± standard deviation; p < 0.05 (*), p < 0.01 (**). A, arteriole; αSMA, alpha smooth muscle actin; cCasp3, cleaved caspase 3; P, postnatal; V, venule.

To determine whether the changes in VSMC coverage led to reactivity in the parenchyma we evaluated the distribution and morphology of astrocytes. The astrocytes surrounding *Jag1*^*Ndr/Ndr*^ arterioles were increased in density (Supp Figure 2G, 2I) and displayed numerous reactive bundles (Supp Figure 2G, white arrowheads). The astrocytes surrounding venules were similar in distribution in *Jag1*^*+/+*^ and *Jag1*^*Ndr/Ndr*^ retinas (Supp Figure 2H, J).

We next explored whether the αSMA negative regions in one-year-old mice were due to cells undergoing apoptosis. We quantified the number of cleaved Caspase 3 positive (cCasp3+) cells along the vessels in VSMCs, or associated with a branching point in ECs, since increased apoptosis was previously shown in VSMCs along the vascular length and in ECs at branch points in *Notch3* mutant mice ^16^. At this stage, the number of branch points was normal in *Jag1*^*Ndr/Ndr*^ arterioles (Figure 5G, H). Surprisingly, *Jag1*^*Ndr/Ndr*^ arterioles displayed fewer cCasp3+ ECs at branching points (Figure 5I) but significantly more cCasp3+ along the arteriole length, often associated with αSMA negative areas (Figure 5G, J). In sum, *Jag1*^*Ndr/Ndr*^ arterioles presented with sparse vascular VSMCs that died via apoptosis, sometimes associated with aneurysm formation. Our results thus demonstrate that JAG1 contributed to arteriole VSMC maturation and maintenance.

### Capillary homeostasis is compromised and retinal ganglion cells break down in Jag1^Ndr/Ndr^ retinas

Blood vessel homeostasis is crucial for the integrity of the blood retina/brain barrier and tissue health, and could be a factor contributing to bleeding in ALGS. A mature vascular plexus is characterized by vessel quiescence ^118^ and by the presence of mural cells, which were abrogated in *Jag1*^*Ndr/Ndr*^ mice (Figure 5 and Supp Figure 2). We therefore investigated adult retinal blood vessel homeostasis in *Jag1*^*Ndr/Ndr*^ mice. We studied all three retinal vascular plexuses in 3 – 6 month old mice, (Figure 6A), and while the superficial capillary plexus was equally branched in both *Jag1*^*+/+*^ and *Jag1*^*Ndr/Ndr*^ retinas (Figure 6A, B), the *Jag1*^*Ndr/Ndr*^ intermediate capillary plexus was less vascularized (Figure 6C) with significantly fewer branching points (Figure 6D). To understand how age impacted on vasculature in this model, we analyzed all three retinal vascular plexuses in one-year-old mice (Figure 6E). Both the superficial (Figure 6F) and the intermediate capillary plexuses were equally branched and vascularized (Figure 6G, H) in *Jag1*^*+/+*^ and *Jag1*^*Ndr/Ndr*^ retinas. The vascularization of the adult (3 – 6 months) *Jag1*^*Ndr/Ndr*^ retina (Figure 6B – D) was thus similar to the one-year-old *Jag1*^*+/+*^ retina (Figure 6F – H) with diminished branching and vascular density in the intermediate capillary plexus.

**Figure 6.**
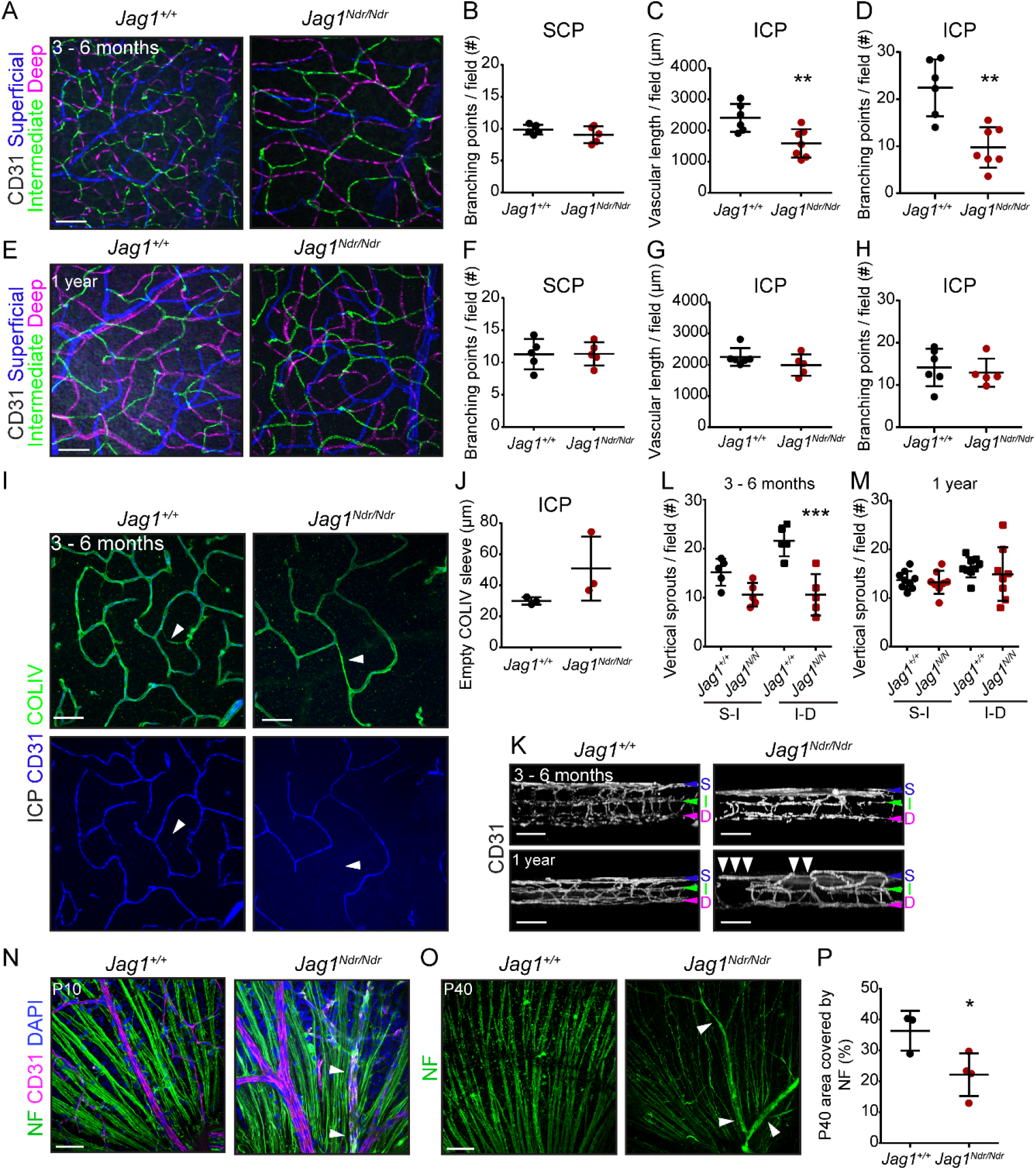
Capillary homeostasis is compromised, and retinal ganglion cells break down in *Jag1*^*Ndr/Ndr*^ retinas. **(A)** Pseudocolored classification of the three CD31+ capillary layers in adult retina (3 – 6 months old). Scale bar 50 μm. **(B)** Number of branching points in the superficial capillary plexus (SCP) of adult mice. **(C)** Vascular length per field in the intermediate capillary plexus (ICP) in adult mice. **(D)** Number of branching points per field in the ICP in adult mice. **(E)** Pseudocolored classification of the three CD31+ capillary layers in one-year-old retina. Scale bar 50 μm. **(F)** Number of branching points per field in SCP in one-year-old mice. **(G)** Quantification of vascular length per field in ICP in one-year-old mice. **(H)** Number of branching points per field in ICP in one-year-old mice. **(I)** Immunofluorescence staining of collagen IV (COLIV) and CD31 in ICP identifies regressed COLIV positive/CD31 negative capillaries (arrowheads). Scale bar 50 μm. **(J)** Quantification of empty collagen IV sleeve length per field in ICP. **(K)** Side view of all the retinal capillary plexuses in 3 – 6 months old (top panels) and one-year-old mice (bottom panels). The ICP in old *Jag1*^*Ndr/Ndr*^ mice was discontinuous, labeled by white arrowheads. The SCP is labeled with a blue arrowhead, the ICP with a green arrowhead and the DCP with a magenta arrowhead. **(L, M)** Number of vertical sprouts between SCP (S) and ICP (I) and ICP (I) and DCP (D) in **(L)** adult and **(M)** one-year-old mice. **(N, O)** Retinal ganglion cell axons stained with Neurofilament (NF) at **(N)** postnatal day 10 (P10) and **(O)** P40. Unspecific staining of blood is labelled with white arrowheads. **(P)** Quantification of area covered by NF in P40 mice. Statistical significance was tested using unpaired Students t-test (B – D, F – H, J, P) or two-way ANOVA followed by multiple comparisons (L, M). Each dot represents quantifications from one animal; Graphs depict mean values ± standard deviation; p < 0.05 (*), p < 0.01 (**). COLIV, collagen IV; D or DCP, deep capillary plexus; I or ICP, intermediate capillary plexus; NF, neurofilament; S or SCP, superficial capillary plexus; P, postnatal.

The vascularization of the intermediate capillary plexus at P15 in *Jag1*^*Ndr/Ndr*^ retinas was incomplete (Supp Figure 1B), hence it was not clear whether the lower vascular length and lower number of branching points in adult *Jag1*^*Ndr/Ndr*^ retinas reflected a persistence of a developmental defect, or whether capillaries had retracted. When a blood vessel retracts, it leaves behind its basal membrane containing Collagen IV (COLIV)^15^. In order to explore if the differences in the intermediate capillary plexus density were a result of capillary regression, we stained for COLIV (Figure 6I) and measured the lengths of empty COLIV sleeves. Empty COLIV sleeves were 1.7x longer in the *Jag1*^*Ndr/Ndr*^ intermediate capillary plexus (Figure 6J), but this difference was not statistically significant. We further quantified the number of vertical sprouts between the superficial and the intermediate (S – I) capillary plexuses and the intermediate and the deep (I – D) capillary plexuses in adult (Figure 6K, L) and one-year-old (Figure 6K, M) mice. The number of I – D vertical sprouts was significantly lower in 3 – 6 month *Jag1*^*Ndr/Ndr*^ retina compared to the *Jag1*^*+/+*^ retina (Figure 6L). In contrast, the number of vertical sprouts was similar between animals in one-year-old mice (Figure 6M). Nevertheless, vascular defects were present at both stages as noted by sparse vascular network in 3 – 6 months old *Jag1*^*Ndr/Ndr*^ retina (Figure 6K, top panel) and discontinued intermediate capillary plexus in the one-year-old *Jag1*^*Ndr/Ndr*^ retina (Figure 6K, bottom panel, white arrowheads).

The retina is a neural tissue, responsible for sight, in which retinal ganglion cells ^119^, the only afferent neurons, transmit signals from photoreceptors (rods and cones, the primary light-sensing cells), to the brain via the optic nerve. Approximately 90% of patients with Alagille syndrome display optic nerve anomalies such as optic disc drusen, which are hyaline-like calcified nodules ^120^. Retinal ganglion cells, whose cell bodies are located between the superficial and intermediate plexuses, are sensitive to hypoxia and to elevated levels of metabolites in the retina ^121,122^, suggesting retinal ganglion cells may be affected by the vascular defects identified here. To evaluate if the changes in retinal vasculature had neurological consequences, we investigated retinal ganglion cell axons by staining for neurofilament (NF). The retinal ganglion cells’ axons appeared healthy in both *Jag1*^*+/+*^ and *Jag1*^*Ndr/Ndr*^ retinas at P10 (Figure 6N, white arrowheads label non-specific non-NF immunostaining of blood, due to use of mouse-on-mouse antibody), suggesting overall development of retinal ganglion cells was not severely disturbed. However, at P40 the retinal ganglion cells axons in *Jag1*^*Ndr/Ndr*^ mice were thinner, tangled (Figure 6O) and covered 40% less area compared to *Jag1*^*+/+*^ mice (Figure 6P). In conclusion, our results show that the *Jag1*^*Ndr/Ndr*^ adult vasculature aged prematurely, marked by retracting retinal capillaries resulting in reduced and later discontinued vascular network. The onset of vascular degeneration was associated with retinal ganglion cell degradation in *Jag1*^*Ndr/Ndr*^ mice.

### High blood pressure compromises vascular smooth muscle cell homeostasis in Jag1^Ndr/Ndr^ mice

High blood pressure is one of the major risk factors contributing to cardiovascular diseases, subarachnoid hemorrhages and vessel tortuosity ^108,123^. Hypertension was reported in several patients with ALGS related to renal artery stenosis ^24,124^, or associated with visceral artery aneurysm ^49^, and may lead to complications during organ transplantation ^125^. *Jag1*^*Ndr/Ndr*^ mice had significantly lower mean blood pressure (Figure 7A), ruling out high blood pressure as a cause of abnormal VSMCs and vessel tortuosity. To investigate whether increased blood pressure impacted on bleeding or vascular health, we treated *Jag1*^*Ndr/Ndr*^ mice with Angiotensin II (AngII), a VSMC vasoconstrictor, for two weeks (experimental set up Figure 7B). Hypertension was defined as a systolic blood pressure above 140 mmHg ^126^ (Figure 7C, dotted line). Responsiveness to AngII was variable, we therefore selected AngII – treated animals with a minimum 20% increase in their mean blood pressure compared to baseline for further study (4 of 9 *Jag1*^*+/+*^ mice and 5 of 7 *Jag1*^*Ndr/Ndr*^ mice, Supp Figure 3A, B). Of note, *Jag1*^*+/+*^ mice tended to lose weight during AngII treatment, while *Jag1*^*Ndr/Ndr*^ mice increased in weight, resulting in a significant difference between these two groups (Supp Figure 3C, D).

**Figure 7.**
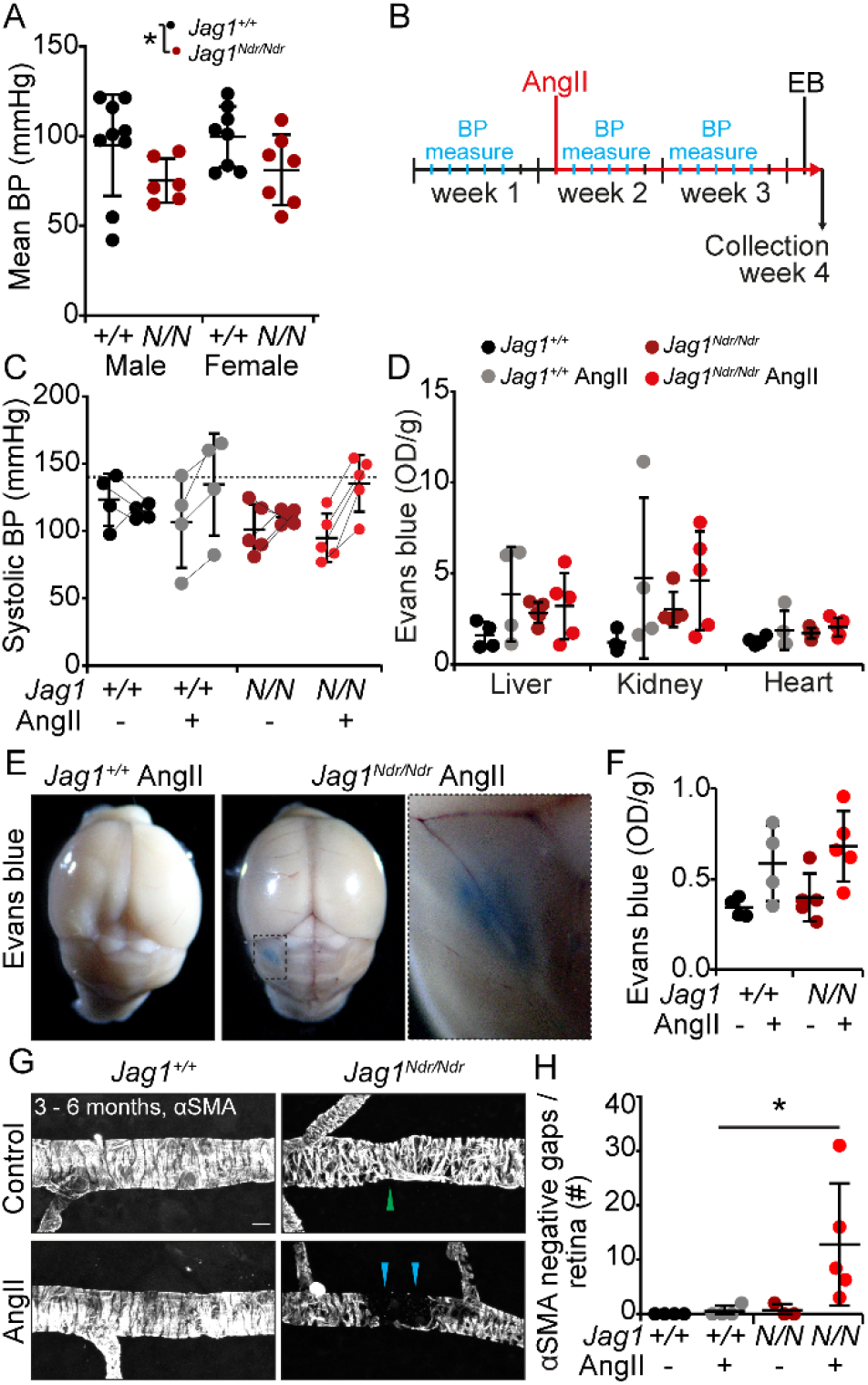
High blood pressure compromises vascular smooth muscle cell homeostasis in *Jag1*^*Ndr/Ndr*^ mice. **(A)** Mean blood pressure (BP) measurement in adult mice grouped by genotype and sex. **(B)** Experimental set up for Angiotensin II treatment. During week one, BP measurements were taken daily (blue tick marks) to establish the baseline levels and habituate the mice to measurements. At the beginning of week two, osmotic pumps with Angiotensin II or PBS control (red line) were implanted and BP was measured daily (blue tick marks). At the beginning of week four, mice were injected with Evans blue (EB) and organs were collected the next day. **(C)** Systolic BP levels before and during Angiotensin II treatment. The thin lines identify the same animal before and during treatment. The dotted line labels the cutoff for hypertension >140 mmHg systolic BP. These data are not tested for significance since animals were selected for analysis based on their response to Angiotensin, see also Supplementary Figure 3. **(D)** Measurement of Evans blue extracted from liver, kidney and heart. **(E)** Qualitative assessment of blood brain barrier permeability with Evans blue after Angiotensin II treatment, in which one *Jag1*^*Ndr/Ndr*^ mouse of 5 displayed a macroscopically obvious Evans blue leakage. **(F)** Measurement of Evans blue extracted from the brain parenchyma, not including the hemisphere with the major leakage in (E). **(G)** Alpha smooth muscle actin (αSMA) coverage of retinal arterioles treated with control (PBS) or Angiotensin II. The green arrowhead marks a focal arterial narrowing in a *Jag1*^*Ndr/Ndr*^ mouse. The blue arrowheads label an αSMA negative gap in an Angiotensin II-treated *Jag1*^*Ndr/Ndr*^ mouse. Scale bar 10 μm. **(H)** Quantification of αSMA negative gaps in arterioles per retina. Statistical significance was tested with two-way ANOVA followed by multiple comparisons (A, D, F, H). Each dot represents one animal; graphs depict mean values ± standard deviation; p < 0.05 (*). AngII, Angiotensin II; αSMA, alpha smooth muscle actin; BP, blood pressure; EB, Evans blue; OD, optical density.

To evaluate if the increased blood pressure affected blood vessel permeability, we injected the mice with Evans blue. The amount of Evans blue extracted from internal organs was similar in AngII-treated *Jag1*^*+/+*^ and *Jag1*^*Ndr/Ndr*^ mice and was similarly mildly increased compared to the untreated mice (Figure 7D). Macroscopic evaluation of the brain revealed that one of five AngII-treated *Jag1*^*Ndr/Ndr*^ mice (a male) displayed Evans blue leakage outside of the intracranial vessels, suggesting a localized bleed in this animal (Figure 7E). However, the amount of Evans blue extracted from the brain tissue (excluding the hemisphere with a bleed) was similar in AngII-treated *Jag1*^*+/+*^ and *Jag1*^*Ndr/Ndr*^ mice (Figure 7F). The amount of Evans blue extracted from brain tissue did not correlate with the change in blood pressure during AngII treatment for all animals tested (data not shown).

Finally, we examined whether increased blood pressure affected VSMCs. *Jag1*^*+/+*^ VSMCs were not affected by AngII treatment, by contrast arteriolar αSMA expressing VSMCs in *Jag1*^*Ndr/Ndr*^ retina were absent in large patches (Figure 7G, blue arrowheads), resulting in a significant increase in αSMA negative gaps (Figure 7H), reminiscent of *Notch3^−/−^* mice at P30 ^16,127^. In summary, our data show that VSMC homeostasis is compromised in AngII treated *Jag1*^*Ndr/Ndr*^ mice, leading to gaps in VSMC coverage, and to intracranial vessel leakage in one of five *Jag1*^*Ndr/Ndr*^ mice.

### Vascular defects identified in Jag1^Ndr/Ndr^ mice can be quantified non-invasively in patients with Alagille syndrome using retina fundus photography

Previous reports have shown that patients with ALGS have abnormal retinal vasculature ^128–131^, but the degree of tortuosity and patterning defects have not been quantified, nor compared to animal models, or CADASIL. We first analyzed retinal vasculature in a previously reported cohort of CADASIL patients ^132,133^. There were no significant differences in the numbers of arteriovenous crossings (Figure 8A, B), major arterioles or venules (Figure 8C, D) nor in arterial or venous tortuosity (Figure 8E, F) in patients with CADASIL compared to age-matched control patients. Arteriovenous crossings are more common in healthy human retina than in wild type mouse retina ^134^, therefore the observed artery-vein crossings in controls and CADASIL patients were expected. Venous tortuosity was mildly increased in some of the patients with CADASIL, but the degree of tortuosity did not correlate with sex (data not shown), nor with Fazekas score (a measure of white matter T2 hyperintense lesions used as a proxy for chronic small vessel ischemia, Supp Figure 4A). Interestingly venous tortuosity, but not arterial tortuosity (Supp Figure 4B, C), correlated positively with age in control patients (r = 0.6783, p= 0.0077, Supp Figure 4C), but not in patients with CADASIL (r = 0.1398, p=0.6337).

**Figure 8.**
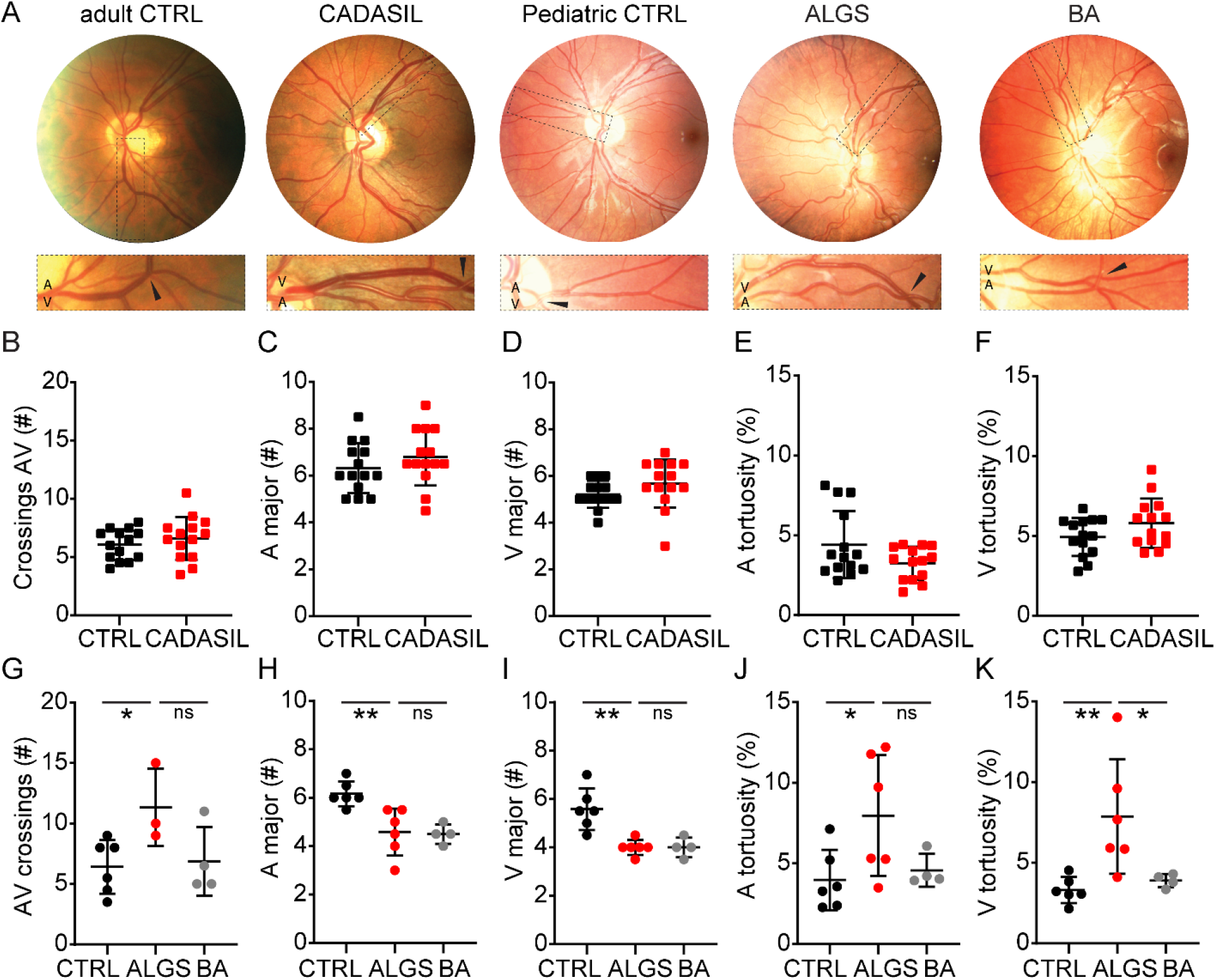
Vascular defects identified in *Jag1*^*Ndr/Ndr*^ mice can be quantified non-invasively in patients with Alagille syndrome using retina fundus photography. **(A)** Retinograph examples from a healthy adult individual, a patient with cerebral autosomal dominant arteriopathy with subcortical infarcts and leukoencephalopathy (CADASIL), a healthy pediatric individual, a patient with Alagille syndrome (ALGS) and a patient with biliary atresia (BA, as a cholestatic control). Boxed regions magnify areas with arteriovenous crossing marked by black arrowheads. **(B)** Number of arteriovenous crossings per retina, per person, in Controls and CADASIL patients. **(C, D)** Quantification of **(C)** major arterioles and **(D)** venules at the border of the optic disc per person in Controls and CADASIL patients. **(E, F)** Measurement of **(E)** arterial and **(F)** venous tortuosity in Controls and CADASIL patients. **(G)** Number of arteriovenous crossings per retina in pediatric Controls, patients with ALGS, and patients with BA. **(H, I)** Quantification of **(H)** major arterioles and **(I)** venules at the border of the optic disc in pediatric Controls, patients with ALGS, and patients with BA. **(J, K)** Measurement of **(J)** arterial and **(K)** venous tortuosity in pediatric Controls, patients with ALGS, and patients with BA. Statistical significance was tested by unpaired Students t-test (B – F) or by one-way ANOVA followed by multiple comparisons (G – K). Each dot represents one individual; graphs depict mean values ± standard deviation; p < 0.05 (*), p < 0.01 (**). A, arteriole; ALGS, Alagille syndrome; AV, arteriovenous; BA, biliary atresia; CADASIL, cerebral autosomal dominant arteriopathy with subcortical infarcts and leukoencephalopathy; CTRL, control; V, venule.

Finally, we analyzed retinal fundus photographs from pediatric patients with ALGS, biliary atresia (BA, as cholestatic controls), and age-matched healthy control patients (previously reported in ^128,135^) (Figure 8A). Similar to the *Jag1*^*Ndr/Ndr*^ mice (Figure 4A, B), arteriovenous crossings were significantly increased in patients with ALGS compared to the control group (Figure 8G). The number of major arterioles and venules were significantly lower in patients with ALGS compared to controls, but similar to patients with BA (Figure 8H, I), suggesting this phenotype (also present in *Jag1*^*Ndr/Ndr*^ mice in Figure 4C, D) may be related to cholestasis. Finally, we examined vascular tortuosity, which was consistently increased in *JagNdr/Ndr* veins (Figure 4G, H). Arterial tortuosity was significantly increased in patients with ALGS compared to controls (Figure 8J), and venous tortuosity was increased in patients with ALGS compared to both patients with BA and healthy controls (Figure 8K). Interestingly, most patients with ALGS had either very high venous or arterial tortuosity, but rarely both (Supp Figure 4D). As expected in this small cohort, no patients in this group experienced intracranial bleeds, or bleeding events linked to known vascular abnormalities, and we could therefore not test whether vascular bleeding events correlated with vascular defects. One patient with ALGS had severe bleeding due to coagulopathy (patient f) that was secondary to the cholestatic liver disease. Three patients with BA had experienced gastrointestinal bleeding (Table 2).

**Table 2.**
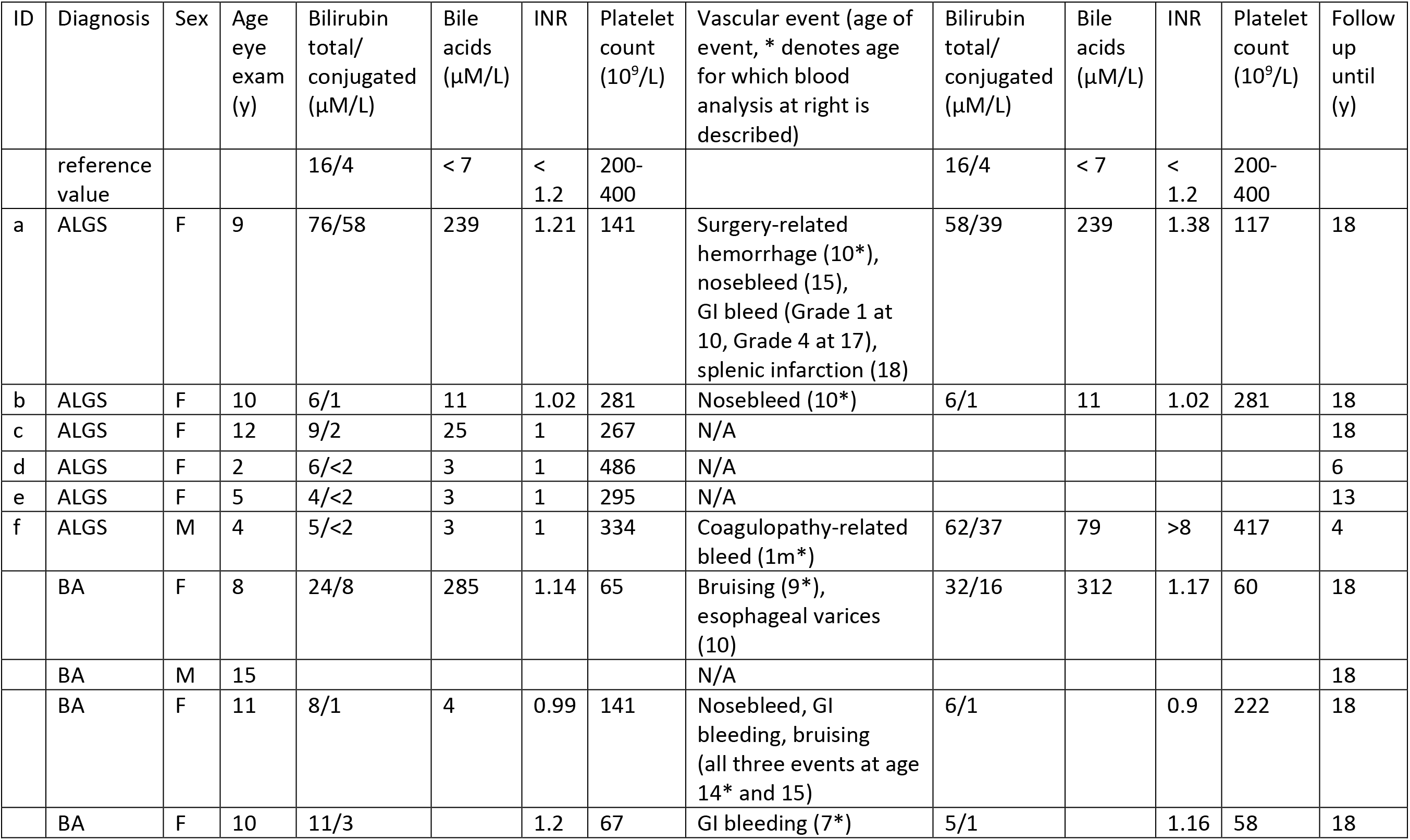
Vascular events in patients with Alagille syndrome and biliary atresia, with vascular anomalies quantified in the retinographs in Figure 8. (*) labels the age and vascular anomaly for which the blood values are provided. ALGS, Alagille syndrome; BA, biliary atresia; m, month.

| ID | Diagnosis | Sex | Age eye exam (y) | Bilirubin total/ conjugated ($\mu\text{M/L}$) | Bile acids ($\mu\text{M/L}$) | INR | Platelet count ($10^9/\text{L}$) | Vascular event (age of event, * denotes age for which blood analysis at right is described) | Bilirubin total/ conjugated ($\mu\text{M/L}$) | Bile acids ($\mu\text{M/L}$) | INR | Platelet count ($10^9/\text{L}$) | Follow up until (y) |
| --- | --- | --- | --- | --- | --- | --- | --- | --- | --- | --- | --- | --- | --- |
|  | reference value |  |  | 16/4 | < 7 | < 1.2 | 200-400 |  | 16/4 | < 7 | < 1.2 | 200-400 |  |
| a | ALGS | F | 9 | 76/58 | 239 | 1.21 | 141 | Surgery-related hemorrhage (10*), nosebleed (15), GI bleed (Grade 1 at 10, Grade 4 at 17), splenic infarction (18) | 58/39 | 239 | 1.38 | 117 | 18 |
| b | ALGS | F | 10 | 6/1 | 11 | 1.02 | 281 | Nosebleed (10*) | 6/1 | 11 | 1.02 | 281 | 18 |
| c | ALGS | F | 12 | 9/2 | 25 | 1 | 267 | N/A |  |  |  |  | 18 |
| d | ALGS | F | 2 | 6/<2 | 3 | 1 | 486 | N/A |  |  |  |  | 6 |
| e | ALGS | F | 5 | 4/<2 | 3 | 1 | 295 | N/A |  |  |  |  | 13 |
| f | ALGS | M | 4 | 5/<2 | 3 | 1 | 334 | Coagulopathy-related bleed (1m*) | 62/37 | 79 | >8 | 417 | 4 |
|  | BA | F | 8 | 24/8 | 285 | 1.14 | 65 | Bruising (9*), esophageal varices (10) | 32/16 | 312 | 1.17 | 60 | 18 |
|  | BA | M | 15 |  |  |  |  | N/A |  |  |  |  | 18 |
|  | BA | F | 11 | 8/1 | 4 | 0.99 | 141 | Nosebleed, GI bleeding, bruising (all three events at age 14* and 15) | 6/1 |  | 0.9 | 222 | 18 |
|  | BA | F | 10 | 11/3 |  | 1.2 | 67 | GI bleeding (7*) | 5/1 |  | 1.16 | 58 | 18 |

Altogether, our data demonstrated that patients with ALGS had vascular abnormalities that could be visualized and quantified non-invasively. Specifically, patients with ALGS have more arteriovenous crossings, fewer major arterioles and venules, and increased vascular tortuosity. Increased venous tortuosity is highly penetrant in patients with ALGS (identified in five of six patients). Importantly, *Jag1*^*Ndr/Ndr*^ mice recapitulate these phenotypes with fewer major blood vessels, more arteriovenous crossings, and increased vascular tortuosity.

## Discussion

In this study, we describe multiple roles for *Jag1* in patients and a mouse model for Alagille syndrome, in regulating blood vessel growth and homeostasis. We further identified risk factors that can lead to bleeds and demonstrated that vascular anomalies can be detected non-invasively in this pediatric populations using retinography. We investigated vascular anomalies in patients using a systematic literature search (150 patients with ALGS and vascular events), retrospective chart analysis (156 patients with ALGS), and retinography combined with retrospective chart analysis (6 patients with ALGS) and compared these findings to our investigation of vascular disease and bleeding in the ALGS mouse model. We showed that *Jag1*^*Ndr/Ndr*^ mice recapitulate spontaneous bleeds, and exhibit arteriovenous crossings, reduction in blood vessel numbers, and an increase in blood vessel tortuosity. Importantly, several identified vascular events or phenotypes demonstrated a sex-bias.

Our data suggest dysfunctional JAG1 compromises blood vessel development and homeostasis, with consequences for both endothelial and vascular smooth muscle cells. *Jag1*^*Ndr/Ndr*^ mice displayed many vascular developmental abnormalities such as delayed vascular outgrowth, reduced branching, abnormal tip cells and arteriovenous crossings. Previous studies indicate that many of the developmental defects observed here can be attributed to a role of JAG1 in endothelial cells ^103,105^. Developmental anomalies described in some patients with ALGS were agenesis of internal carotid artery ^24,71^ or aortic, renal and hepatic artery hypoplasia ^25,28,29,33,58^. Similarly, we and other groups have reported hypoplastic hepatic artery in *Jag1* mutant mice ^23,136^, though this defect is probably secondary to bile duct agenesis ^137^.

Notch signaling is crucial for arterial specification during embryogenesis and arterial smooth muscle cell maintenance throughout life ^138–141^. *Jag1*^*Ndr/Ndr*^ arterioles displayed delayed VSMC maturation and later poor VSMC coverage accompanied by accumulation of reactive astrocytes. The vast majority of vascular abnormalities in our literature review of patients with ALGS and vascular events, with sex data, were specific for arteries (196 events). Only twelve patients were reported with venous anomalies, which included cephalic vein stenosis ^48^, developmental venous anomaly, persistent falcine sinus (three patients) ^10^, abdominal wall venous dilation ^73^, displaced portal vein (three patients) ^69^, portal vein obstruction or stenosis ^63,94^ and arteriovenous malformations in liver and skin ^81^.

Retinal major arterioles and venules were tortuous in *Jag1*^*Ndr/Ndr*^ mice and patients with ALGS. Surprisingly, although most reported anomalies in ALGS are related to arterioles, venules were more consistently tortuous than arterioles in both patients and the mouse model. Retinal vessel tortuosity is associated with hypertension, diabetes, genetic disorders, hypoxia related retinopathies, cerebral vessel disease, stroke and ischemic heart disease ^142–145^. Tortuosity of vessels can also arise as a consequence of dolichoectasia, vessel wall weakening leading to dilation and elongation of blood vessels (usually weakening of the tunica intima/endothelial cells, but sometimes also tunica media/vascular smooth muscle cells). Dolichoectasia is fairly common in Alagille syndrome: vascular neuroimaging of a cohort of 19 patients identified neurovascular defects in six patients, of which four were dolichoectasia (4/19 = 21%) ^10^. Previous reports qualitatively described retinal vasculature from patients with ALGS as tortuous or with abnormally angulated vessels ^128,129,146^. In our study, five of six patients with ALGS (83%) had tortuous venules (tortuosity above 5%), while patients with BA and control patients were consistently under 5%, suggesting venous vessel tortuosity as a suitable additional diagnostic feature. Abnormal retinal vessels may be correlated with other abnormal vessels in the body ^147^, and non-invasive retinal vascular imaging may thus serve as a screening method to investigate vascular compromise in patients with ALGS. We suggest that this could be studied in larger patient cohorts, for example via GALA (Global Alagille Syndrome Alliance), to determine whether retinal vascular defects correlate with, or are predictive of, cardiovascular accidents. The retinal vascular abnormalities found in patients with ALGS were absent in retinographs from patients with CADASIL, with the exception of arteriovenous nicking, which was only present in patients with CADASIL retinas (data not shown). Vascular changes in CADASIL patients have been identified using optical coherence tomography and optical coherence tomography angiography ^132,133^. Future studies in patients with ALGS should include high-resolution techniques as they may provide other new clinical biomarkers of vascular disease severity and prognosis for ALGS.

Our systematic literature review of 150 patients with ALGS and vascular events revealed unexpected sex differences. Female patients with ALGS were more often reported with intracranial bleeds, specifically subarachnoid hemorrhage (SAH). SAH was reported four-fold more for females than males with ALGS, while in the general population females are 1.3 times more likely to suffer from SAH ^148^. While reporting bias may affect the relative frequency of reported minor symptoms such as hematuria versus a more serious complication such as SAH, it is unlikely that reporting bias would lead to more publications of females with SAH, in particular since bleeding events in general were balanced between males and females. In line with this data in humans, *Jag1*^*Ndr/Ndr*^ female mice exhibited more severely disrupted vasculature, with fewer major vessels. However, macroscopic bleeds were present in both male and female *Jag1*^*Ndr/Ndr*^ mice in this study.

Ten patients in the literature review were reported with epidural/subdural/intracranial hematoma, often associated with minor head trauma and thin temporal bones ^4,9^. The skull was also thin with fused compact bones in *Jag1*^*Ndr/Ndr*^ mice corroborating another risk factor in this mouse model. These data also further support previous reports for the role of Notch signaling in skeletal development and homeostasis ^149^.

Aging is a major risk factor contributing to the development of cardiovascular disease and is partly attributed to vessel wall changes leading to endothelial cell dysfunction ^150^ and loss of contractile VSMCs ^151^. Retinal vascular health was severely affected in adult (3 – 6 months old) *Jag1*^*Ndr/Ndr*^ mice as shown by decreased vascular density and sparse VSMC coverage. Specifically, the number of branching points in the intermediate capillary plexus and the number of vertical sprouts in the adult *Jag1*^*Ndr/Ndr*^ mice were similar to one-year-old wild type mice, demonstrating a premature aging-like phenotype. Reduced retinal vessel branching is also present in patients with ALGS ^129^ and could be seen in the retina fundus photographs from patients with ALGS (Figure 8), with reduced numbers of major arterioles and venules at the optic disc border. In the general population, ruptured intracranial aneurysms occur between 56 – 58 years of age ^152^, whereas the mean age for ruptured aneurysms in ALGS was 21.4 years ^24,39,51,64,65,67,87,88^. This may suggest that premature vascular aging in patients with ALGS leads to earlier onset of vascular bleeding and abnormalities. However, the presence of intracranial bleeding in newborns and infants (Table 1, Supplementary Table 1 ^85,90^) also suggest a developmental cause of bleeding.

The VSMC pathology we describe for *Jag1*^*Ndr/Ndr*^ mice (Figure 5, 7 and Supp Figure 2) is reminiscent of CADASIL, which may imply a risk for vascular dementia in patients with ALGS. Weak VSMC coverage in a ruptured intracranial aneurysm in patient with ALGS ^115^, further corroborates that VSMC pathology may be of concern in ALGS. VSMC health can be compromised by hypertension, indicating that patients’ blood pressure should be monitored carefully. Hypertension can lead to severe cardiovascular health complications like stroke, heart attack or SAH ^123^. Hypertension was reported in several patients with ALGS often related to renal artery stenosis ^24,38,124^ and associated with visceral artery aneurysm ^49^. We show that two weeks of hypertension in *Jag1*^*Ndr/Ndr*^ mice caused degeneration of VSMCs, forming αSMA-negative gaps in the VSMC layer, while the wild type VSMCs were unaffected. Hence, hypertension might pose as an even greater risk factor for patients with ALGS compared to the general population. Our study has not directly examined VSMCs in patients, therefore future studies should aim to systematically determine whether VSMCs are compromised in patients with ALGS, whether this interacts with elevated blood pressure (as our results in mice suggest), and how this is similar or dissimilar to CADASIL.

Patients with ALGS exhibit several ocular abnormalities, of which the most common optic disc anomaly is optic disc drusen ^130^, which compresses adjacent ganglion cell axons and promotes retrograde axonal degeneration ^153^. Our data suggest that vascular pathology in the JAG1-mutant retina leads to local tissue hypoxia, negatively influencing the surrounding neurons, especially the retinal ganglion cells, which might explain the high incidence of optic disc drusen in ALGS patients. Fortunately, even though ALGS patients display many ocular abnormalities, complete vision loss is very rare ^154^. However, monitoring retinal vasculature and the development of optic disc drusen could also be investigated as a prognostic marker for vascular anomalies in the rest of the body.

Our data demonstrate that multiple risk factors likely contribute to bleeding in patients with ALGS, and that these pathologies are possible to study in the *Jag1*^*Ndr/Ndr*^ mouse model (impact of sex, skull thinning and both EC and VSMC defects). Our data suggest that efforts should be made to identify and monitor patients that are at a greater risk of vascular disease (e.g. females with ALGS, aged 15 – 30 years old, who appear to have higher risk of SAH, or hypertensive patients with ALGS) and highlight the need for screening patients with ALGS for vascular and neurovascular changes. Future studies should aim to identify the patients at greatest risk of vascular disease in patients with ALGS, and devise treatments for vascular defects in this disease. *Jag1*^*Ndr/Ndr*^ mice present a suitable tool for pre-clinical trials of vascular intervention therapies, which could draw inspiration from the field of CADASIL, also aiming at treating VSMC compromise.

## Material and Methods

### Systematic review search strategy

The Medline (Ovid), Embase and Web of Science Core Collection database were used for publication search. After the initial search, the duplicate publications were removed. During the screening process records that were not written in English, or publications that were impossible to access were removed. Publications that were included in the systematic review contained information about the patient sex and vascular event except for pulmonary artery stenosis, which is a hallmark of Alagille syndrome.

Search strategy for Medline:

Field labels: exp/ = exploded MeSH term; / = non exploded MeSH term; .ti,ab,kf. = title, abstract and author keywords; adjx = within x words, regardless of order; * = truncation of word for alternate endings

1. Alagille Syndrome/
2. (alagill* adj3 (syndrome* or watson or disease)).ti,ab,kf.
3. (watson miller or arteriohepatic dysplasia or cardiovertebral syndrome or hepatic ductular hypoplasia or hepatofacioneurocardiovertebral or cholestasis with peripheral pulmonary stenosis or paucity of interlobular bile ducts or hepatic ductular hypoplasia).ti,ab,kf.
4. or/1-3
5. exp Cardiovascular System/ab, pa [Abnormalities, Pathology]
6. exp Vascular Diseases/
7. exp Hemorrhage/
8. exp Vascular Malformations/
9. (vascula* or cerebrovasc* or stroke* or aneurysm* or blood vessel* or artery or arteries or vein* or venous or moyamoya or moya moya or bleed* or hemorrhag*).ti,ab,kf.
10. or/5-9
11. 4 and 10
12. remove duplicates from 11

Search strategy for Embase:

Field labels: /exp = exploded Emtree term; /de = non exploded Emtree term; ti,ab = title and abstract; NEAR/x = within x words, regardless of order; * = truncation of word for alternate endings

((‘alagille syndrome’/de) OR ((alagill* NEAR/3 (syndrome* OR watson OR disease)):ti,ab,kw) OR (‘watson miller’:ti,ab,kw OR ‘arteriohepatic dysplasia’:ti,ab,kw OR ‘cardiovertebral syndrome’:ti,ab,kw OR hepatofacioneurocardiovertebral:ti,ab,kw OR ‘cholestasis with peripheral pulmonary stenosis’:ti,ab,kw OR ‘paucity of interlobular bile ducts’:ti,ab,kw OR ‘hepatic ductular hypoplasia’:ti,ab,kw))

AND

((‘cardiovascular system’/exp AND (abnormal* OR patholog*)) OR (‘vascular disease’/exp) OR (‘bleeding’/exp) OR (vascula*:ti,ab,kw OR cerebrovasc*:ti,ab,kw OR stroke*:ti,ab,kw OR aneurysm*:ti,ab,kw OR ‘blood vessel*’:ti,ab,kw OR artery:ti,ab,kw OR arteries:ti,ab,kw OR vein*:ti,ab,kw OR venous:ti,ab,kw OR moyamoya:ti,ab,kw OR ‘moya moya’:ti,ab,kw OR bleed*:ti,ab,kw OR hemorrhag*:ti,ab,kw))

Search strategy for Web of Science Core Collection:

Field labels: TS/Topic = title, abstract, author keywords and Keywords Plus; NEAR/x = within x words, regardless of order; * = truncation of word for alternate endings

TS=(alagill* NEAR/3 (syndrome* OR watson OR disease)) OR TS=(“watson miller” OR “arteriohepatic dysplasia” OR “cardiovertebral syndrome” OR hepatofacioneurocardiovertebral OR “cholestasis with peripheral pulmonary stenosis” OR “paucity of interlobular bile ducts” OR “hepatic ductular hypoplasia”)

AND

TS=(vascula* OR cerebrovasc* OR stroke* OR aneurysm* OR “blood vessel*” OR artery OR arteries OR vein* OR venous OR moyamoya OR “moya moya” OR bleed* OR hemorrhag*)

### Mouse maintenance and breeding

All animal experiments were performed in accordance with local rules and regulations and all animal experiments were approved by Stockholms Norra Djurförsöksetiska nämnd (Stockholm animal research ethics board, ethics approval numbers: N50/14, N61/16, N5253/19, N2987/20). The Nodder (*Jag1*^*+/Ndr*^ colony) mice were maintained in a mixed C3H/C57bl6 genetic background as reported previously ^23^. In brief, *Jag1+/Ndr* mice are maintained in a C3H background (EMMA C3HeB/FeJ-Jag1^Ndr^/Ieg, EM: 13207) and outbred to C57bl6 for experiments. *Jag1*^*Ndr/Ndr*^ mice are obtained from heterozygous crossings, and where possible littermates are used as controls. Because half of *Jag1*^*Ndr/Ndr*^ mice die within 10 days of birth (this study and^23^), with further deaths at adult stages (^23^), the analysis of 3 mice at age one year entails the generation of at least 15 *Jag1*^*Ndr/Ndr*^ mice and 45 wild types or heterozygous mice. Nodder mice were genotyped for the Ndr allele and sex by the Transnetyx^®^ (USA) automated qPCR genotyping company. Mice were housed in cages with enrichment and maintained on a standard day-night (12 hour) cycle, with ad libitum access to food (standard chow SDS RM3 or SDS CRM, Special Diet Services) and water. Experiments generally include a mix of mice of both male and female sex. Experiments in which the results suggested there were sex differences were expanded with additional mice of each sex to determine whether sex differences were present.

### Patient samples

Color photographs of the ocular fundii of patients with ALGS, BA and their age-matched controls were obtained after pupil dilatation using fundus cameras Canon EOS-1 Kodak Professional DCS 520C (Canon, Rochester, New York, USA) or Canon CRDG non-mydriatic retinal camera (Canon, Tokyo, Japan). Only correctly focused photographs either from eyes with both the macula and the optic disc visible, or from eyes with the optic disc in the center, were used for the analysis. Data collection and analysis was performed under the 335/00, and 2019-00202 ethical permits approved by Regional ethics review board in Stockholm. The retina fundus photographs were analyzed from seven healthy pediatric control patients (median age 8 years, range 7 – 8 years; 1 male, 6 females), six patients with ALGS (median age 7.5 years, range 2 – 11 years; 1 male, 5 females), four patients with BA (median age 10.5 years, range 8 – 16 years; 1 male, 3 females). The patients included in this study and/or their guardians gave written informed consent, and were previously reported in part ^128,135^.

Color fundus photography of patients with CADASIL and their aged-matched controls was performed using Visucam (Carl Zeiss Meditech, Germany). The data collection and analysis were approved by the IRB of the Ärztekammer Westfalen-Lippe and University of Münster (2015-402-f-S). All subjects gave written informed consent. Data from these patients have been published in part in previous studies ^132,133^. Fourteen healthy adult control patients (median age 48 year, range 23 – 61 years; 5 males, 9 females) and 14 patients with CADASIL (median age 51 year, 6 males, 8 females).

Retrospective chart analysis for patients at Children’s Hospital of Fudan University was performed in 2018, under ethical permit No (2015) 178, approved by the ethical board of Children’s Hospital of Fudan University. Retrospective chart analysis did not require informed consent.

The medical history from patients with ALGS or BA, whose retina fundus photographs were analyzed here, follows the decision 2017/1394-31 by the Regional ethics review board in Stockholm. Permission was given to retrospectively collect data from charts of patients with chronic cholestatic disease without additional consent from patients.

### Immunofluorescence staining of retina

Eyes were fixed with 3.8% Formaldehyde solution (Sigma-Aldrich, cat. #F1635) overnight and whole retinas were dissected out, blocked and permeabilized in phosphate buffered saline (PBS) containing 1% Bovine serum albumin (Sigma-Aldrich, cat. # A2153) with 0.3% TritonX-100 (Sigma-Aldrich, cat. #T8787). Primary and secondary antibodies were diluted in blocking solution. PBS : blocking solution (1:1) was used as a washing buffer. Each step was performed at 4°C, overnight. Retinas were flat-mounted in Vectashield (cat. no. H-1000, Vector Laboratories). Primary antibodies used: rat anti-mouse CD31 (cat. #553370, BD Biosciences, 1:100), rabbit anti-ERG (cat. #ab92513, Abcam, 1:200), goat-anti DLL4 (cat. #AF1389, RnD systems, dilution 1:200), goat anti-CD13 (cat. #AF2335, RnD Systems), goat anti-Jagged1 (cat. #J4127, Sigma-Aldrich, 1:500), mouse anti-human α-smooth muscle actin (Cy3 conjugated, cat. #C6198, Sigma Aldrich, 1:500; FITC conjugated, cat. #F3777, Sigma Aldrich, 1:500), rabbit anti-HISTONE H3 (phospho S10) (cat. #ab5176, Abcam, 1:500), rabbit anti-NOTCH3 (cat. #Ab23426, Abcam, 1:500), mouse anti-GFAP (Cy3 conjugated, cat. # MAB3402C3, Sigma-Aldrich, 1:500), rabbit anti-Cleaved CASPASE3 (cat. #9661 (D175), Cell Signaling, 1:700), rabbit anti-COLLAGEN type IV (cat. # AB756P, Merck Millipore, 1:500), mouse anti-Neurofilament (cat. #RT97, DSHB, 1:200) nuclei were labeled with DAPI (cat. #D9542, Sigma Aldrich, 1:1000). Images were captured using LSM 510 META or LSM 880 (Carl Zeiss AG) microscopes and processed in Image J (NIH), and/or Adobe image suite software (Adobe Inc). Any image modifications were applied identically to images being compared.

### Quantitative analysis of the retinal vasculature

#### Vascular outgrowth analysis

The distance from the optic nerve to the periphery of a retina was measured in ImageJ (NIH) using the straight line tool. Distance was measured on tile-scanned images (10x objective) of whole retina. 6 measurements, regularly radially spaced at approximately 60 degree intervals, were performed per retina from 4 *Jag1*^*+/+*^ and 4 *Jag1*^*Ndr/Ndr*^ animals at P5.

#### Tip cells and filopodia analysis

Tip cells were defined as CD31^+^Erg^+^ cells with extended filopodia at the vascular front. The number of tips per tip cell was quantified as the number of tips with filopodia bundles (single membrane protrusions coming out of a tip cell) extending in a single direction, divided by the number of tip cells. Filopodia was defined as single hair-like membrane protrusions extending from a tip cell. The number of filopodia was then divided either by the number of tip cells or by the number of tips. All quantifications were counted manually in Image J, in 40x images, 4 images per animal, in 5 *Jag1*^*+/+*^ and 6 *Jag1*^*Ndr/Ndr*^ mice at P5.

#### Quantification of vascular density, branching point and number of ERG+ endothelial cells

The number of branching points per field (40x objective) was manually quantified in the middle of the outgrowing vascular plexus (at P10, P15, adult and 1 year halfway between an arteriole and venule). The quantification of branching points was performed in 3-6 images per animal, in 5 *Jag1*^*+/+*^ and 4 *Jag1*^*Ndr/Ndr*^ mice at P5, 3 *Jag1*^*+/+*^ and 3 *Jag1*^*Ndr/Ndr*^ mice at P10, P15, and 5 *Jag1*^*+/+*^ and 5 *Jag1*^*Ndr/Ndr*^ adult mice (3 – 6 months) or aged animals (1 year). In each 40x field the number of ERG^+^ nuclei were manually quantified in 3 images per animal, in 5 *Jag1*^*+/+*^ and 5 *Jag1*^*Ndr/Ndr*^ mice at P5, in 5 images per animal, and in 3 *Jag1*^*+/+*^ and 3 *Jag1*^*Ndr/Ndr*^ mice at P10, and P15. In the same field, the CD31+ vascular length was manually measured using the free hand tool in ImageJ. The number of ERG+ endothelial cells was divided by vascular length.

#### Phosphorylated Histone3 proliferation analysis

Whole retina tile-scans (objective 10x) were taken from 4 *Jag1*^*+/+*^ and 4 *Jag1*^*Ndr/Ndr*^ mice at P5, and 3 *Jag1*^*+/+*^ and 3 *Jag1*^*Ndr/Ndr*^ mice at P10 and P15. Phospho-Histone H3+ (PH3+) CD31+ cells were manually counted in a field within a 45° wedge originating at the optic nerve, in zones of radius 200 μm, in 12 zones altogether (Supp Figure 1D). The average number of PH3+CD31+ per zone was normalized to the zone area (μm^2^) and multiplied by 100 (per 100 μm^2^ area).

#### Vessel diameter analysis

Vessel diameter was measured at a distance of 220 μm and 460 μm from the optic nerve in ImageJ for all major arterioles and venules in 3 *Jag1*^*+/+*^ and 3 *Jag1*^*Ndr/Ndr*^ mice at 1 year. The vessel diameter from the two measurements was averaged.

#### Cleaved Caspase 3

The number of cleaved caspase3 (Cl. CASP3) positive cells was quantified manually along arterioles. The arterioles were imaged in confocal z-stacks and inspected for cl. CASP3+ cells throughout the stacks. The number of branching points per image (field), the number of cl. CASP3+ cells at a branching point and the number of cl. CASP3+ outside of a branching point were manually quantified. The number of Cl. CASP3 positive cells was addressed in aged mice (>1 year old) in 7 – 13 stacks per mouse, in 5 *Jag1*^*+/+*^ and 6 *Jag1*^*Ndr/Ndr*^ mice.

#### Arterial and venous segregation

In murine retinas, the number of major arterioles and venules originating at the optic nerve, and the number of arteriovenous crossings, were quantified in whole mount retinas stained for CD31 and αSMA. Six male and 6 female *Jag1*^*+/+*^ and 4 male and 4 female *Jag1*^*Ndr/Ndr*^ adults were included for quantification of arterioles and venules. Arteriole/arteriole and arteriole/venule crossings were counted and grouped together. No venule/venule crossings were observed. Altogether 6 male and 5 female *Jag1*^*+/+*^ mice and 7 male and 5 female *Jag1*^*Ndr/Ndr*^ mice were included for quantification of aberrant crossings in adult mice (3 – 6 months).

In human retinas, the number of major arterioles and venules were quantified at the border of the optic nerve head from fundus photographs in which the optic nerve head was the center of the image. The venules were recognized by their darker color. The number of arteriovenous crossings was counted. No distinction was made between arteriovenous crossing and nicking. Patients with ALGS – 3, BA – 4, pediatric control – 6, CADASIL – 14, adult control – 14.

#### Vessel tortuosity

Blood vessel tortuosity was addressed by manually tracing the curve length of major arterioles and venules from the optic nerve towards the periphery in murine whole retina (20x objective) or human retina fundus photographs, in Adobe Illustrator. In murine retinas, only the main branches were traced for analysis. In human retinas, only branches that extended to the periphery (edge of the image) were included for analysis. The chord length was manually measured as a straight line connecting the start point to the end point of a vessel. The tortuosity was calculated by dividing the vessel curved length by its chord length. The result was transferred into percentage and subtracted by 100 to yield the % increase in tortuosity compared to straight (0% would be a perfectly straight blood vessel). The analysis was performed on whole retinas from one eye from 4 *Jag1*^*+/+*^ and 4 *Jag1*^*Ndr/Ndr*^ P30 mice, 3 *Jag1*^*+/+*^ and 3 *Jag1*^*Ndr/Ndr*^ one-year-old mice and all the patients included in the study (both eyes).

#### Vertical sprouting analysis

To address the overall number of sprouts connecting SCP to ICP and DCP to ICP a single image from a z-stack halfway between SCP to ICP and DCP to ICP was analyzed. A sprout in this image appears as a single dot. One Z-stack image (40x objective) taken in the middle of a retina between arteriole and venule was analyzed per animal. 5 *Jag1*^*+/+*^ and 5 *Jag1*^*Ndr/Ndr*^ adult mice (3 – 6 months) and 9 *Jag1*^*+/+*^ and 8 *Jag1*^*Ndr/Ndr*^ aged animals (one year) were used.

#### Superficial and intermediate capillary plexus integrity

The number of SCP and ICP branching points was quantified in z-projected images (ImageJ) of the SCP or ICP (40x objective) taken in the middle of the retina between arteriole and venule. Four images from 6 *Jag1*^*+/+*^ and 7 *Jag1*^*Ndr/Ndr*^ adult animals and 2 - 5 images from 6 *Jag1*^*+/+*^ and 5 *Jag1*^*Ndr/Ndr*^ one-year-old animals were analyzed. In the same field, the CD31+ vascular length was measured by free hand tool in ImageJ. In the same image, in 3 *Jag1*^*+/+*^ and 3 *Jag1*^*Ndr/Ndr*^ adult animals (3 – 6 months), the length of empty CollagenIV sleeves was measured using the free hand tool in ImageJ.

#### Neurofilament quantification

Neurofilament coverage was quantified in ImageJ and calculated as % of area covered by neurofilaments of the total area (100%). Neurofilament area was measured using the following script in ImageJ:

run(“Measure”);

run(“8-bit”);

run(“Median...”, “radius=2”);

setAutoThreshold(“Mean dark”);

//run(“Threshold...”);

//setThreshold(6, 255);

setOption(“BlackBackground”, false);

run(“Convert to Mask”);

run(“Watershed”);

run(“Analyze Particles...”);

Total area was measured by Measure (ctrl+M command)

The measurement was performed on 2-4 images from 3 *Jag1*^*+/+*^ and 4 *Jag1*^*Ndr/Ndr*^ P40 animals.

#### Fluorescence intensity

The blood vessel of interest was outlined using the lasso tool. The intensity was assessed by the histogram function for individual channels in Adobe Photoshop.

GFAP intensity was quantified in adults from 3 images from 4 *Jag1*^*+/+*^ and 4 *Jag1*^*Ndr/Ndr*^ mice.

JAG1 intensity was quantified in adults from 4 images from 4 *Jag1*^*+/+*^ and 4 *Jag1*^*Ndr/Ndr*^ mice.

NOTCH3 intensity was quantified in adults from 4 images from 4 *Jag1*^*+/+*^ and 4 *Jag1*^*Ndr/Ndr*^ mice.

#### ASMA gaps

The number of gaps in αSMA coverage was quantified manually. An αSMA gap was considered a gap between two vascular smooth muscle cells that is greater than one cell width (~ >10 μm). The total number of gaps per retina was counted. The number of gaps was quantified in 4 *Jag1*^*+/+*^, 3 *Jag1*^*Ndr/Ndr*^, 4 *Jag1*^*+/+*^ treated with Angiotensin II and 5 *Jag1*^*Ndr/Ndr*^ treated with Angiotensin II.

### Whole organ hemorrhage analysis and imaging

Freshly dissected brains and retinas were inspected for the presence of hemorrhages under a stereomicroscope Stemi 305 (Carl Zeiss Microscopy). Whole organs or tissue images were taken with a stereomicroscope Stemi 305 (Carl Zeiss Microscopy) combined with Canon camera (Cannon, PowerShot S3 IS). The background around the tissue was covered by a solid black color in Adobe Photoshop for esthetic purposes.

### Image processing

Capillary plexus visualization was accomplished by splitting z-stack images into 3 equally sized z-stacks per animal, color coding each plexus and merging the 3 images. Due to different sizes of *Jag1*^*+/+*^ and *Jag1*^*Ndr/Ndr*^ retinas, stack sizes were different among animals. Images at P15 and 1 year (Figure 1B, Sup Figure 2A) were further processed in ImageJ by median filter (radius = 1).

Retina whole vasculature side images were further processed in Volocity in which high opacity and noise reduction filters were included.

### Liver resin casting

Adult liver portal vein vasculature was injected with synthetic resin MICROFIL^®^ and scanned with micro computed tomography as previously described ^102^.

### Micro CT analysis of ruptures

Following the micro-CT measurements, the reconstructed data was imported in VG Studio MAX software (Volume Graphics GmbH, https://www.volumegraphics.com). The Microfil-filled structures were separated from the background by global thresholding creating a main region of interest (ROI), then the areas of Microfil leaked from ruptured vessels were manually selected into separate ROIs. The volume of each ROI of leaked Microfil was calculated by the software. The analysis was performed in 4 *Jag1*^*+/+*^, 4 *Jag1*^*Ndr/Ndr*^ samples (analyses of portal vein and biliary architecture, but not vascular leakage data, for 3 *Jag1*^*+/+*^ and 3 *Jag1*^*Ndr/Ndr*^ samples of these casts were previously published in ^102^).

### Skull micro computed tomography

Micro-computed tomography (microCT) was performed on 7 and 8 month old skulls (3 *Jag1*^*+/+*^ and 3 *Jag1*^*Ndr/Ndr*^ mice, all males) using a MicroCT50 (Scanco Medical), 55kVp, 109μA, 6W at 20μm voxel size, with a 500 ms integration time and a 20.5 mm field of view.

### Skull thickness and volume analysis

The skull thickness and volume analyses were conducted in VG Studio MAX 3.4 (Volume Graphics GmbH, Germany). A sample was registered within the coordinate system, and the skull surface was determined, including the jaws. The volume was calculated by multiplying the number of voxels of a given surface by the volume of one voxel. The skull thickness was measured by utilizing a ray, searching for the opposite surface for each point on the skull surface. The resulting thickness map depicts the shortest distances between the inner and outer surface. The thickness and volume of the skulls were measured in two different ways, one of the whole skull thickness and one taking into account the trabeculation of the spongy bone. The spongy bone segmentation was conducted using the VGEasyPore module. The analysis was performed in 3 *Jag1*^*+/+*^, 3 *Jag1*^*Ndr/Ndr*^ adult samples.

### Skull length

The skull length was measured in Image J using the straight line tool. The length was measured from the occipital bone to the nasal bone from a dorsal view. The measurement was performed for 3 *Jag1*^*+/+*^, 3 *Jag1*^*Ndr/Ndr*^ skulls.

### Vessel permeability assay

Mice were injected with 200 μl 0.5% Evans blue (cat. #E2153, Sigma-Aldrich) via the tail vein. The dye was allowed to circulate for 17 hours. Mice were anesthetized by CO2 inhalation and transcardially perfused with Hanks buffered salt solution (HBSS), at a perfusion rate of 5 ml/min for 3 min. Internal organs were dissected out, weighed, and placed in Formamide solution (cat. # 15515-026, Invitrogen) for 17 hours at 56°C. Formamide solution containing Evans blue extracted from the organs was measured for absorbance at 610 nm in a VersaMax™ microplate reader (Molecular Devices Versa Max microplate reader). The amount of Evans blue measured was divided by the organ weight. Brain vasculature was inspected under the stereomicroscope Stemi 305 (Carl Zeiss Microscopy) and photographed with a camera (Cannon, PowerShot S3 IS).

### Blood pressure measurements

Blood pressure was measured using the CODA^®^ High Throughput (Kent Scientific) tail cuff system. The animals were awake, on a heating pad, and in a restraining device, during the experiment. The animals were accustomed to the tail cuff for 5 min before data recording. The tail temperature was assessed between 32 – 34°C. The systolic, diastolic and mean blood pressure were recorded. The mean baseline blood pressure was calculated as an average from 5 recordings on 5 days. In the Angiotensin II experiment, the mean blood pressure is the average of nine recordings. The increase in blood pressure was calculated by comparing the average mean blood pressure before Angiotensin II treatment (100%) and during two weeks of Angiotensin II treatment.

### Angiotensin II treatment

Mice were treated with Angiotensin II (cat. #A6402, Sigma-Aldrich) for two weeks. Angiotensin II was delivered via osmotic mini pumps (Alzet®, 2002 model) and the infusion rate was 0.025 μg/g/h. The animals were anesthetized by Isoflurane inhalation (~2%) and IP injected with 200 μl Rimadyl (0.5 mg/ml) prior to the surgery to relieve pain. The osmotic mini pumps were implanted subcutaneously and the incision was sealed with surgical glue. Animals in the experiment were weighed daily and the blood pressure was measured to verify the effect of Angiotensin II treatment. 2 of 11 *Jag1*^*+/+*^ mice, and 1 of 8 *Jag1*^*Ndr/Ndr*^ mice demonstrated a strong reaction to Angiotensin II treatment (weight loss > 20% and piloerection) and had to be sacrificed. 5 *Jag1*^*+/+*^ and 2 *Jag1*^*Ndr/Ndr*^ mice had weak or no response to Angiotensin II (no blood pressure increase) and were therefore excluded from the analysis. Animals were given extra “porridge” during the treatment to avoid dehydration. Control animals were implanted with mini-pumps loaded with Dulbecco’s phosphate buffer saline (DPBS), in total 4 *Jag1*^*+/+*^ and 5 *Jag1*^*Ndr/Ndr*^ mice.

### Body weight

*Jag1*^*+/+*^ and *Jag1*^*Ndr/Ndr*^ body weight was recorded from the first day of blood pressure measurement and throughout the Angiotensin II treatment. In the body weight graph, day 0 is body weight before the Angiotensin II treatment and day 1 is body weight from the first day of Angiotensin II treatment. Body weight change is calculated by comparing the body weight from the last day of Angiotensin II treatment to the body weight from the first day of AngII treatment (100%).

### Thrombin-Antithrombin ELISA

Thrombin-Antithrombin (TAT) complexes were measured by ELISA using a commercial kit (cat. #ab137994, Abcam). The assay was performed according to manufacturer’s instructions. The chromogen substrate was incubated for 20 min and the absorbance was read at 450 nm using VersaMax™ microplate reader. The TAT ELISA was performed on P10 plasma (4 *Jag1*^*+/+*^ and 4 *Jag1*^*Ndr/Ndr*^ mice).

### Plasma collection

Blood from P10 mice was collected from the trunk after decapitation. To obtain blood plasma the blood was mixed with 10 μl of Heparin (LEO Pharma) and kept cold throughout handling. Afterwards, the blood-Heparin mix was centrifuged at 3000 g for 15 mins at 4°C. Plasma was carefully removed from the top layer, placed in a new tube, and frozen (−20°C short term (max 2 weeks), −80°C long term storage).

### Liver enzyme analysis

Liver enzymes (including total bilirubin) were analyzed from blood plasma of P10 mice diluted six times with DPBS. The blood plasma was tested using mammalian liver profile (Abaxis, PN 500-1040) and VetScan2 system (Abaxis Inc).

### Survival analysis

Newborn pups were observed and counted daily during the first 10 days after birth. The number of deceased pups was noted. The pups were biopsied, tattooed and genotyped. 23 *Jag1*^*+/+*^ and 16 *Jag1*^*Ndr/Ndr*^ were included in the analysis, 6 *Jag1*^*Ndr/Ndr*^ pups died during the first 10 days.

### Electron microscopy

Animals were anesthetized and transcardially perfused with HBSS and electron microscopy fixative (2% formaldehyde, 2.5% glutaraldehyde and 0.02% sodium azide in 0.05 M sodium cacodylate buffer, pH 7.2) as previously described ^140^. For retina and heart analysis 6 *Jag1*^*+/+*^ and 6 *Jag1*^*Ndr/Ndr*^ mice were used. Images were further processed in Adobe Photoshop to pseudocolor cell types.

### Quantitative real time qPCR

RNA was isolated from adult whole retinas of 3 *Jag1*^*+/+*^ and 3 *Jag1*^*Ndr/Ndr*^ mice. mRNA was extracted using the RNeasy Mini Kit (QIAGEN, cat. #74104), including on-column DNase I digestion (QIAGEN, cat. #79254). Complementary DNA was synthesized from 1 μg total RNA using the Thermo Scientific™ Maxima™ First Strand cDNA Synthesis Kit for RT-qPCR (Thermo Scientific, cat. #K1641) according to the manufacturer instructions. Quantitative real-time PCR (qPCR) was performed as described ^155^. Primers used for qPCR are listed below. mRNA levels are normalized to the average housekeeping RNA levels of 18S and β-actin.

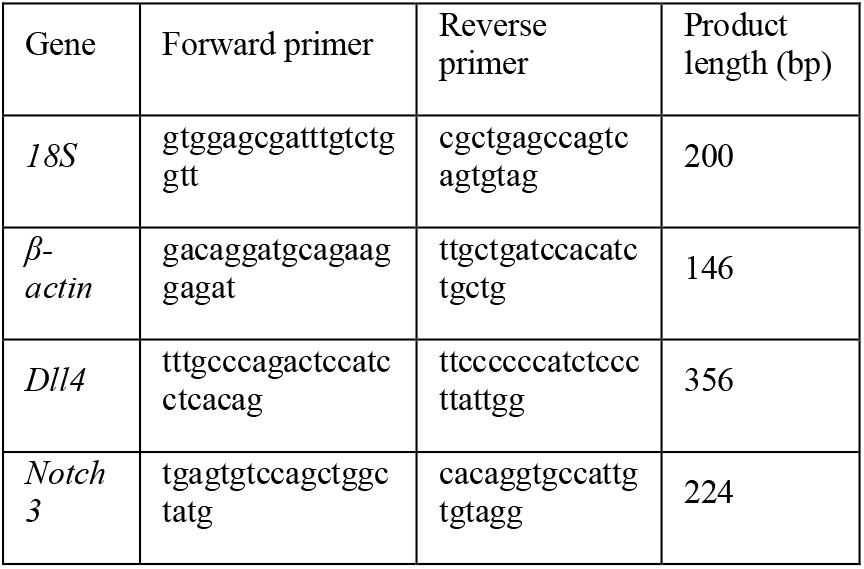

### Statistical analysis

Statistical analyses of differences between *Jag1*^*+/+*^ and *Jag1*^*Ndr/Ndr*^ animals was evaluated using two-sided unequal variance *t* test or Mann-Whitney test. When more than 2 conditions were compared, a one-way of two-way ANOVA combined with Sidak’s multiple comparisons test was used, as appropriate and as described in figure legends. Pearson correlation was used for correlation analysis. *P* value was considered as statistically significant if *p*<0.05 (*p*<0.05 = *, *p*<0.01 = **, *p*<0.0001 = ***).

## Abbreviations

A: artery/arteriole
ALGS: Alagille syndrome
AngII: Angiotensin II
ASMA: alpha smooth muscle cell actin
AV: arteriovenous
BA: biliary atresia
BP: blood pressure
cCasp3: cleaved Caspase 3
CADASIL: cerebral autosomal dominant arteriopathy with subcortical infarcts and leukoencephalopathy
CD31: PECAM-1
ColIV: Collagen IV
CTRL: control
D: deep
DCP: deep capillary plexus
Dll4: Delta-like 4
EB: Evans blue
EC: endothelial cell
Erg: ETS-related gene
g: gram
GI: gastrointestinal
GFAP: Glial fibrillary acidic protein
I: intermediate
Hg: mercury
IC: intracranial
ICP: intermediate capillary plexus
Jag1: Jagged1
mm: millimeter
n: number of biological replicates
Ndr: Nodder
NF: neurofilament
ns: not significant
OD: optical density
P: postnatal day
PAS: pulmonary artery stenosis
PH3: phospho-histone 3
S: superficial
SCP: superficial capillary plexus
TAT: thrombin-antithrombin
V: vein/venule
VSMC: vascular smooth muscle cell
mCT: micro computed tomography
mm: micrometer

## Author contributions

SH conceptualization, data curation, formal analysis, investigation, visualization and writing of original draft. NVH investigation and writing – review and editing. JL investigation, formal analysis and writing review and editing. NH investigation, validation and visualization. EV investigation, validation and visualization. TZ conceptualization and project administration. FD investigation. MK investigation and writing – review and editing. JS investigation. LL investigation. MS investigation. BRJ investigation. MH investigation. VB funding acquisition and writing – review and editing. UL funding acquisition and writing – review and editing.

AJ funding acquisition, investigation. JSW investigation, resources. FA resources and writing – review and editing. KTF resources and writing – review and editing. BF investigation, resources and writing – review and editing. JK funding acquisition, project administration, supervision and writing – review and editing. ERA conceptualization, funding acquisition, project administration, supervision and writing – original draft.

## Acknowledgments

We would like to thank Soniya Savant for her input regarding vascular phenotypes and single cell sequencing library preparation (not included in manuscript). We would like to thank Katarina Tiklova for her advice and help with single cell sequencing library preparation and FACS sorting (not included in manuscript). We thank Liqun He and Christer Betsholtz for their help with single cell analysis (not included in manuscript). Further, we would like to thank Tomasz Krzywkowski and Mats Nilsson for their efforts to help us adapt retina padlock probe in situ hybridization to retina (not included in manuscript). We would also like to thank the Rickard Sandberg group especially Gösta Winberg for providing us with Tn5 enzyme and other buffers for SmartSeq2 libraries and their input on the library quality (not included in manuscript). We would like to thank SciLife for their help with single cell RNA sequencing and bioinformatic analysis (not included in manuscript). The monoclonal antibody NF-RT97 developed by Wood, J. obtained from the Developmental Studies Hybridoma Bank, created by the NICHD of the NIH and maintained at The University of Iowa, Department of Biology, Iowa City, IA 52242.

## Supplementary Figures

**Supplementary Figure 1.**
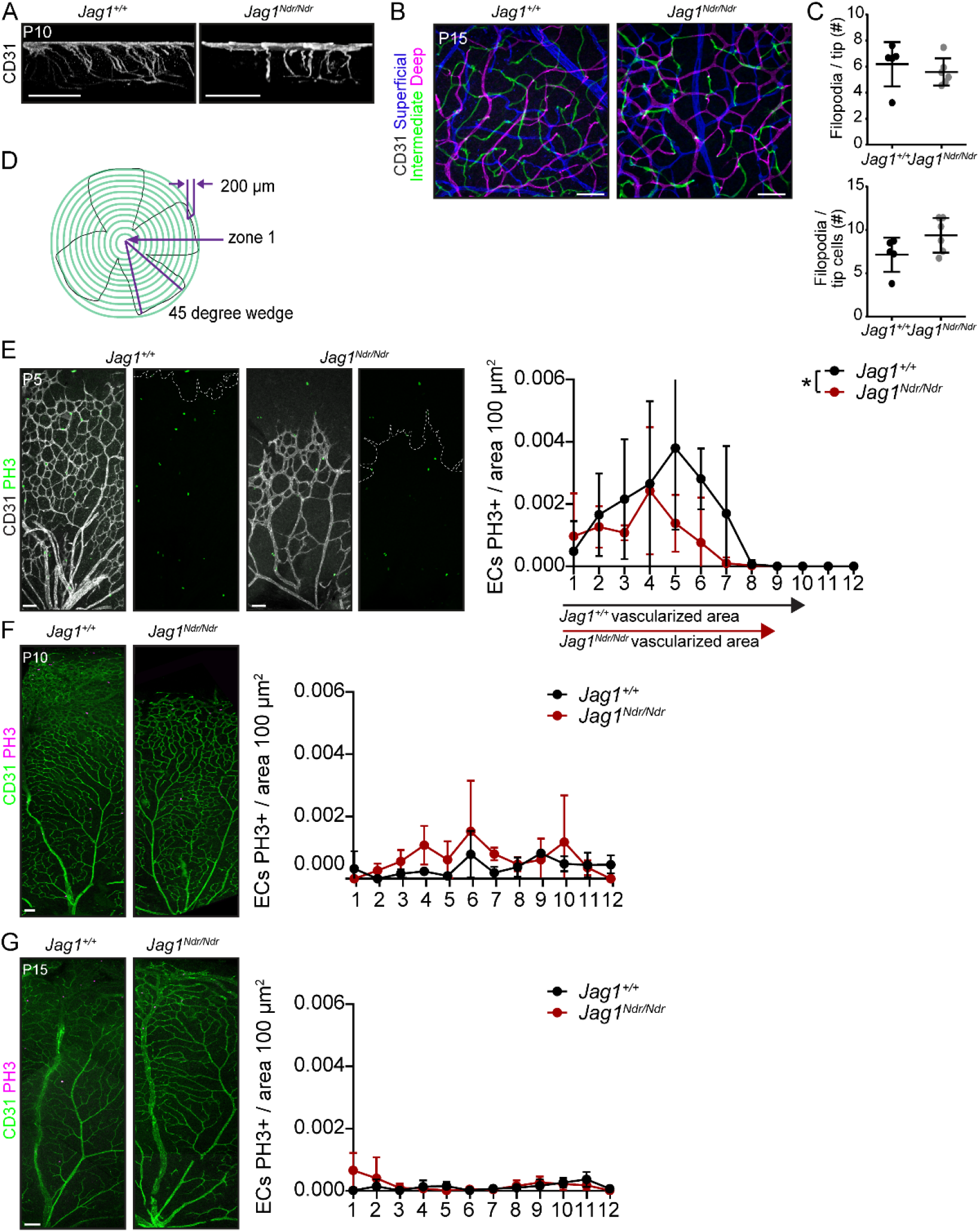
Delayed retinal vascular outgrowth and remodeling in *Jag1*^*Ndr/Ndr*^ mice. **(A)** Capillary extension into deep capillary plexus at postnatal day 10 (P10). Scale bars 100 μm. **(B)** Overview of three pseudocolored retinal capillary layers at P15. Scale bars 50 μm. **(C)** Quantification at P5 of number of filopodia per tip (top panel) and filopodia per tip cell (bottom panel). **(D)** Schematic representing radial zone delineation for quantification of phosphorylated histone 3 (PH3) positive endothelial cells. The CD31+PH3+ cells were counted in 200 μm zones in a 45° wedge (4 wedges per retina). **(E)** Immunofluorescence of PH3+ proliferating CD31+ endothelial cells at P5. The dotted line labels the edge of the vascular front. Quantification of the number of proliferating cells per radial zone, normalized to area size at P5. Scale bars 50 μm. **(F, G)** Number of proliferating endothelial cells per radial zone, normalized to area size at **(F)** P10 and **(G)** P15. Scale bars 100 μm. Statistical significance was tested using unpaired Students t-test (C) or two-way ANOVA followed by multiple comparisons (E – G). In C, each dot represents one animal; graphs depict mean values ± standard deviation, in E each dot represents the mean value ± standard deviation from 4 animals per genotype in F and G each dot represents the mean value from 3 animals per genotype; p < 0.05 (*).

**Supplementary Figure 2.**
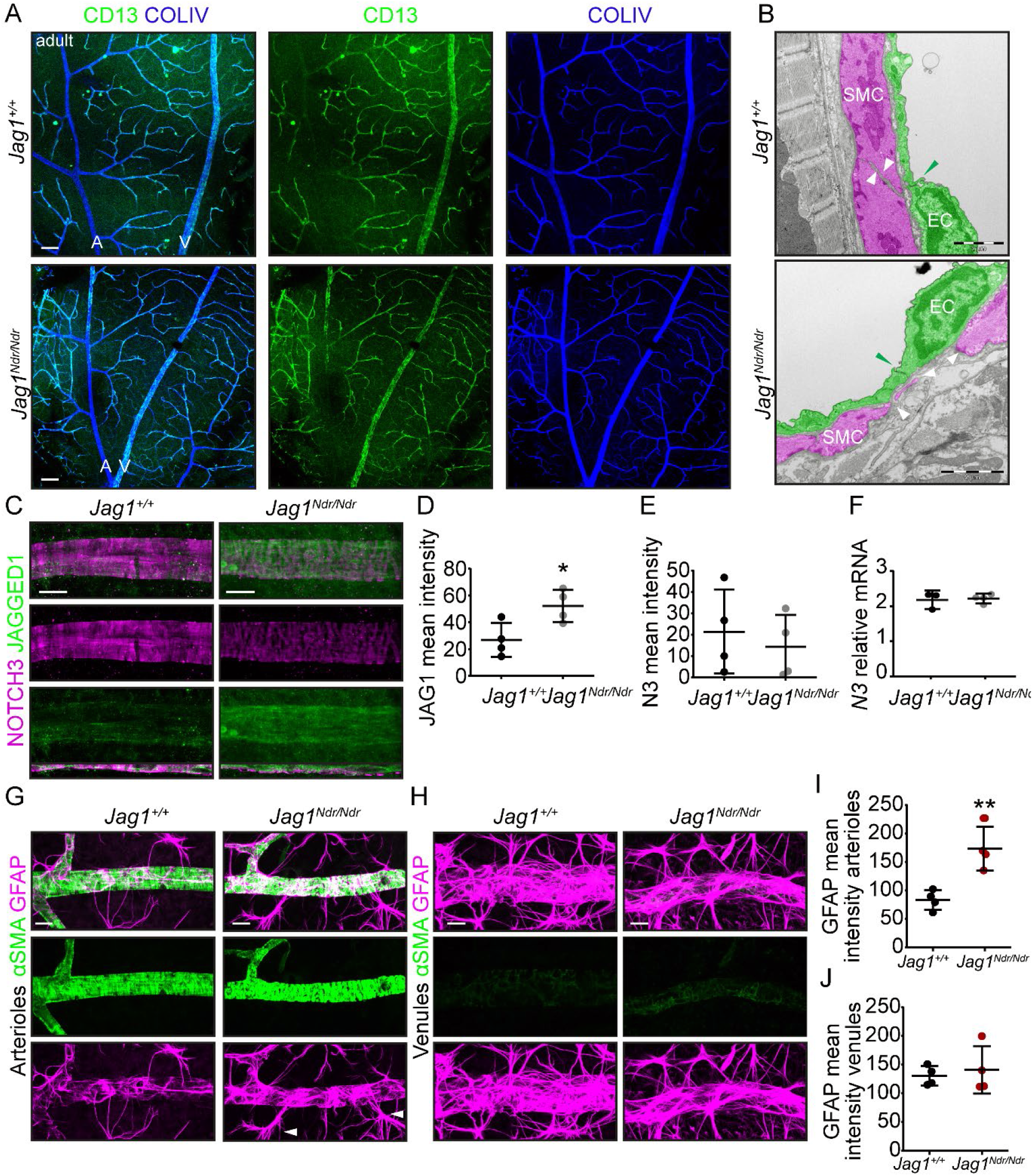
*Jag1*^*Ndr/Ndr*^ mice display CADASIL-like sparse vascular smooth muscle cell coverage of arteries with an increase in artery-associated reactive astrocytes. **(A)** Pericyte coverage of blood vessels stained for CD13 was not reduced in *Jag1*^*Ndr/Ndr*^ mice. Scale bars 100 μm. **(B)** Transmission electron microscopy of coronary arteries of adult mice. Vascular smooth muscle cells (SMC) are pseudo colored in magenta and endothelial cells (ECs) in green. White arrowheads label SMC edges and the distances between SMCs. Green arrowhead marks the tight junctions. Scale bars 2 μm. **(C)** JAG1 and NOTCH3 expression in retinal arterioles. Scale bars 10 μm. **(D)** JAG1 mean intensity was significantly increased in *Jag1*^*Ndr/Ndr*^ mice. **(E)** NOTCH3 mean intensity on arterioles and mRNA levels **(F)** in whole retina lysates. **(G – J)** GFAP+ astrocytes are more prevalent around *Jag1*^*Ndr/Ndr*^ **(G)** arterioles **(H)** but there was no difference around veins. White arrowheads label reactive astrocytes. Scale bars 20 μm. **(I, J)** GFAP mean intensity on retinal **(I)** arterioles and **(J)** venules. Statistical significance was tested using unpaired Students t-test (D – F, I, J). Each dot represents one animal; graphs depict mean values ± standard deviation; p < 0.05 (*), p < 0.01 (**). A, arteriole; EC, endothelial cell; SMC, smooth muscle cell; V, venule.

**Supplementary Figure 3.**
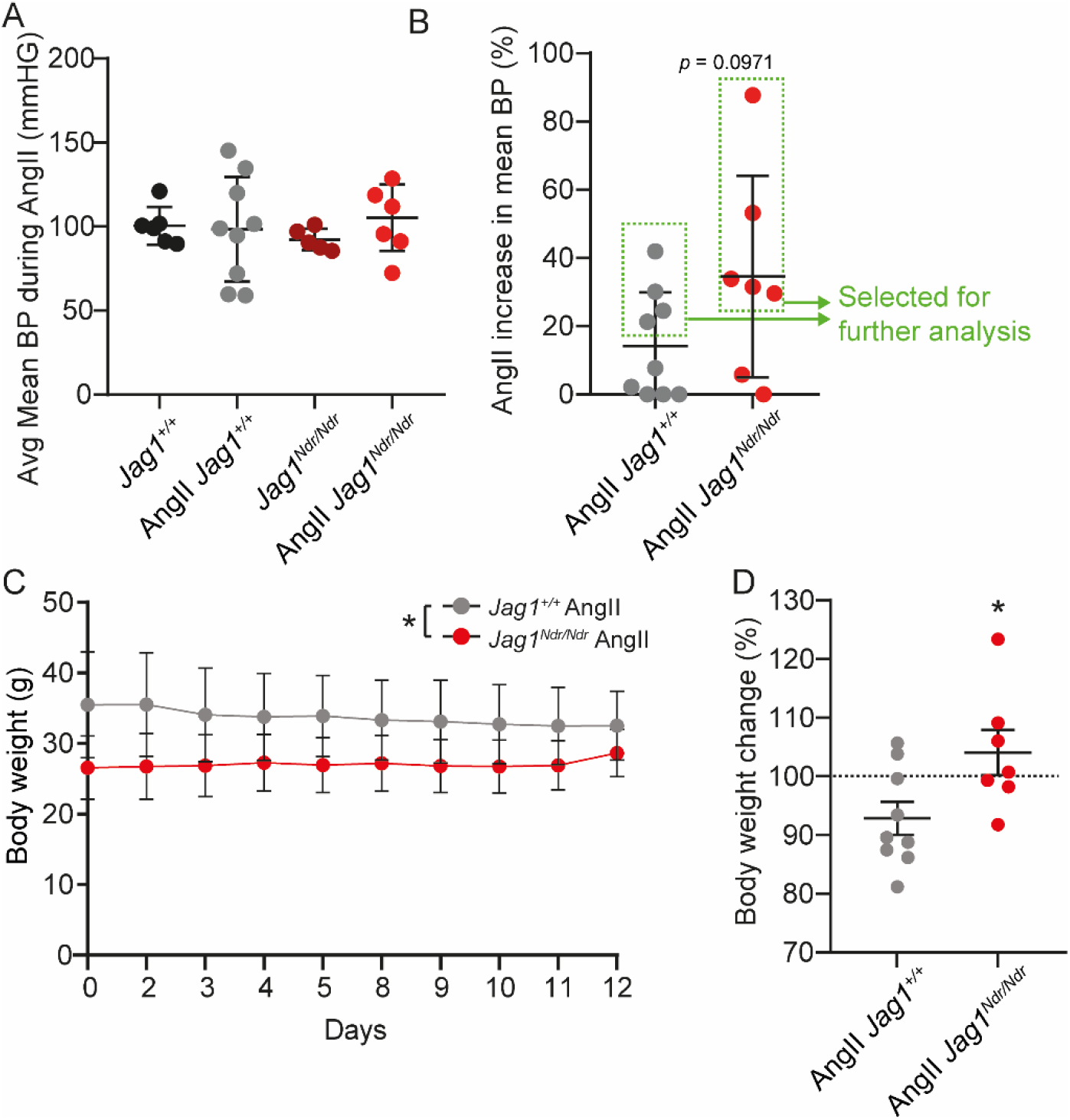
Animal response to Angiotensin II treatment, and selection of responsive animals for analysis. **(A)** Average mean blood pressure during Angiotensin II treatment of all animals. **(B)**Change in the mean blood pressure during Angiotensin II treatment compared to average before the treatment. Mice with no change in blood pressure, or a decrease in blood pressure, were set to zero for this analysis. Green arrows and boxes demarcate the animals that responded with a minimum 20% increase in mean blood pressure and were used in further analysis (Figure 7). **(C)** Body weight of all mice during treatment with Angiotensin II. In total 9 *Jag1*^*+/+*^ and 7 *Jag1*^*Ndr/Ndr*^ mice. **(D)** Change in body weight at day 12 of Angiotensin II treatment compared to day 0 (before Angiotensin II treatment). Statistical significance was tested using unpaired Students t-test (B, D) or two way ANOVA (A, C). In A, B and D, each dot represents one animal; graphs depict mean values ± standard deviation; in C each dot represents the mean values from 9 and 7 mice as described above ± standard deviation; p < 0.05 (*). AngII, Angiotensin II.

**Supplementary Figure 4.**
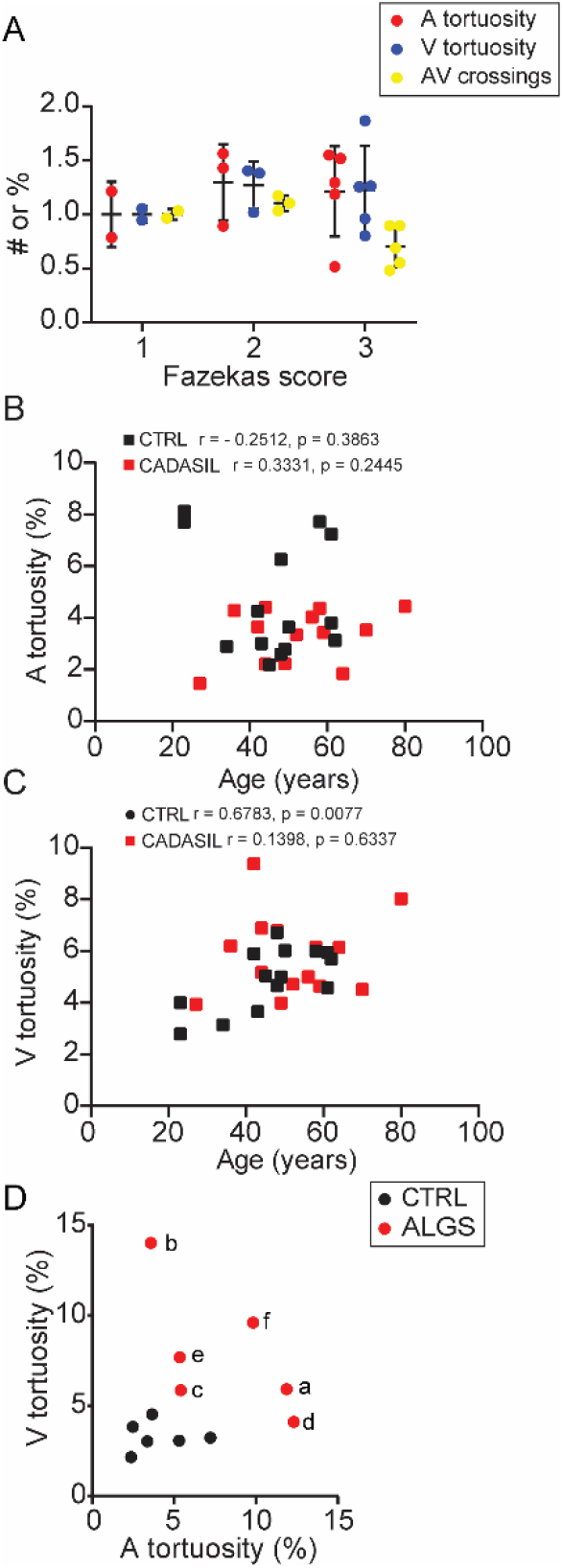
Correlation of retinographs vascular abnormalities with different factors. **(A)** Lack of correlation for arteriolar and venous tortuosity and arteriovenous crossing to Fazekas scores for CADASIL and control patients. The data for each parameter are normalized to the average value of each parameter corresponding to Fazekas score 1 value. **(B)** Correlation analysis between arterial tortuosity and age for CADASIL and age-matched control patients. **(C)** Correlation analysis between venous tortuosity and age for CADASIL and age-matched control patients. **(D)** Correlation analysis between arterial and venous tortuosity in patients with Alagille syndrome and pediatric controls. Statistical significance was tested by Pearson correlation (A – D). Each dot represents one individual; graphs depict mean values ± standard deviation. A, arteriole; ALGS, Alagille syndrome; AV, arteriovenous; CADASIL, cerebral autosomal dominant arteriopathy with subcortical infarcts and leukoencephalopathy; V, venule.

## Supplementary Table

**Supplementary table 1.**
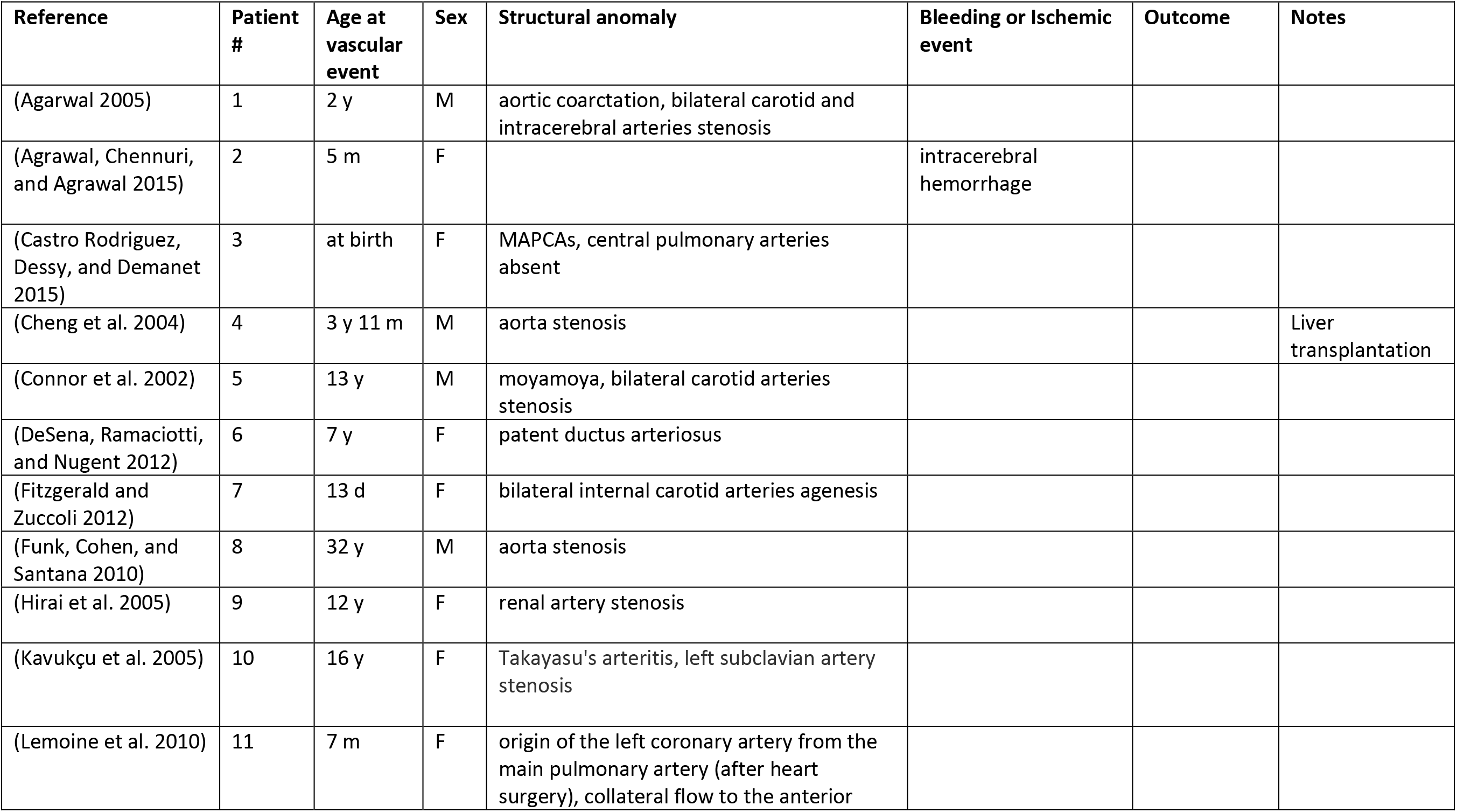
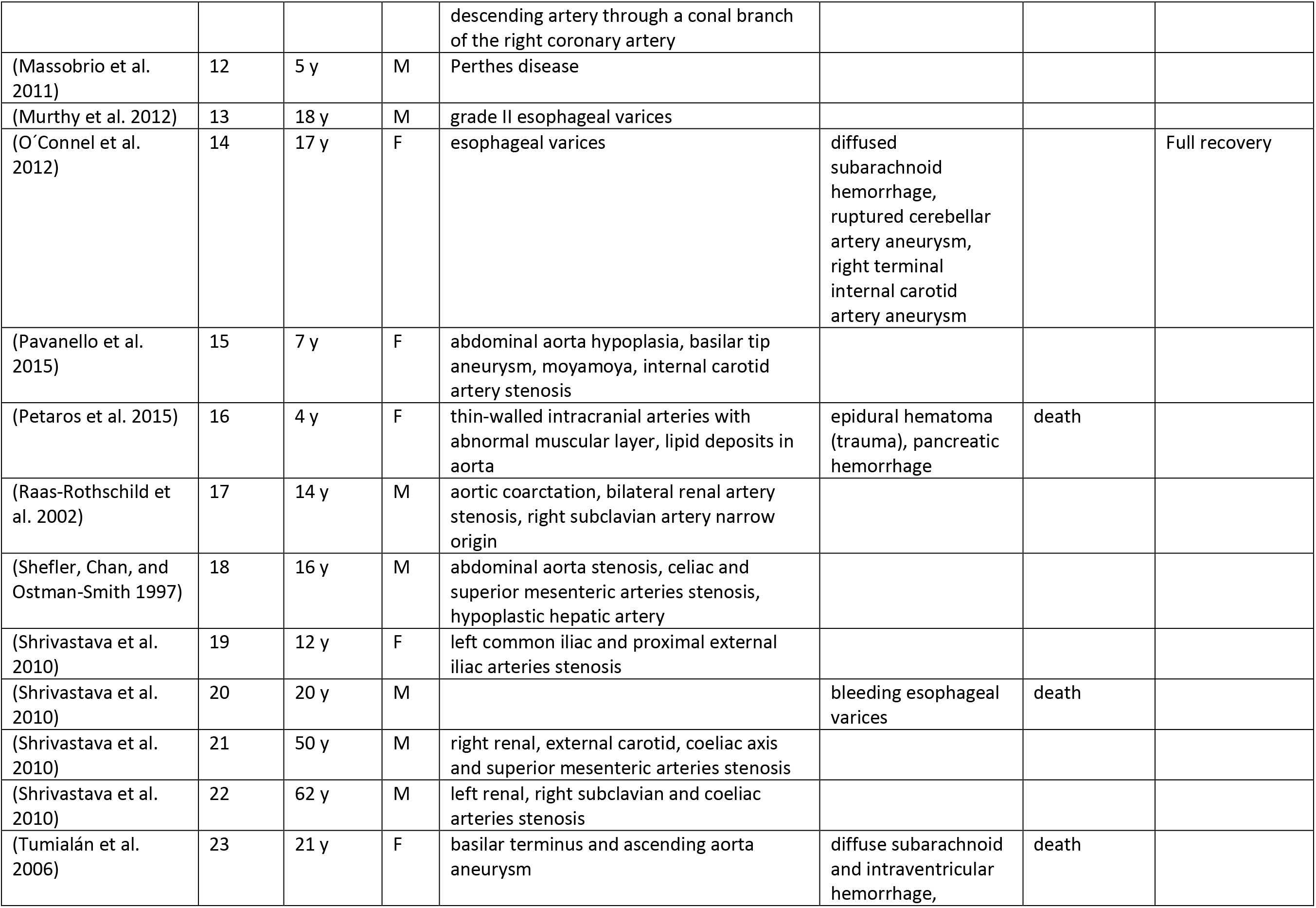
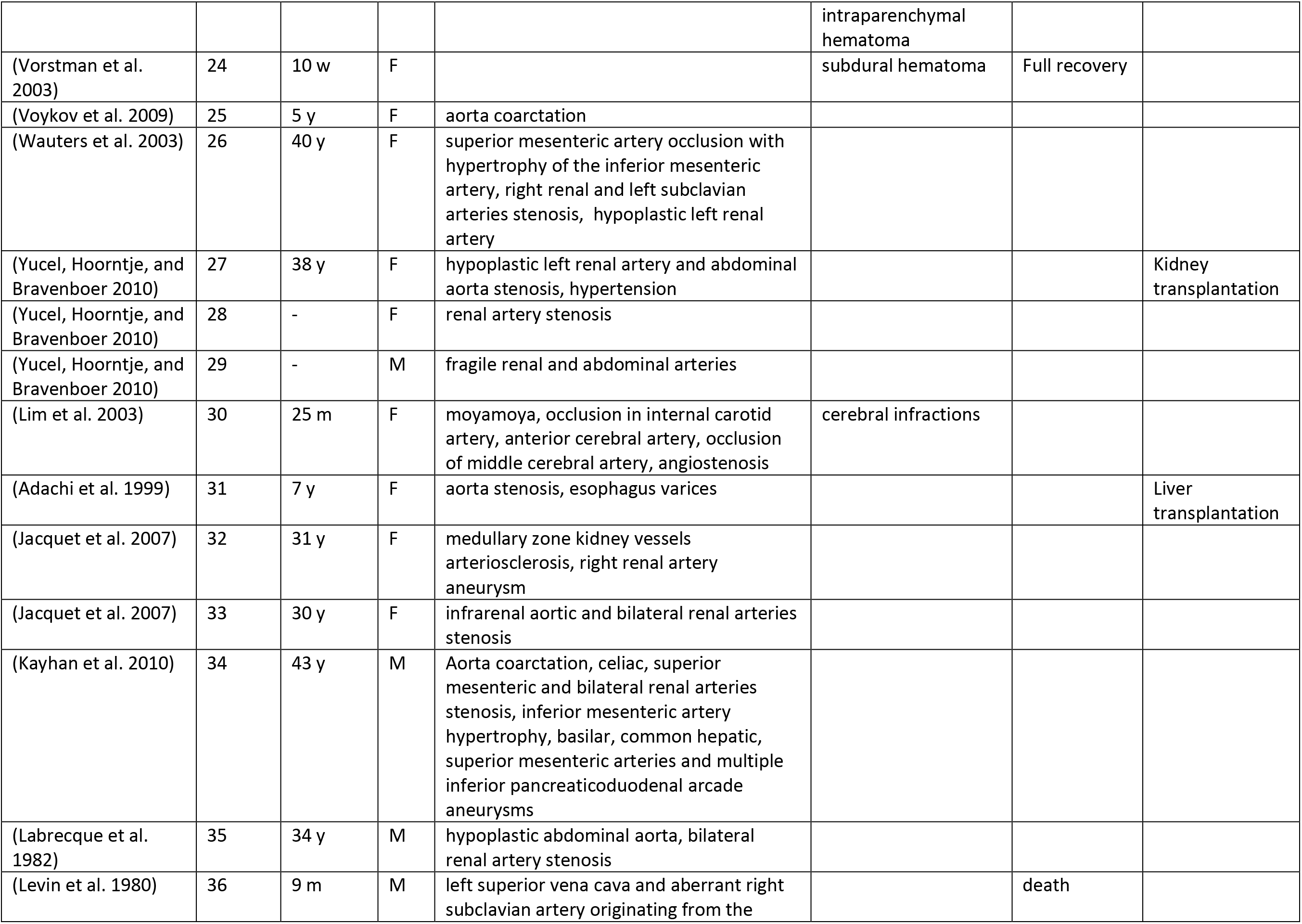
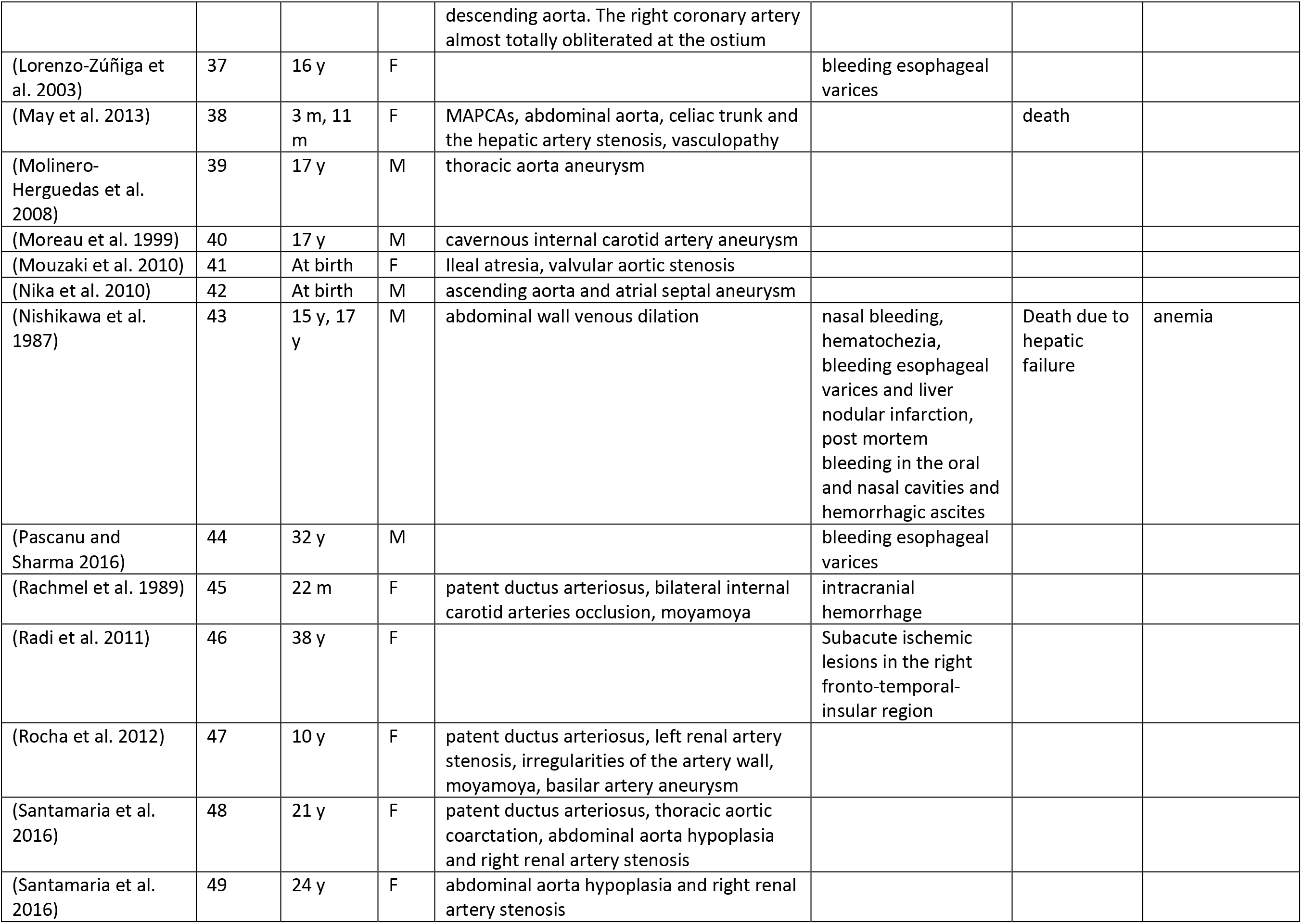
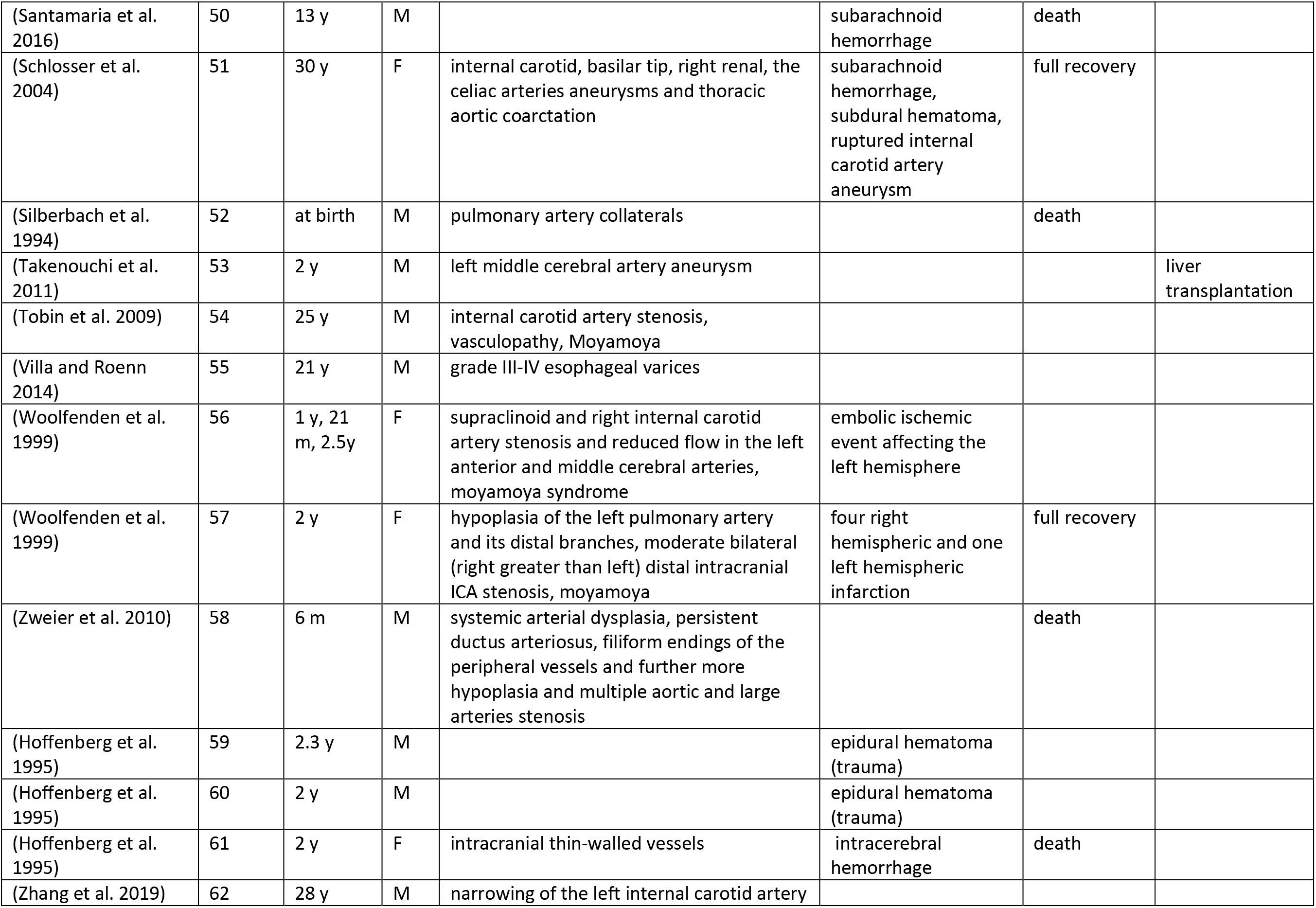
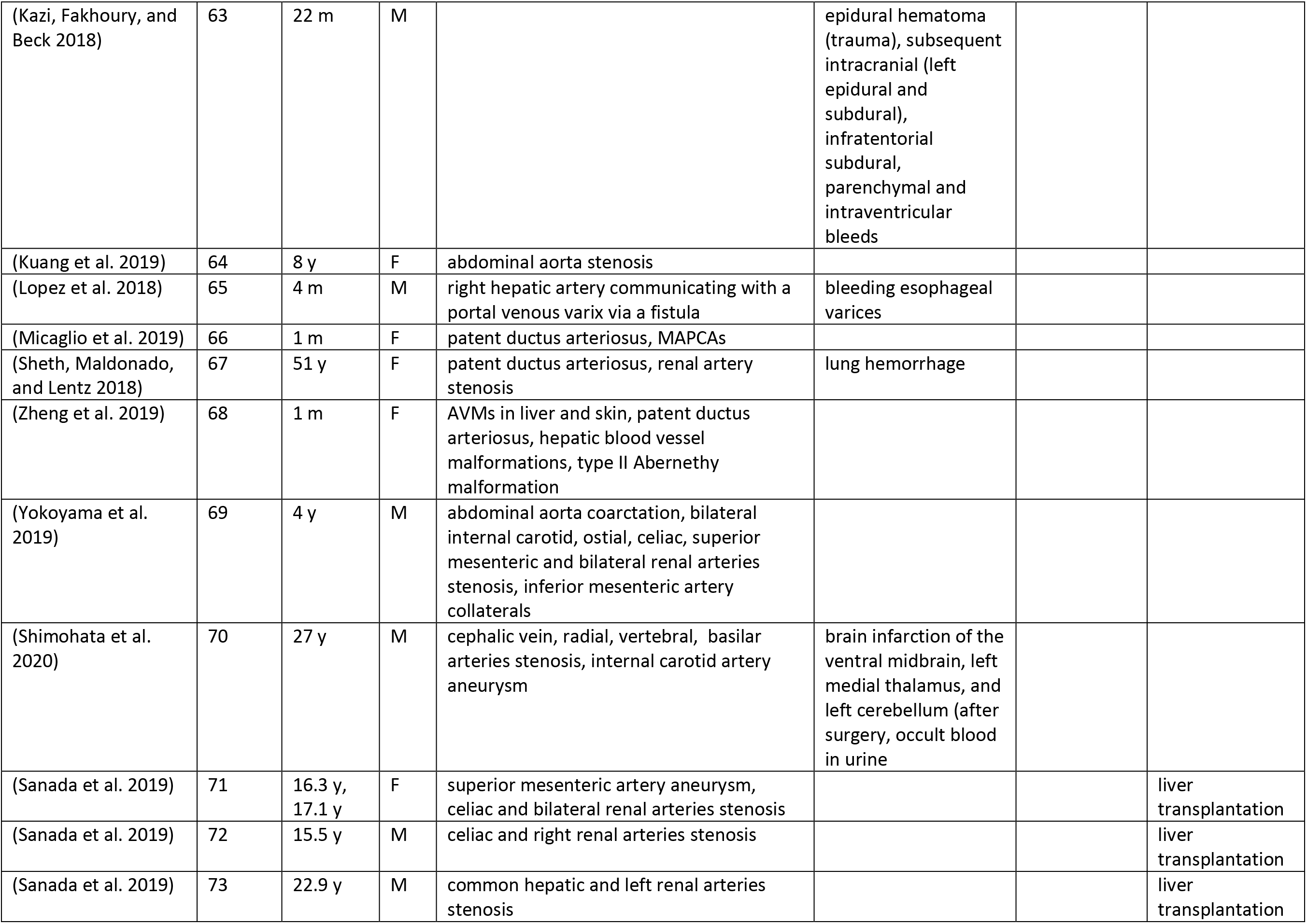
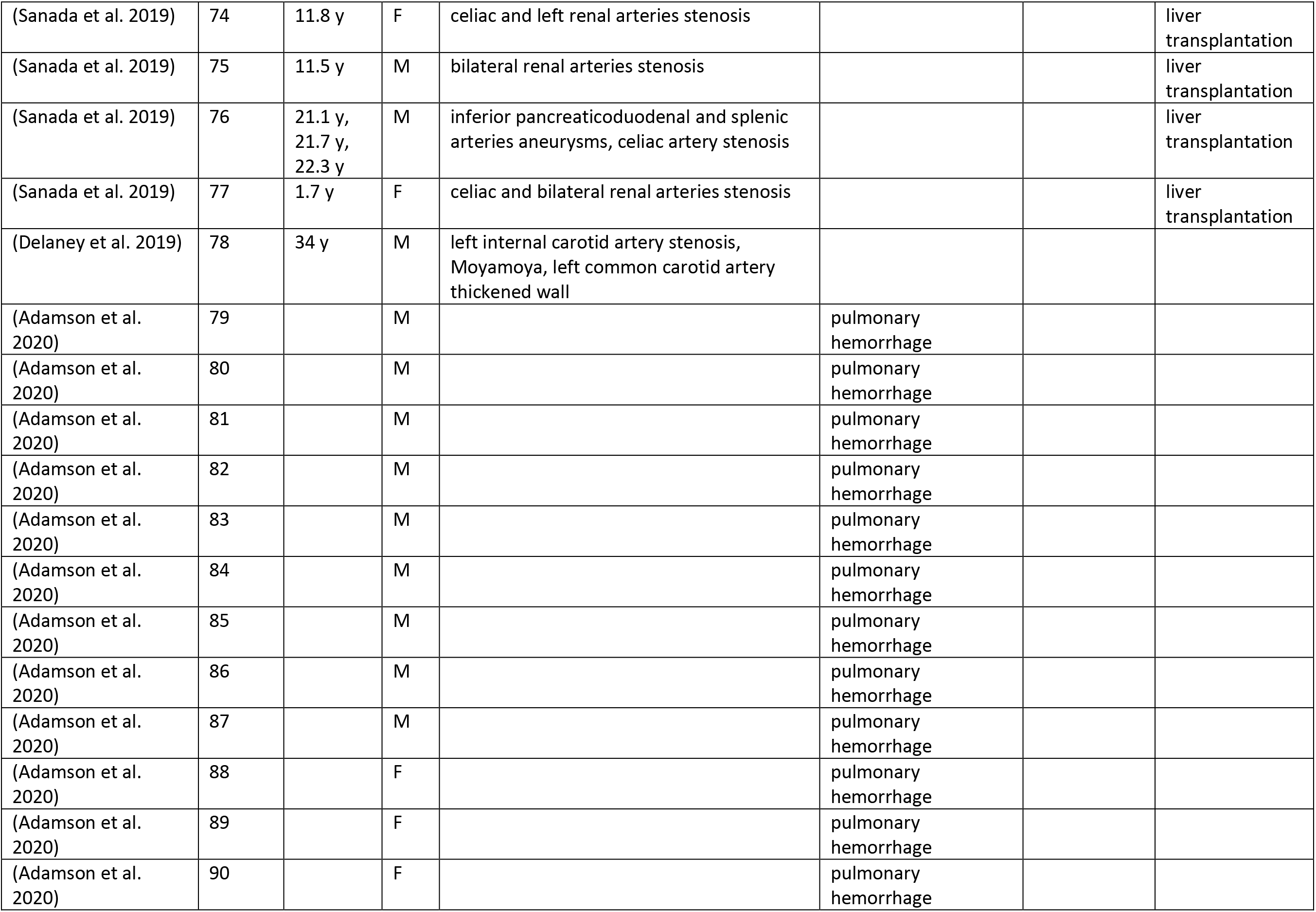
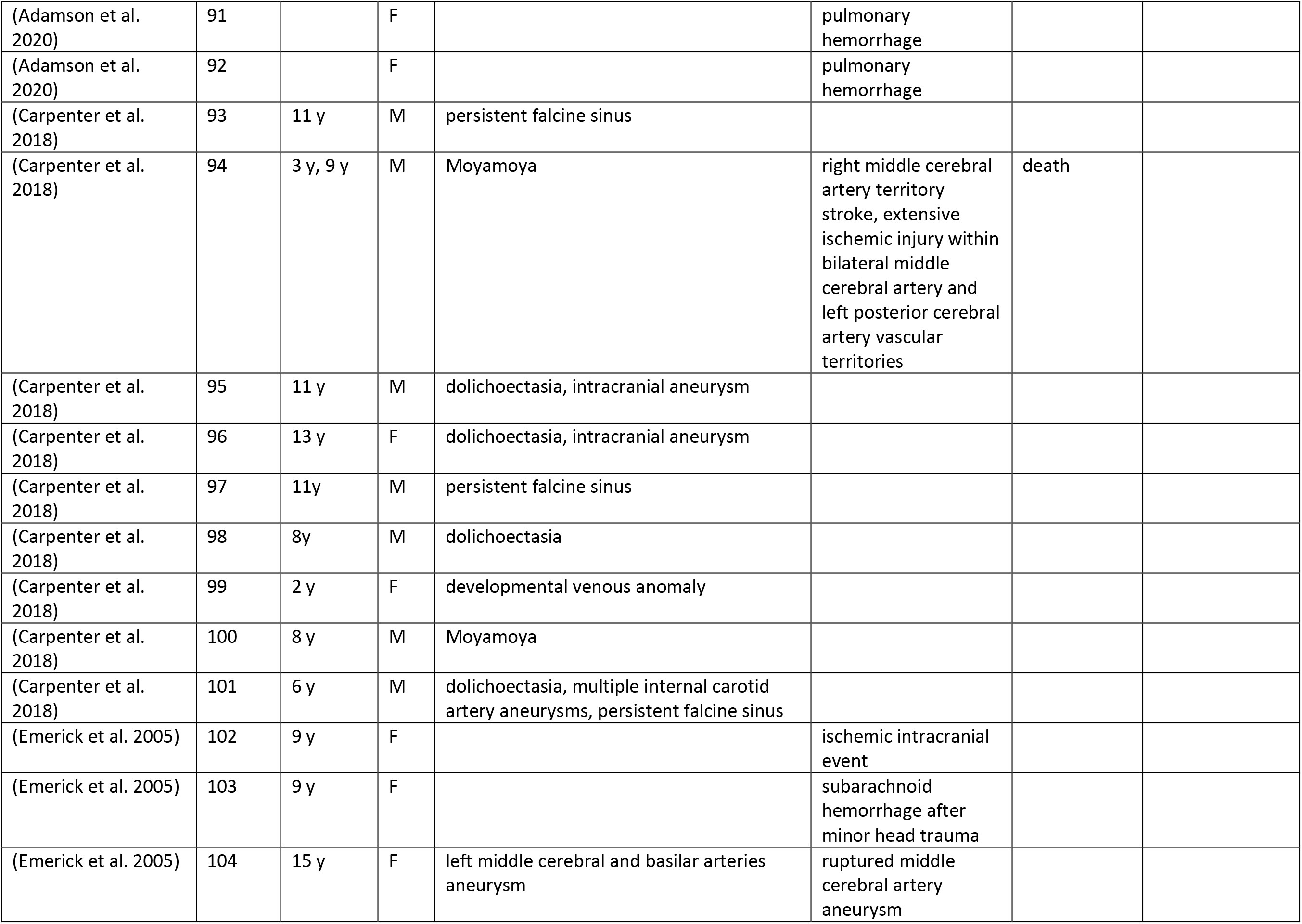
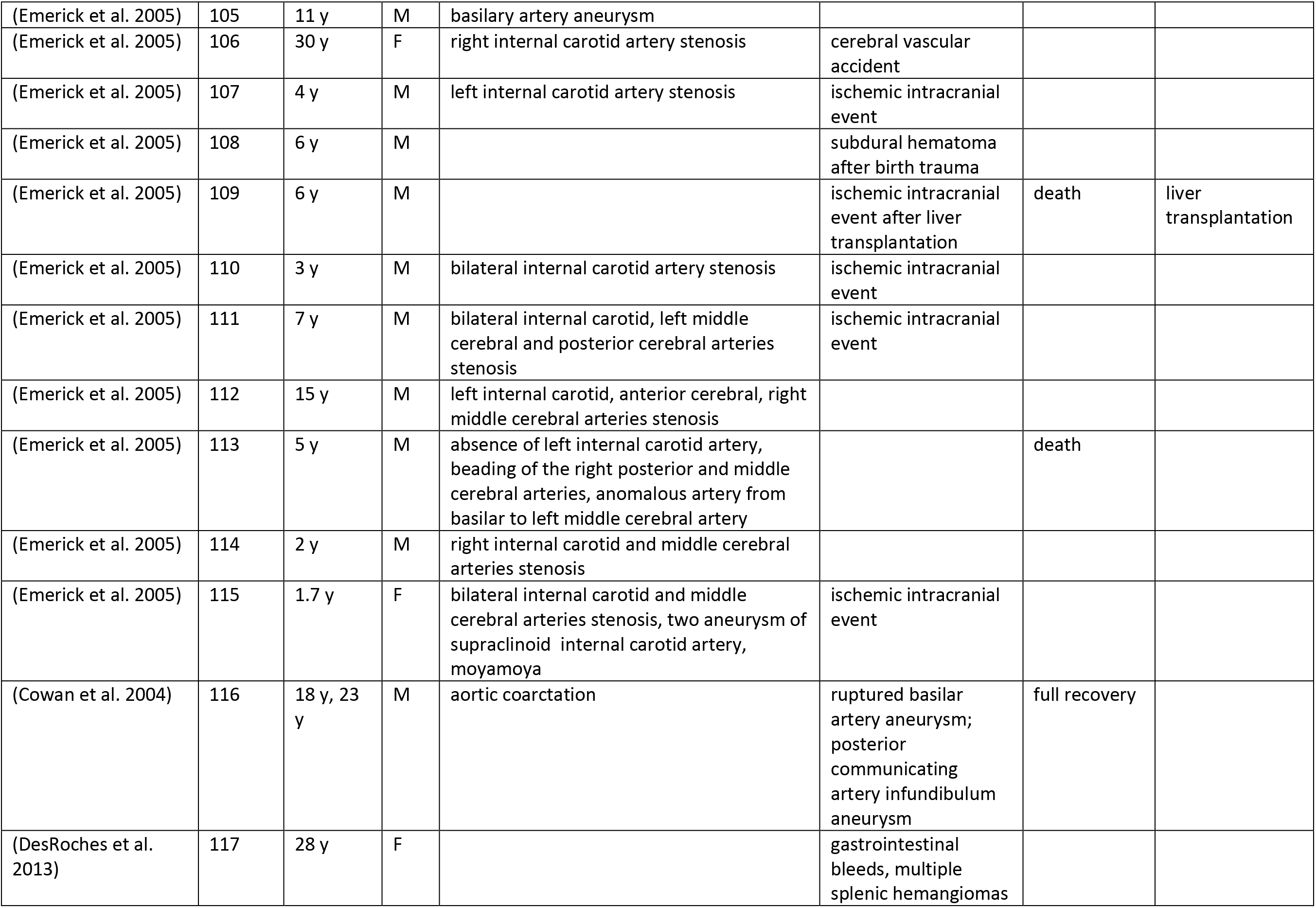
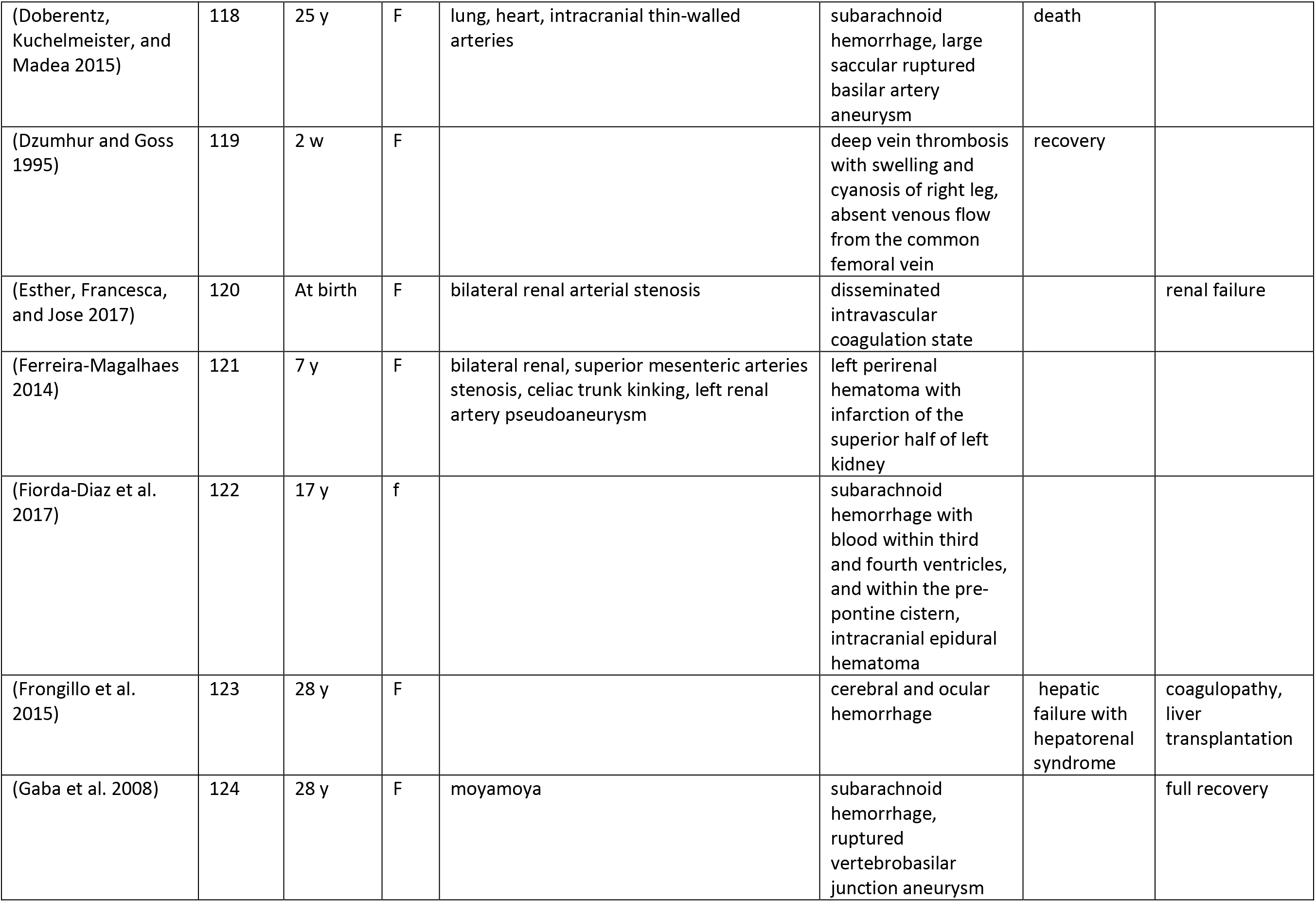
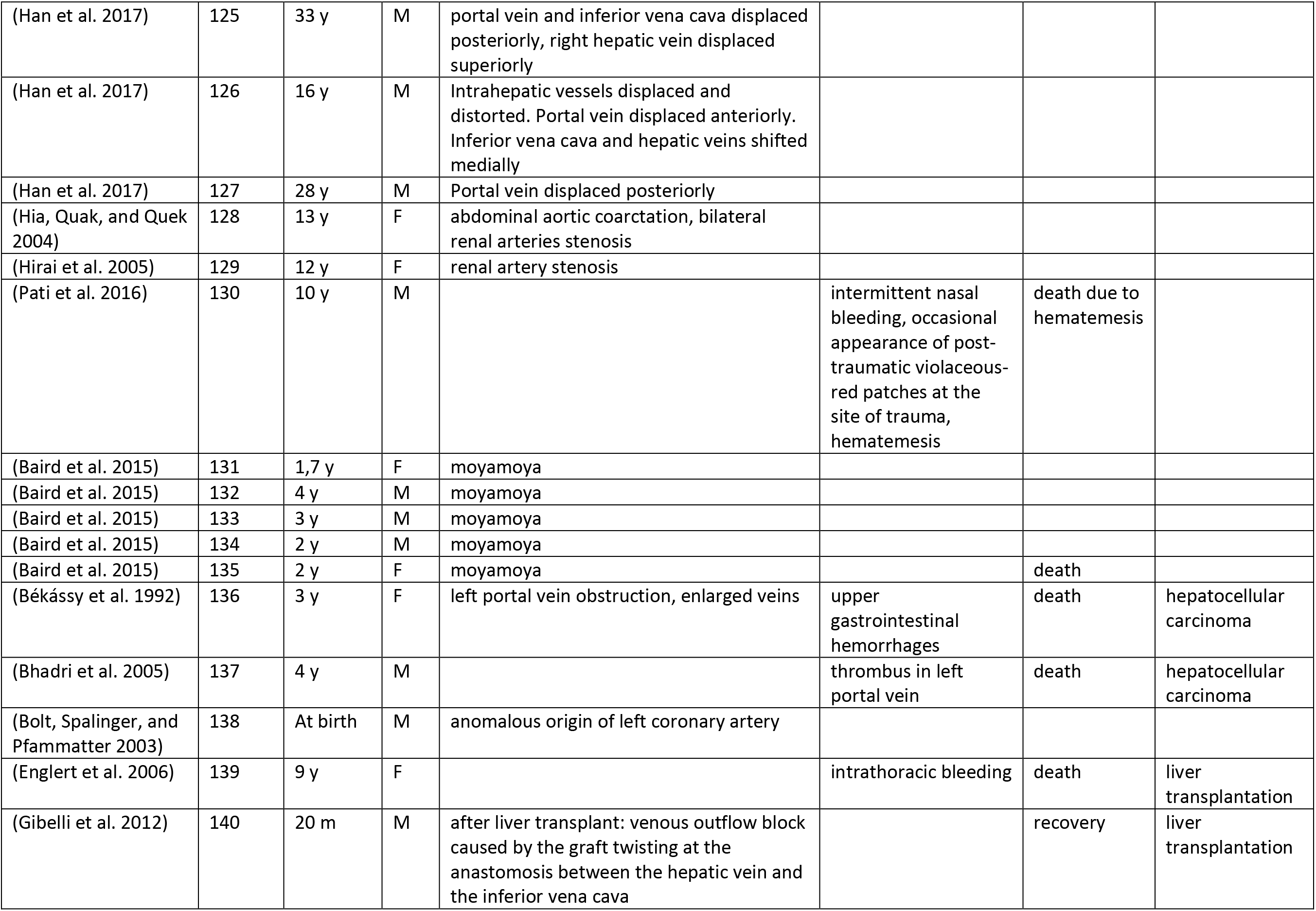
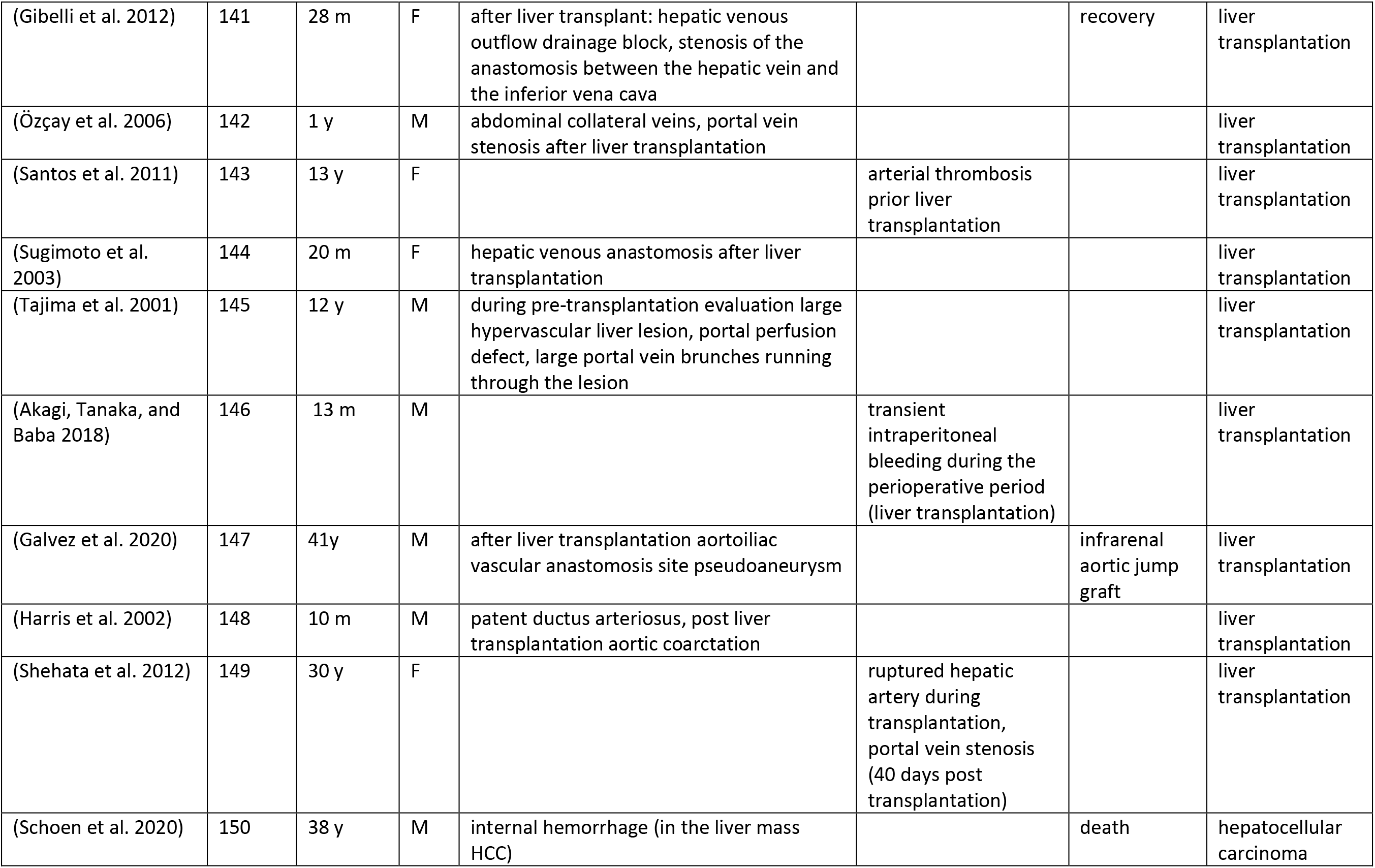
Overview of 150 patients with Alagille syndrome identified in the systematic literature review. Table describes patients’ age, sex, structural anomaly and functional event. F, female; M, male; m, month; MAPCA, major aortopulmonary collateral arteries; y, years.

| Reference | Patient # | Age at vascular event | Sex | Structural anomaly | Bleeding or Ischemic event | Outcome | Notes |
| --- | --- | --- | --- | --- | --- | --- | --- |
| (Agarwal 2005) | 1 | 2 y | M | aortic coarctation, bilateral carotid and intracerebral arteries stenosis |  |  |  |
| (Agrawal, Chennuri, and Agrawal 2015) | 2 | 5 m | F |  | intracerebral hemorrhage |  |  |
| (Castro Rodriguez, Dessy, and Demanet 2015) | 3 | at birth | F | MAPCAs, central pulmonary arteries absent |  |  |  |
| (Cheng et al. 2004) | 4 | 3 y 11 m | M | aorta stenosis |  |  | Liver transplantation |
| (Connor et al. 2002) | 5 | 13 y | M | moyamoya, bilateral carotid arteries stenosis |  |  |  |
| (DeSena, Ramaciotti, and Nugent 2012) | 6 | 7 y | F | patent ductus arteriosus |  |  |  |
| (Fitzgerald and Zuccoli 2012) | 7 | 13 d | F | bilateral internal carotid arteries agenesis |  |  |  |
| (Funk, Cohen, and Santana 2010) | 8 | 32 y | M | aorta stenosis |  |  |  |
| (Hirai et al. 2005) | 9 | 12 y | F | renal artery stenosis |  |  |  |
| (Kavukçu et al. 2005) | 10 | 16 y | F | Takayasu's arteritis, left subclavian artery stenosis |  |  |  |
| (Lemoine et al. 2010) | 11 | 7 m | F | origin of the left coronary artery from the main pulmonary artery (after heart surgery), collateral flow to the anterior descending artery through a conal branch of the right coronary artery |  |  |  |
| (Massobrio et al. 2011) | 12 | 5 y | M | Perthes disease |  |  |  |
| (Murthy et al. 2012) | 13 | 18 y | M | grade II esophageal varices |  |  |  |
| (O'Connell et al. 2012) | 14 | 17 y | F | esophageal varices | diffused subarachnoid hemorrhage, ruptured cerebellar artery aneurysm, right terminal internal carotid artery aneurysm |  | Full recovery |
| (Pavanello et al. 2015) | 15 | 7 y | F | abdominal aorta hypoplasia, basilar tip aneurysm, moyamoya, internal carotid artery stenosis |  |  |  |
| (Petaros et al. 2015) | 16 | 4 y | F | thin-walled intracranial arteries with abnormal muscular layer, lipid deposits in aorta | epidural hematoma (trauma), pancreatic hemorrhage | death |  |
| (Raas-Rothschild et al. 2002) | 17 | 14 y | M | aortic coarctation, bilateral renal artery stenosis, right subclavian artery narrow origin |  |  |  |
| (Shefler, Chan, and Ostman-Smith 1997) | 18 | 16 y | M | abdominal aorta stenosis, celiac and superior mesenteric arteries stenosis, hypoplastic hepatic artery |  |  |  |
| (Shrivastava et al. 2010) | 19 | 12 y | F | left common iliac and proximal external iliac arteries stenosis |  |  |  |
| (Shrivastava et al. 2010) | 20 | 20 y | M |  | bleeding esophageal varices | death |  |
| (Shrivastava et al. 2010) | 21 | 50 y | M | right renal, external carotid, coeliac axis and superior mesenteric arteries stenosis |  |  |  |
| (Shrivastava et al. 2010) | 22 | 62 y | M | left renal, right subclavian and coeliac arteries stenosis |  |  |  |
| (Tumialán et al. 2006) | 23 | 21 y | F | basilar terminus and ascending aorta aneurysm | diffuse subarachnoid and intraventricular hemorrhage, intraparenchymal<br>hematoma | death |  |
| (Vorstman et al. 2003) | 24 | 10 w | F |  | subdural hematoma | Full recovery |  |
| (Voykov et al. 2009) | 25 | 5 y | F | aorta coarctation |  |  |  |
| (Wauters et al. 2003) | 26 | 40 y | F | superior mesenteric artery occlusion with hypertrophy of the inferior mesenteric artery, right renal and left subclavian arteries stenosis, hypoplastic left renal artery |  |  |  |
| (Yucel, Hoorntje, and Bravenboer 2010) | 27 | 38 y | F | hypoplastic left renal artery and abdominal aorta stenosis, hypertension |  |  | Kidney transplantation |
| (Yucel, Hoorntje, and Bravenboer 2010) | 28 | - | F | renal artery stenosis |  |  |  |
| (Yucel, Hoorntje, and Bravenboer 2010) | 29 | - | M | fragile renal and abdominal arteries |  |  |  |
| (Lim et al. 2003) | 30 | 25 m | F | moyamoya, occlusion in internal carotid artery, anterior cerebral artery, occlusion of middle cerebral artery, angiostenosis | cerebral infarctions |  |  |
| (Adachi et al. 1999) | 31 | 7 y | F | aorta stenosis, esophagus varices |  |  | Liver transplantation |
| (Jacquet et al. 2007) | 32 | 31 y | F | medullary zone kidney vessels arteriosclerosis, right renal artery aneurysm |  |  |  |
| (Jacquet et al. 2007) | 33 | 30 y | F | infrarenal aortic and bilateral renal arteries stenosis |  |  |  |
| (Kayhan et al. 2010) | 34 | 43 y | M | Aorta coarctation, celiac, superior mesenteric and bilateral renal arteries stenosis, inferior mesenteric artery hypertrophy, basilar, common hepatic, superior mesenteric arteries and multiple inferior pancreaticoduodenal arcade aneurysms |  |  |  |
| (Labrecque et al. 1982) | 35 | 34 y | M | hypoplastic abdominal aorta, bilateral renal artery stenosis |  |  |  |
| (Levin et al. 1980) | 36 | 9 m | M | left superior vena cava and aberrant right subclavian artery originating from the descending aorta. The right coronary artery almost totally obliterated at the ostium |  | death |  |
| (Lorenzo-Zúñiga et al. 2003) | 37 | 16 y | F |  | bleeding esophageal varices |  |  |
| (May et al. 2013) | 38 | 3 m, 11 m | F | MAPCAs, abdominal aorta, celiac trunk and the hepatic artery stenosis, vasculopathy |  | death |  |
| (Molinero-Herguedas et al. 2008) | 39 | 17 y | M | thoracic aorta aneurysm |  |  |  |
| (Moreau et al. 1999) | 40 | 17 y | M | cavernous internal carotid artery aneurysm |  |  |  |
| (Mouzaki et al. 2010) | 41 | At birth | F | Ileal atresia, valvular aortic stenosis |  |  |  |
| (Nika et al. 2010) | 42 | At birth | M | ascending aorta and atrial septal aneurysm |  |  |  |
| (Nishikawa et al. 1987) | 43 | 15 y, 17 y | M | abdominal wall venous dilation | nasal bleeding, hematochezia, bleeding esophageal varices and liver nodular infarction, post mortem bleeding in the oral and nasal cavities and hemorrhagic ascites | Death due to hepatic failure | anemia |
| (Pascanu and Sharma 2016) | 44 | 32 y | M |  | bleeding esophageal varices |  |  |
| (Rachmel et al. 1989) | 45 | 22 m | F | patent ductus arteriosus, bilateral internal carotid arteries occlusion, moyamoya | intracranial hemorrhage |  |  |
| (Radi et al. 2011) | 46 | 38 y | F |  | Subacute ischemic lesions in the right fronto-temporal-insular region |  |  |
| (Rocha et al. 2012) | 47 | 10 y | F | patent ductus arteriosus, left renal artery stenosis, irregularities of the artery wall, moyamoya, basilar artery aneurysm |  |  |  |
| (Santamaria et al. 2016) | 48 | 21 y | F | patent ductus arteriosus, thoracic aortic coarctation, abdominal aorta hypoplasia and right renal artery stenosis |  |  |  |
| (Santamaria et al. 2016) | 49 | 24 y | F | abdominal aorta hypoplasia and right renal artery stenosis |  |  |  |
| (Santamaria et al. 2016) | 50 | 13 y | M |  | subarachnoid hemorrhage | death |  |
| (Schlosser et al. 2004) | 51 | 30 y | F | internal carotid, basilar tip, right renal, the celiac arteries aneurysms and thoracic aortic coarctation | subarachnoid hemorrhage, subdural hematoma, ruptured internal carotid artery aneurysm | full recovery |  |
| (Silberbach et al. 1994) | 52 | at birth | M | pulmonary artery collaterals |  | death |  |
| (Takenouchi et al. 2011) | 53 | 2 y | M | left middle cerebral artery aneurysm |  |  | liver transplantation |
| (Tobin et al. 2009) | 54 | 25 y | M | internal carotid artery stenosis, vasculopathy, Moyamoya |  |  |  |
| (Villa and Roenn 2014) | 55 | 21 y | M | grade III-IV esophageal varices |  |  |  |
| (Woolfenden et al. 1999) | 56 | 1 y, 21 m, 2.5y | F | supraclinoid and right internal carotid artery stenosis and reduced flow in the left anterior and middle cerebral arteries, moyamoya syndrome | embolic ischemic event affecting the left hemisphere |  |  |
| (Woolfenden et al. 1999) | 57 | 2 y | F | hypoplasia of the left pulmonary artery and its distal branches, moderate bilateral (right greater than left) distal intracranial ICA stenosis, moyamoya | four right hemispheric and one left hemispheric infarction | full recovery |  |
| (Zweier et al. 2010) | 58 | 6 m | M | systemic arterial dysplasia, persistent ductus arteriosus, filiform endings of the peripheral vessels and further more hypoplasia and multiple aortic and large arteries stenosis |  | death |  |
| (Hoffenberg et al. 1995) | 59 | 2.3 y | M |  | epidural hematoma (trauma) |  |  |
| (Hoffenberg et al. 1995) | 60 | 2 y | M |  | epidural hematoma (trauma) |  |  |
| (Hoffenberg et al. 1995) | 61 | 2 y | F | intracranial thin-walled vessels | intracerebral hemorrhage | death |  |
| (Zhang et al. 2019) | 62 | 28 y | M | narrowing of the left internal carotid artery |  |  |  |
| (Kazi, Fakhoury, and Beck 2018) | 63 | 22 m | M |  | epidural hematoma (trauma), subsequent intracranial (left epidural and subdural), infratentorial subdural, parenchymal and intraventricular bleeds |  |  |
| (Kuang et al. 2019) | 64 | 8 y | F | abdominal aorta stenosis |  |  |  |
| (Lopez et al. 2018) | 65 | 4 m | M | right hepatic artery communicating with a portal venous varix via a fistula | bleeding esophageal varices |  |  |
| (Micaglio et al. 2019) | 66 | 1 m | F | patent ductus arteriosus, MAPCAs |  |  |  |
| (Sheth, Maldonado, and Lentz 2018) | 67 | 51 y | F | patent ductus arteriosus, renal artery stenosis | lung hemorrhage |  |  |
| (Zheng et al. 2019) | 68 | 1 m | F | AVMs in liver and skin, patent ductus arteriosus, hepatic blood vessel malformations, type II Abernethy malformation |  |  |  |
| (Yokoyama et al. 2019) | 69 | 4 y | M | abdominal aorta coarctation, bilateral internal carotid, ostial, celiac, superior mesenteric and bilateral renal arteries stenosis, inferior mesenteric artery collaterals |  |  |  |
| (Shimohata et al. 2020) | 70 | 27 y | M | cephalic vein, radial, vertebral, basilar arteries stenosis, internal carotid artery aneurysm | brain infarction of the ventral midbrain, left medial thalamus, and left cerebellum (after surgery, occult blood in urine) |  |  |
| (Sanada et al. 2019) | 71 | 16.3 y, 17.1 y | F | superior mesenteric artery aneurysm, celiac and bilateral renal arteries stenosis |  |  | liver transplantation |
| (Sanada et al. 2019) | 72 | 15.5 y | M | celiac and right renal arteries stenosis |  |  | liver transplantation |
| (Sanada et al. 2019) | 73 | 22.9 y | M | common hepatic and left renal arteries stenosis |  |  | liver transplantation |
| (Sanada et al. 2019) | 74 | 11.8 y | F | celiac and left renal arteries stenosis |  |  | liver transplantation |
| (Sanada et al. 2019) | 75 | 11.5 y | M | bilateral renal arteries stenosis |  |  | liver transplantation |
| (Sanada et al. 2019) | 76 | 21.1 y,<br>21.7 y,<br>22.3 y | M | inferior pancreaticoduodenal and splenic arteries aneurysms, celiac artery stenosis |  |  | liver transplantation |
| (Sanada et al. 2019) | 77 | 1.7 y | F | celiac and bilateral renal arteries stenosis |  |  | liver transplantation |
| (Delaney et al. 2019) | 78 | 34 y | M | left internal carotid artery stenosis, Moyamoya, left common carotid artery thickened wall |  |  |  |
| (Adamson et al. 2020) | 79 |  | M |  | pulmonary hemorrhage |  |  |
| (Adamson et al. 2020) | 80 |  | M |  | pulmonary hemorrhage |  |  |
| (Adamson et al. 2020) | 81 |  | M |  | pulmonary hemorrhage |  |  |
| (Adamson et al. 2020) | 82 |  | M |  | pulmonary hemorrhage |  |  |
| (Adamson et al. 2020) | 83 |  | M |  | pulmonary hemorrhage |  |  |
| (Adamson et al. 2020) | 84 |  | M |  | pulmonary hemorrhage |  |  |
| (Adamson et al. 2020) | 85 |  | M |  | pulmonary hemorrhage |  |  |
| (Adamson et al. 2020) | 86 |  | M |  | pulmonary hemorrhage |  |  |
| (Adamson et al. 2020) | 87 |  | M |  | pulmonary hemorrhage |  |  |
| (Adamson et al. 2020) | 88 |  | F |  | pulmonary hemorrhage |  |  |
| (Adamson et al. 2020) | 89 |  | F |  | pulmonary hemorrhage |  |  |
| (Adamson et al. 2020) | 90 |  | F |  | pulmonary hemorrhage |  |  |
| (Adamson et al. 2020) | 91 |  | F |  | pulmonary hemorrhage |  |  |
| (Adamson et al. 2020) | 92 |  | F |  | pulmonary hemorrhage |  |  |
| (Carpenter et al. 2018) | 93 | 11 y | M | persistent falcine sinus |  |  |  |
| (Carpenter et al. 2018) | 94 | 3 y, 9 y | M | Moyamoya | right middle cerebral artery territory stroke, extensive ischemic injury within bilateral middle cerebral artery and left posterior cerebral artery vascular territories | death |  |
| (Carpenter et al. 2018) | 95 | 11 y | M | dolichoectasia, intracranial aneurysm |  |  |  |
| (Carpenter et al. 2018) | 96 | 13 y | F | dolichoectasia, intracranial aneurysm |  |  |  |
| (Carpenter et al. 2018) | 97 | 11y | M | persistent falcine sinus |  |  |  |
| (Carpenter et al. 2018) | 98 | 8y | M | dolichoectasia |  |  |  |
| (Carpenter et al. 2018) | 99 | 2 y | F | developmental venous anomaly |  |  |  |
| (Carpenter et al. 2018) | 100 | 8 y | M | Moyamoya |  |  |  |
| (Carpenter et al. 2018) | 101 | 6 y | M | dolichoectasia, multiple internal carotid artery aneurysms, persistent falcine sinus |  |  |  |
| (Emerick et al. 2005) | 102 | 9 y | F |  | ischemic intracranial event |  |  |
| (Emerick et al. 2005) | 103 | 9 y | F |  | subarachnoid hemorrhage after minor head trauma |  |  |
| (Emerick et al. 2005) | 104 | 15 y | F | left middle cerebral and basilar arteries aneurysm | ruptured middle cerebral artery aneurysm |  |  |
| (Emerick et al. 2005) | 105 | 11 y | M | basilary artery aneurysm |  |  |  |
| (Emerick et al. 2005) | 106 | 30 y | F | right internal carotid artery stenosis | cerebral vascular accident |  |  |
| (Emerick et al. 2005) | 107 | 4 y | M | left internal carotid artery stenosis | ischemic intracranial event |  |  |
| (Emerick et al. 2005) | 108 | 6 y | M |  | subdural hematoma after birth trauma |  |  |
| (Emerick et al. 2005) | 109 | 6 y | M |  | ischemic intracranial event after liver transplantation | death | liver transplantation |
| (Emerick et al. 2005) | 110 | 3 y | M | bilateral internal carotid artery stenosis | ischemic intracranial event |  |  |
| (Emerick et al. 2005) | 111 | 7 y | M | bilateral internal carotid, left middle cerebral and posterior cerebral arteries stenosis | ischemic intracranial event |  |  |
| (Emerick et al. 2005) | 112 | 15 y | M | left internal carotid, anterior cerebral, right middle cerebral arteries stenosis |  |  |  |
| (Emerick et al. 2005) | 113 | 5 y | M | absence of left internal carotid artery, beading of the right posterior and middle cerebral arteries, anomalous artery from basilar to left middle cerebral artery |  | death |  |
| (Emerick et al. 2005) | 114 | 2 y | M | right internal carotid and middle cerebral arteries stenosis |  |  |  |
| (Emerick et al. 2005) | 115 | 1.7 y | F | bilateral internal carotid and middle cerebral arteries stenosis, two aneurysm of supraclinoid internal carotid artery, moyamoya | ischemic intracranial event |  |  |
| (Cowan et al. 2004) | 116 | 18 y, 23 y | M | aortic coarctation | ruptured basilar artery aneurysm; posterior communicating artery infundibulum aneurysm | full recovery |  |
| (DesRoches et al. 2013) | 117 | 28 y | F |  | gastrointestinal bleeds, multiple splenic hemangiomas |  |  |
| (Doberentz, Kuchelmeister, and Madea 2015) | 118 | 25 y | F | lung, heart, intracranial thin-walled arteries | subarachnoid hemorrhage, large saccular ruptured basilar artery aneurysm | death |  |
| (Dzumhur and Goss 1995) | 119 | 2 w | F |  | deep vein thrombosis with swelling and cyanosis of right leg, absent venous flow from the common femoral vein | recovery |  |
| (Esther, Francesca, and Jose 2017) | 120 | At birth | F | bilateral renal arterial stenosis | disseminated intravascular coagulation state |  | renal failure |
| (Ferreira-Magalhaes 2014) | 121 | 7 y | F | bilateral renal, superior mesenteric arteries stenosis, celiac trunk kinking, left renal artery pseudoaneurysm | left perirenal hematoma with infarction of the superior half of left kidney |  |  |
| (Fiorda-Diaz et al. 2017) | 122 | 17 y | f |  | subarachnoid hemorrhage with blood within third and fourth ventricles, and within the pre-pontine cistern, intracranial epidural hematoma |  |  |
| (Frongillo et al. 2015) | 123 | 28 y | F |  | cerebral and ocular hemorrhage | hepatic failure with hepatorenal syndrome | coagulopathy, liver transplantation |
| (Gaba et al. 2008) | 124 | 28 y | F | moyamoya | subarachnoid hemorrhage, ruptured vertebrobasilar junction aneurysm |  | full recovery |
| (Han et al. 2017) | 125 | 33 y | M | portal vein and inferior vena cava displaced posteriorly, right hepatic vein displaced superiorly |  |  |  |
| (Han et al. 2017) | 126 | 16 y | M | Intrahepatic vessels displaced and distorted. Portal vein displaced anteriorly. Inferior vena cava and hepatic veins shifted medially |  |  |  |
| (Han et al. 2017) | 127 | 28 y | M | Portal vein displaced posteriorly |  |  |  |
| (Hia, Quak, and Quek 2004) | 128 | 13 y | F | abdominal aortic coarctation, bilateral renal arteries stenosis |  |  |  |
| (Hirai et al. 2005) | 129 | 12 y | F | renal artery stenosis |  |  |  |
| (Pati et al. 2016) | 130 | 10 y | M |  | intermittent nasal bleeding, occasional appearance of post-traumatic violaceous-red patches at the site of trauma, hematemesis | death due to hematemesis |  |
| (Baird et al. 2015) | 131 | 1,7 y | F | moyamoya |  |  |  |
| (Baird et al. 2015) | 132 | 4 y | M | moyamoya |  |  |  |
| (Baird et al. 2015) | 133 | 3 y | M | moyamoya |  |  |  |
| (Baird et al. 2015) | 134 | 2 y | M | moyamoya |  |  |  |
| (Baird et al. 2015) | 135 | 2 y | F | moyamoya |  | death |  |
| (Békássy et al. 1992) | 136 | 3 y | F | left portal vein obstruction, enlarged veins | upper gastrointestinal hemorrhages | death | hepatocellular carcinoma |
| (Bhadri et al. 2005) | 137 | 4 y | M |  | thrombus in left portal vein | death | hepatocellular carcinoma |
| (Bolt, Spalinger, and Pfammatter 2003) | 138 | At birth | M | anomalous origin of left coronary artery |  |  |  |
| (Englert et al. 2006) | 139 | 9 y | F |  | intrathoracic bleeding | death | liver transplantation |
| (Gibelli et al. 2012) | 140 | 20 m | M | after liver transplant: venous outflow block caused by the graft twisting at the anastomosis between the hepatic vein and the inferior vena cava |  | recovery | liver transplantation |
| (Gibelli et al. 2012) | 141 | 28 m | F | after liver transplant: hepatic venous outflow drainage block, stenosis of the anastomosis between the hepatic vein and the inferior vena cava |  | recovery | liver transplantation |
| (Özçay et al. 2006) | 142 | 1 y | M | abdominal collateral veins, portal vein stenosis after liver transplantation |  |  | liver transplantation |
| (Santos et al. 2011) | 143 | 13 y | F |  | arterial thrombosis prior liver transplantation |  | liver transplantation |
| (Sugimoto et al. 2003) | 144 | 20 m | F | hepatic venous anastomosis after liver transplantation |  |  | liver transplantation |
| (Tajima et al. 2001) | 145 | 12 y | M | during pre-transplantation evaluation large hypervascular liver lesion, portal perfusion defect, large portal vein branches running through the lesion |  |  | liver transplantation |
| (Akagi, Tanaka, and Baba 2018) | 146 | 13 m | M |  | transient intraperitoneal bleeding during the perioperative period (liver transplantation) |  | liver transplantation |
| (Galvez et al. 2020) | 147 | 41y | M | after liver transplantation aortoiliac vascular anastomosis site pseudoaneurysm |  | infrarenal aortic jump graft | liver transplantation |
| (Harris et al. 2002) | 148 | 10 m | M | patent ductus arteriosus, post liver transplantation aortic coarctation |  |  | liver transplantation |
| (Shehata et al. 2012) | 149 | 30 y | F |  | ruptured hepatic artery during transplantation, portal vein stenosis (40 days post transplantation) |  | liver transplantation |
| (Schoen et al. 2020) | 150 | 38 y | M | internal hemorrhage (in the liver mass HCC) |  | death | hepatocellular carcinoma |

